# Behavioral control through the direct, focal silencing of neuronal activity

**DOI:** 10.1101/2023.05.31.543113

**Authors:** Anna V. Elleman, Nikola Milicic, Damian J. Williams, Christine J. Liu, Allison L. Haynes, Jane Simko, David E. Ehrlich, Christopher D. Makinson, J. Du Bois

## Abstract

Voltage-gated sodium channel (Na_V_) activity underlies electrical signaling, synaptic release, circuit function, and, ultimately, behavior. Molecular tools that enable precise control of Na_V_ subpopulations make possible temporal regulation of neuronal activity and cellular communication. To rapidly modulate Na_V_ currents, we have rendered a potent Na_V_ inhibitor, saxitoxin, transiently inert through chemical protection with a novel nitrobenzyl-derived photocleavable group. Light-induced uncaging of the photocaged toxin, STX-bpc, effects rapid inhibitor release and focal Na_V_ block. We demonstrate the efficacy of this reagent for manipulating action potentials in mammalian neurons and brain slice and for altering locomotor behavior in larval zebrafish. Photo-uncaging of STX-bpc is a non-invasive, effective method for reversible, spatiotemporally precise tuning of Na_V_ currents, application of which requires no genetic manipulation of the biological sample.

## INTRODUCTION

A desire to understand how information is processed in the nervous system has motivated the development of methods for real-time, spatiotemporal manipulation of neuronal activity. An array of technologies^1, 2^ now exist for controlling and characterizing the function of different brain regions, circuits, and cell types; none, however, act directly on voltage-gated sodium (Na_V_) channels, the proteins responsible for action potential (AP) generation and propagation. Instead, methods like optogenetics^3, 4^ rely on the expression of exogenous ion channels that constructively (e.g, ChR2)^5^ or destructively (e.g., WiChR)^6^ interfere with endogenous currents to produce a net change in electrical output. While optogenetic methods have significantly improved our ability to control cellular activity and regulate neural circuits^7^, such technologies are often limited by toxicity,^8^ delivery,^9^ and expression^10^ of the requisite ectopic proteins, as well as the altered capacitance,^11^ electrochemical gradients, ^12^ and pH ^13^ of the target cells. ^14^ We have developed an alternative tool for manipulating APs both *in vitro* and *in vivo* that avoids genetic manipulation. Herein, we describe the design, optimization, and validation of a broadly applicable small molecule reagent (STX-bpc) for precise, light-mediated inhibition of Na_V_s in neurons.

The control of endogenous sodium currents requires a modulator with high efficacy and specificity towards neuronal Na_V_ isoforms. Saxitoxin is a potent inhibitor of Na_V_s that acts by binding to the outer mouth of the channel pore, just above the selectivity filter, to occlude ion passage into cells.^15, 16^ Characteristics that favor STX as opposed to other Na_V_ inhibitors^17^ for Na_V_ targeting include: (1) its high potency against all channel conformational states (open, closed, inactive) such that binding occurs independently of stimulus; (2) its fast rate of association (k_on_ ∼ 10^6^ M^−1^ s^−1^),^18^ enabling rapid block; and (3) its extracellular binding site, which allows for complete, reversible inhibition of channels. Additionally, saxitoxin is potent against most vertebrate^19^ and insect^20, 21^ neuronal and cardiovascular Na_V_s. In mammals, STX has nanomolar affinity for seven of nine Na_V_ subtypes (Na_V_1.1–1.7), including all Na_V_s expressed in the central nervous system.^17, 22^ We therefore anticipated that modification of STX with a destabilizing, photocleavable protecting group would yield a small molecule tool that is completely inert upon application to cells and tissue and rapidly uncaged upon exposure to light.

In this report, we demonstrate the utility of a photocaged saxitoxin, STX-bpc, for both tuning electrical excitability and achieving complete AP block with focal precision, in dissociated rat neurons, in mouse cortical brain slice, and in larval zebrafish. We further illustrate the practicality of this technology for the targeted manipulation of locomotor behaviors. Given its efficacy across cell, tissue, and live organism absent any genetic manipulation, STX-bpc is a valuable complement to available optogenetic methods for spatiotemporal modulation of electrical signaling.

## RESULTS

### Design of Photocaged Saxitoxin

We previously described the development of coumarin-caged saxitoxins (e.g., STX-eac), providing proof-of-principle that brief exposure of these compounds to light can block AP firing and slow the propagation of compound action potentials along axon fiber tracks.^23^ The utility of STX-eac for neural control, however, is limited by: (1) a small concentration operating window (the result of a ∼20-fold change in IC_50_ between caged and uncaged toxin) that limits the degree to which Na_V_ block can be fine-tuned; and (2) the need to deliver high-powered UV light to promote photocleavage, raising concerns of phototoxicity and tissue heating, particularly in cases of prolonged neuronal silencing, when long-duration exposure is required. Additional optimization of this reagent was hampered by the coumarin protecting group, as derivatization is limited and can reduce the efficiency of photocleavage.^23, 24^ We thus pursued alternative photocage designs.

To improve the potency differential between caged and uncaged STX, we identified a photocleavable protecting group, [methyl-2-nitropiperonyl-oxy]carbonyl (MeNPOC, **Figure 1A**),^25, 26^ amenable to chemical modifications including increase of its steric size and introduction of anionic units to destabilize toxin binding to the electronegative Na_V_ pore.^15, 16^ Replacement of the methyl moiety in MeNPOC with an electron-deficient group was expected to substantially improve uncaging efficiency, as demonstrated by the ∼100-fold increase in quantum yield for the analogous CF_3_-modified cage.^27, 28^ To maximize uncaging efficiency while minimizing the affinity of the photocaged STX for Na_V_, we replaced the Me-substituent in MeNPOC with a sterically large and/or negatively charged aryl amide, aryl ester, or aryl carboxylate group, thus affording a small collection of electron-deficient nitrobenzyl photocages [Hammett σ-constants(para): 0.36, CONHMe; 0.45, COOMe; 0.45, COOH; vs. 0.54, CF_3_].^29^

**Fig. 1:**
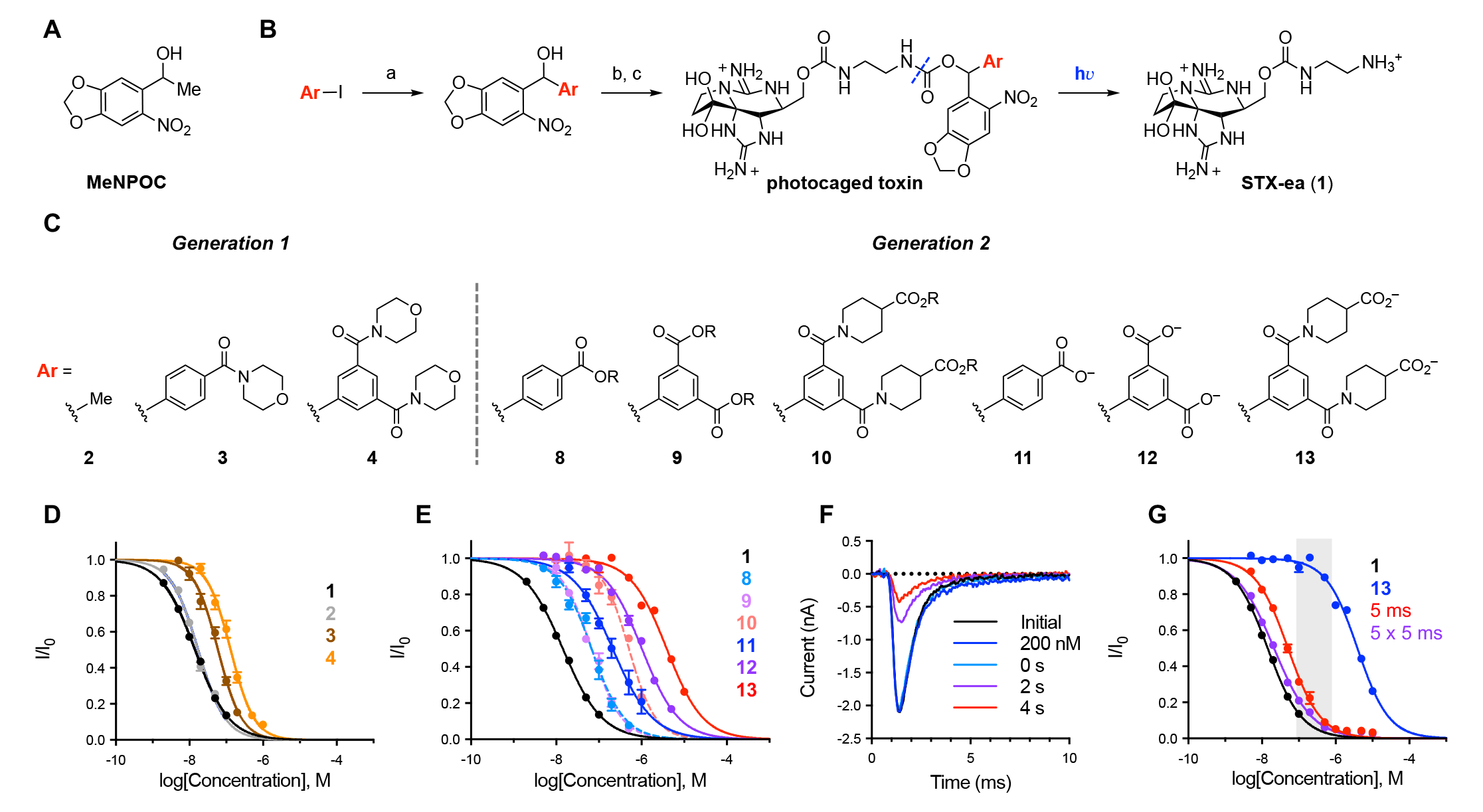
Optimized photocaged toxin enables precise control of rNa_V_1.2 current. (**A**) MeNPOC (3,4-(methylenedioxy)-6-nitrophenylethoxycarbonyl) photo-protecting group. (**B**) Scheme depicting generalized synthesis and uncaging of photocaged STXs. a, *i*-PrMgCl•LiCl then 6-nitropiperonal (Ar = aryl); b, *N,N’*-disuccinimidyl carbonate, Et_3_N; c, saxitoxin-ethylamine **1**; details are available in the Extended Data. Purple dashed line depicts location of photocleavage. (**C**) Generation 1 and 2 photocaged STXs. R = allyl. (**D**) Electrophysiological characterization of Generation 1 photocaged STXs against Na_V_1.2 CHO. IC_50_s, Hill coefficients: **1** = 14.4 ± 0.3 nM, –0.94 ± 0.02 (n = 6–7); **2** = 17.3 ± 0.7 nM, –1.17 ± 0.05 (n = 6–7); **3** = 59.8 ± 3.3 nM, –1.35 ± 0.10 (n = 3); **4** = 132.1 ± 7.3 nM, –1.35 ± 0.10 (n = 6). (**E**) Electrophysiological characterization of Generation 2 photocaged STXs against Na_V_1.2 CHO. IC_50_s, Hill Coefficients: **1** = 14.4 ± 0.3 nM, –0.94 ± 0.02 (n = 6–7); **8** = 67.6 ± 4.9 nM, –1.13 ± 0.09 (n = 4); **9** = 67.6 ± 3.6 nM, –1.23 ± 0.08 (n = 4–5); **10** = 507.2 ± 39.0 nM, –1.38 ± 0.15 (n = 3–4); **11** = 211.3 ± 25.7 nM, –0.93 ± 0.12 (n = 5–9); **12** = 1024.9 ± 38.6 nM, –1.01 ± 0.04 (n = 4–5); **13** = 3919.4 ± 172.6 nM, –1.02 ± 0.05 (n = 5–7). (**F**) Uncaging of 200 nM **13** against Na_V_1.2 CHO. Traces collected in the order: Initial, 200 nM **13**, 0 s, 2 s, 4 s. Laser applied immediately prior to 0 s trace. Currents observed at 4 s used to calculate uncaging data described in (**G**). (**G**) Electrophysiological characterization and laser uncaging of **13** against Na_V_1.2 CHO. Apparent IC_50_s, Hill Coefficients: **1** = 14.4 ± 0.3 nM, –0.94 ± 0.02 (n = 6–7); **13** = 3919.4 ± 172.6 nM, –1.02 ± 0.05 (n = 5–7); **13** uncaged with 5 ms laser = 51.0 ± 1.9 nM, –0.98 ± 0.03 (n = 5–7); **13** uncaged with 5 x 5 ms laser = 21.3 ± 0.7 nM, –0.84 ± 0.02 (n = 5–7). Potency window highlighted in grey.

### Synthesis and Optimization of Photocaged STXs

Caged compounds were readily prepared by 1,2-addition of the appropriate aryl Grignard reagent to 6- nitropiperonal and subsequent conjugation of the corresponding N-hydroxysuccinimide ester to STX ethylamine **1**^30, 31^ (**Figure 1B**). [See extended data for full synthesis and characterization.] Photocaged STXs were then screened for potency against Na_V_1.2 stably expressed in CHO cells with uncaging elicited by 5 ms pulses of 130 mW, 355 nm light. For clarity of discussion, caged STXs have been divided into two categories: Generation 1, comprising the ‘base model’ STX MeNPOC, **2**, as well as amide-modified photocaged derivatives of varying steric bulk and electronic substitution (**3**–**7**); and Generation 2, composed of carboxylic acid-modified (i.e., anionic) structures (**11**, **12**, **13**) and their uncharged, allyl ester counterparts (**8**, **9**, **10**) (**Figure 1C, Extended Data Figure 1**).

Examination of Generation 1 photocaged STXs revealed that steric substitution alone was insufficient to effectively destabilize binding. The simplest construct, STX MeNPOC **2**, was similarly potent to the uncaged inhibitor **1** (IC_50_: 17.3 nM vs. 14.4 nM; **Figure 1D, Extended Data Figure 2A**) despite a seventy percent increase in molecular mass (582 vs. 301 Da). While other sterically larger derivatives diminished the potency of the caged toxin (IC_50_: **4** > **3** > **2**, 132.1 nM > 58.9 nM > 17.3 nM; **Figure 1D, Extended Data Figure 2A–C**), at best, only a nine-fold increase in the IC_50_ value compared to **1** could be obtained (IC_50_: **4** ∼ **5** ∼ **6** ∼ **7**, 132.1 nM ∼ 114.2 nM ∼ 128.1 nM ∼ 121.9 nM; **Figure 1D, Extended Data Figure 2C–F**).

**Fig. 2:**
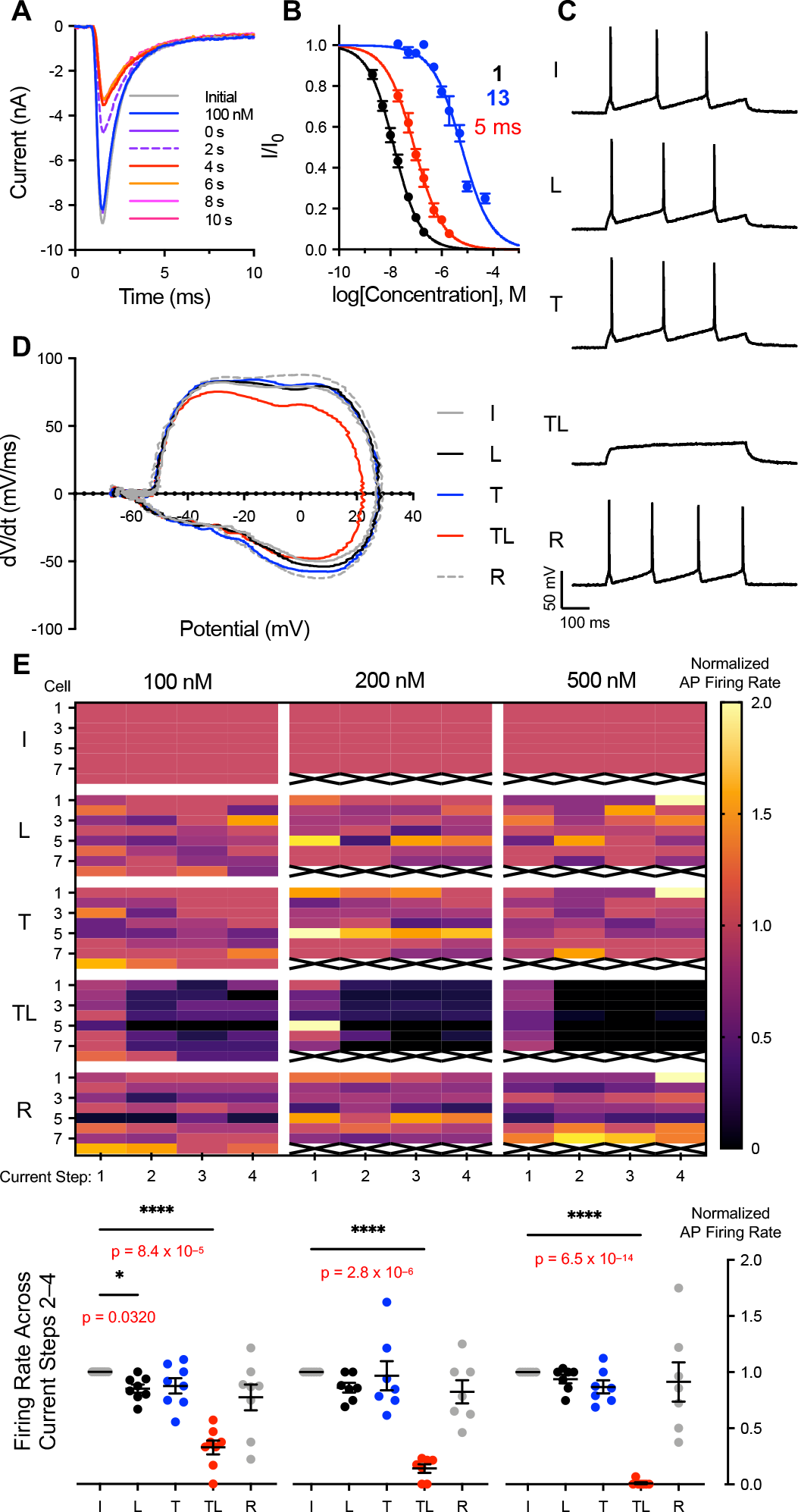
Uncaging of **13** rapidly blocks action potential propagation in dissociated embryonic hippocampal neurons. (**A**) Uncaging of 100 nM **13**. Traces collected in the order: Initial, 100 nM **13**, 0 s, 2 s, etc. Laser applied immediately prior to 0 s trace. Currents observed at 10 s used to calculate uncaging data described in (B). (**B**) Electrophysiological characterization of **13**. Apparent IC_50_s, Hill coefficients: **1** = 14.1 ± 0.8 nM, –0.86 ± 0.04 (n = 4–5); **13** = 5210.0 ± 409.1 nM, –0.88 ± 0.06 (n = 5–6); **13** uncaged with 5 ms laser = 87.6 ± 6.8 nM, –0.78 ± 0.05 (n = 5). (**C**) Representative traces depicting initial (I), laser applied (L), 200 nM **13** applied (T), 200 nM **13** and laser applied (TL), and recovered (R) after wash-off AP trains evoked by 500 ms, 50–150 pA current injections. Data taken from replicate current step 2 vis-à-vis (**E**). (**D**) Representative phase plot depicting application and uncaging of 100 nM **13**. Calculated from first action potential in current step 2 vis-à-vis (**E**). (**E**) Heat map summary of AP firing rate after application and uncaging of three different concentrations of **13** (100 nM, 200 nM, 500 nM) color-coded by number of action potentials per step (four replicate current steps at 0.25 Hz per condition). Equilibrated normalized action potential firing rate (i.e., over current steps 2–4) compared below (n = 7–8,*p<0.05, ****p<0.0001, one-way ANOVA with Tukey’s correction).

Among Generation 2 photocaged STXs, we found carboxylate-substituted structures (**11**, **12**, **13**) to be fifteen-fold less potent than analogous allyl esters (**8**, **9**, **10**, respectively), thus establishing the importance of anionic charge incorporation to destabilize binding of the caged compound (**Figure 1E, Extended Data Figure 2**). Steric bulk also decreased affinity of the caged inhibitor for Na_V_s, as bis-piperidine carboxylate **13** (STX-bpc) is approximately four-fold less potent than bis-carboxylate **12** (IC_50_: **13**, 3.9 µM vs. **12**, 1.0 µM; also compare compounds **10**, 0.5 µM vs. **9**, 0.07 µM). STX-bpc was ultimately selected as the optimal probe for subsequent validation studies (*vide infra*).

All anionic photocaged STXs are photo-deprotected with unexpectedly high efficiency, particularly compared to the equivalent allyl ester derivatives (**Figure 1F, 1G, Extended Data Figure 2**). Upon application of 5 x 5 ms pulses of 355 nm light, the apparent potency of esters **8**–**10** decreased only minimally (e.g., IC_50_ ratio caged vs. uncaged: **8**, 1.5; **9**, 1.6; **10**, 2.4; **Extended Data Table 2**). In contrast, carboxylates **11**–**13** uncage almost completely under the same protocol, with post-exposure potencies approaching that of the parent **1** (IC_50_ following 5 x 5 ms pulses of 355 nm light: **11**, 27.1 nM; **12**, 25.1 nM; **13**, 21.3 nM vs. IC_50_ **1**, 14.4 nM). Carboxylate incorporation thus dramatically improves uncaging efficiency, as photo-release of the ‘base model’ photocaged toxin **2** does not occur under our conditions (**Extended Data Figure 2A**, quantum yield for this protecting group is estimated at ϕ = 0.0075^32^). Why carboxylate substitution has this effect is unclear, as the absorbance spectra of photocaged STXs **11**–**13** are unchanged from that of **2** (**Extended Data Figure 3**).

**Fig. 3:**
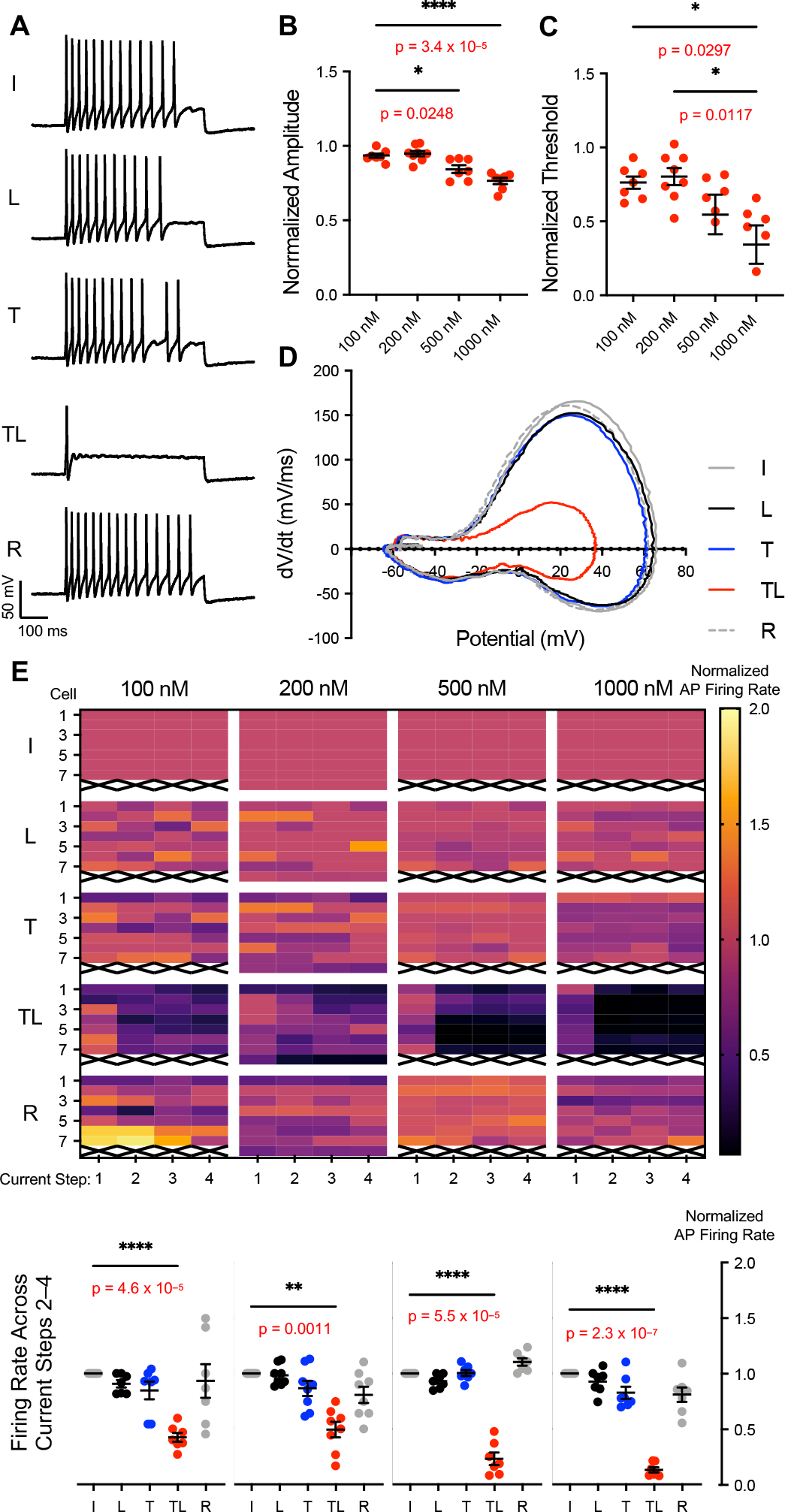
Uncaging of **13** effects dose-dependent changes in action potential shape and propagation in dissociated embryonic dorsal root ganglia cells. (**A**) Representative traces depicting initial (I), laser applied (L), 500 nM **13** applied (T), 500 nM **13** and laser applied (TL), and recovered (R) after wash-off AP trains evoked by 500 ms, 250–550 pA current injections. Data taken from replicate current step 2 vis-à-vis (E). (**B**) Dose-dependent reduction in AP amplitude after application and uncaging of **13** at listed concentrations. Data calculated from first action potential in current step 2 vis-à-vis (E). Unlisted significant p-values: 200 nM vs 500 nM, p = 0.0064; 500 nM vs 1000 nM, 0.0628 (n = 6–8). (**C**) Dose-dependent reduction in AP threshold after application and uncaging of **13** at listed concentrations (n = 6–8). Data calculated from first action potential in current step 2 vis-à-vis (**E**). (**D**) Representative phase plot depicting application and uncaging of 1000 nM **13**. Calculated from first action potential in current step 2 vis-à-vis (**E**). (**E**) Heat map summary of AP firing rate after application and uncaging of four different concentrations of **13** (100 nM, 200 nM, 500 nM, 1000 nM) color-coded by number of action potentials per step (four replicate current steps at 0.25 Hz per condition). Equilibrated normalized action potential firing rate (i.e., over current steps 2–4) compared below (n = 7–8). (*p<0.05, **p<0.01, ****p<0.0001, one-way ANOVA with Tukey’s correction).

STX-bpc **13** is over 270-fold less potent than the parent compound **1** (IC_50_ = 3.9 µM vs. 14.4 nM; **Figure 1E**) and uncages rapidly upon application of 355 nm light (**Figure 1F**). The large concentration window over which this caged toxin may be employed (**Figure 1G**, grey) enables tuning of Na_V_ current amplitude, as [STX-bpc] ≤ 500 nM blocks ≤ 10% of channels prior to uncaging, but as many as 90% following deprotection. As the concentration of STX-bpc can be varied, so too can the time constant (***τ***) for Na_V_ block. At 200 nM STX-bpc, toxin release and inhibition of channels occurs with a ***τ*** = 1.0 ± 0.1 seconds (**Extended Data Figure 4**); at 500 nM **13**, this value decreases (***τ*** = 0.7 ± 0.07 seconds). Focal uncaging of STX-bpc thus precisely modulates Na_V_ current, allowing the speed and degree of Na_V_ block to be fine-tuned through changes in STX-bpc concentration, light intensity, and exposure duration.

**Fig. 4:**
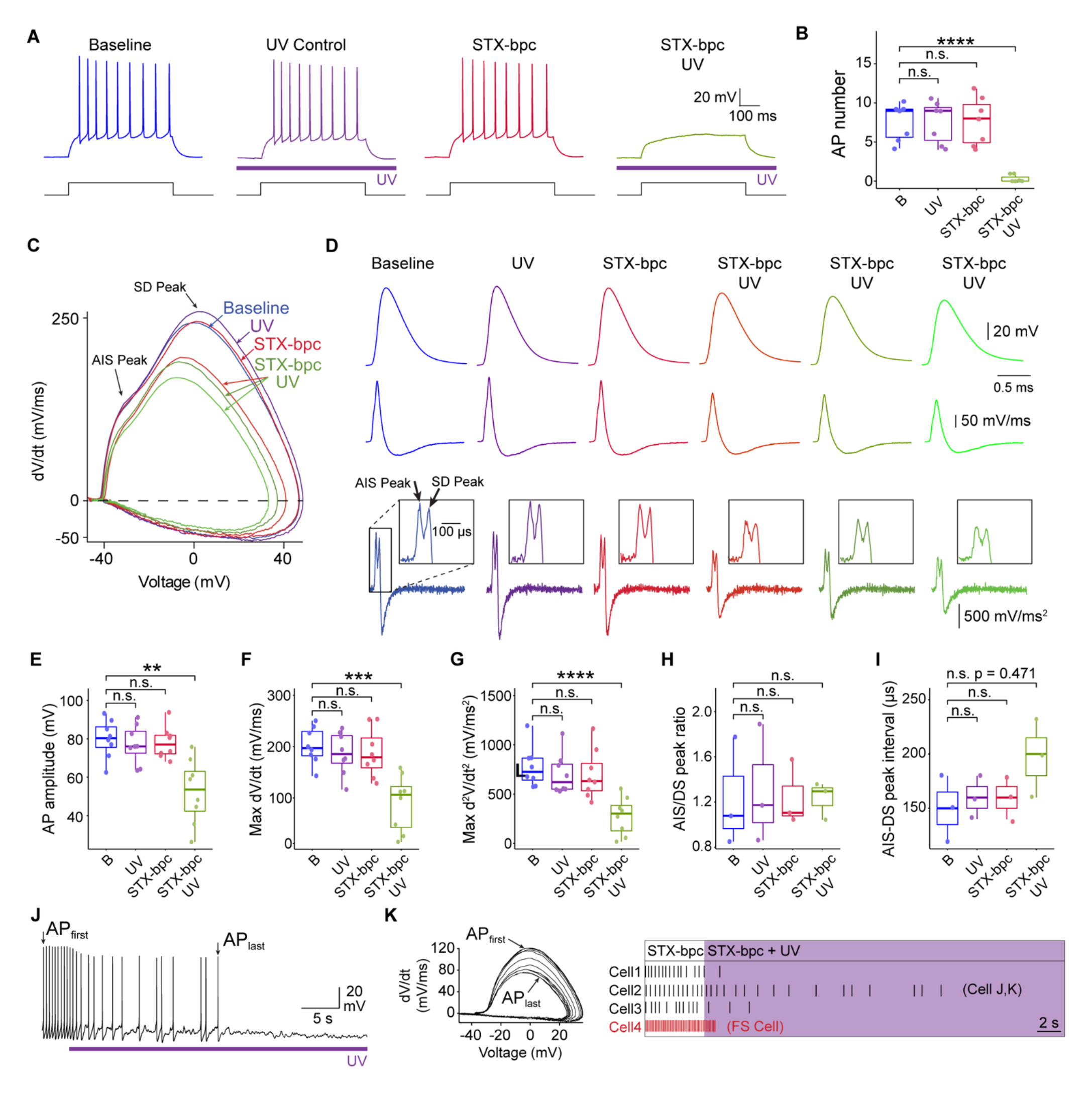
Activation of STX-bpc **13** abolishes action potentials in layer 4 cortical neurons. (**A**) Electrophysiological response of layer 4 cortical neurons (S1bf) to 500 ms 50–150 pA current steps. APs were present during a baseline period shown in blue when exposed to 365 nm LED UV light alone (purple) and in the presence of 250 nM STX-bpc **13** (red), but were blocked following exposure to both UV and **13** (green). The same color scheme is used throughout the figure. (**B**) Box plot showing the number of APs evoked under the conditions shown in (**A**). Each point represents a different cell. Dunnett’s test \*\*\**P* < 0.001. (**C**) Phase plane plot derived from the first evoked AP. Experimental conditions are indicated by the colored arrows. The axon initial segment (AIS) and somatodendritic (SD) peaks are shown by the black arrows. (**D**) AP traces (upper trace), and first (middle trace) and second (lower trace) derivative plots of the same APs in (**C**). The inset on the second derivative plots shows an enlarged portion of the same plot in which the AIS and SD peaks are clearly discernable (**E–I**). Box plots showing quantification of AP characteristics. The peak ratio (**H**) and peak amplitude (**I**) could only be plotted where both SD and AIS peaks were distinguishable (n = 3). Dunnett’s test \*\*\**P* < 0.001,** *P* < 0.01 (**J**) Continuous recording shows APs recorded from a neuron in the presence of **13**. UV exposure is indicated by a purple line. (**K**) Phase plane plot of the recording shown in (**J**). The portions of the plots representing the first and last APs are indicated by the arrows. (**L**) Raster plot showing action potential firing over time in the presence of **13** before and during UV light exposure (purple). Four neurons including one fast spiking neuron (red) are displayed. Each AP is represented by a vertical bar and time is shown on the horizontal axis.

### Na_V_ Block in Rat Dissociated Hippocampal Neurons

The advantageous properties of STX-bpc against Na_V_1.2 (CHO cells) are evident in experiments with embryonic rat hippocampal neurons. In electrophysiology recordings with dissociated neuronal cells, STX-bpc has an IC_50_ = 5.2 µM, 370-fold greater than that of **1** (14.1 nM; **Figure 2A, 2B**).^23^ Uncaging proceeds rapidly (***τ*** = 1.6 seconds at 200 nM STX-bpc) and channel block extends for tens of seconds prior to wash-off (**Figure 2A**, **Extended Data Figure 5**). A single 5 ms laser pulse releases sufficient concentrations of **1** to give an apparent IC_50_ = 87.6 nM. Accordingly, Na_V_ currents in these cells can be modulated across a large range (e.g., 25% block of peak current following uncaging of 20 nM STX-bpc; 81% block of peak current upon uncaging 500 nM STX-bpc).

**Fig. 5:**
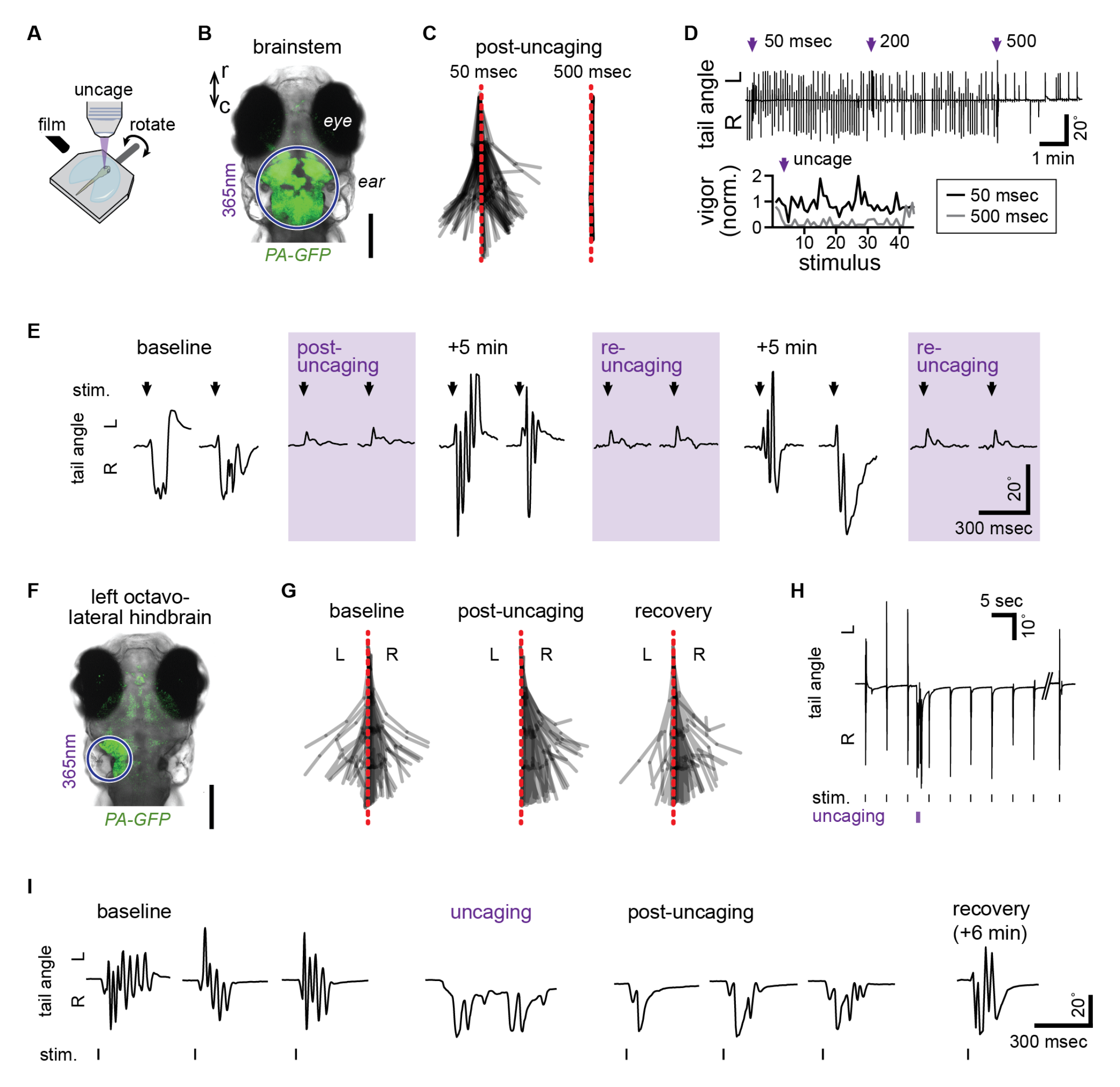
Uncaging of STX-bpc **13** *in vivo* manipulates larval zebrafish behavior. (**A**) Schematic for swim tracking following rotation stimulus presentation and focal uncaging. (**B**) Confocal photomicrograph of dorsal perspective of a larval zebrafish head expressing photoactivatable GFP (PA-GFP), showing the region of UV light exposure (*purple circle*) during brainstem-wide uncaging. Scale bar = 200 µm. (**C**) Superimposed tail segment positions from a tracked larva across 3 stimulus presentations following 50 msec (*left*) or 500 msec (*right*) uncaging in the brainstem. (**D**) Tail angle (*top*) of a larva throughout a train of stimulus presentations every 5 sec, interspersed with brainstem uncaging of increasing duration (*purple arrow*). Swimming vigor (*bottom*) is plotted as a function of stimulus number for 3 stimuli before and 42 after uncaging for 50 and 500 msec. (**E**) Tail angle during the first 2 stimulus presentations (arrow) of series without uncaging or immediately following uncaging (purple shading, *post-uncaging* or *re-uncaging*). (**F**) Photomicrograph, as in (**B**), showing the location of activated PA-GFP following focal uncaging in the left octavolateral hindbrain to unilaterally inactivate sensory areas for rotation stimuli. Scale bar = 200 µm. (**G**) Superimposed tail positions from a tracked larva across 3 stimulus presentations at baseline, immediately following uncaging in the left octavolateral hindbrain (*post-uncaging*), and 6 min after uncaging (*recovery*). (**H**,**I**) Tail angle of the larva in (**G**) in response to rotation stimulus presentation before and after uncaging (**H**,**I**) and following recovery (**I**).

Because uncaging proceeds efficiently, AP trains in neurons are abrogated using low concentrations of STX-bpc (e.g., 200 nM, **Figure 2C**). Application of 100–500 nM STX-bpc to hippocampal cells gives no observable changes in action potential shapes or firing frequencies (**Figure 2E**, **Extended Data Figure 6**). Photolysis of STX-bpc, however, decreases AP firing rate in a dose-dependent manner (remaining percent of initial firing rate: 100 nM, 33%; 200 nM, 14%; 500 nM, < 1%) with uncaging of 500 nM STX-bpc (i.e., 81% peak Na^+^ current inhibition) blocking all APs (**Figure 2E**). Uncaging 100–200 nM STX-bpc also alters action potential shape (**Figure 2C, Extended Data Figure 7**), with uncaging of 200 nM STX-bpc reducing AP amplitude by 17% (p = 0.02) and shifting firing threshold to more positive potentials by approximately 12 mV (p = 0.14) (**Extended Data Figure 7B, 7C**), as expected given the decrease in the number of functional Na_V_s.^33^ The characteristic kink at threshold voltage in the AP phase plot, ascribed to cooperativity among Na_V_s^34^ or AP initiation at the distal axon initial segment,^35^ disappears upon uncaging STX-bpc, also consistent with a reduction in Na_V_ density (**Extended Data Figure 7A, 7D**). Parameters describing the velocity and acceleration of AP rise similarly decrease upon uncaging STX-bpc, with maximal rate of rise dropping by 54% (p = 0.03) and acceleration by 58% (p = 0.002; **Extended Data Figure 7E, 7G**). AP firing recovers fully upon toxin wash-off at all concentrations (**Figure 2C, Extended Data Figure 7**). Collectively, these data demonstrate that photo-deprotection of STX-bpc can be used to precisely tune APs in hippocampal neurons through the rapid and selective block of Na_V_s.

### Electrical Silencing in Rat Dissociated Dorsal Root Ganglia

As with hippocampal neurons, STX-bpc displays dose-dependent effects on AP firing frequency (**Figure 3A, 3E**) and shape (**Figure 3B–D, Extended Data Figure 8–9**) in embryonic rat dorsal root ganglia (DRG) cells. These effects are fully resolved upon toxin wash-off. Application and uncaging of 200 nM STX-bpc reduces AP firing rate to 49% of the initial value; with 500 and 1000 nM STX-bpc, these values fall to 23% 13%, respectively (**Figure 3E**). Photo-deprotecting 1000 nM STX-bpc almost completely blocks AP trains (6/7 cells); the first action potential spike, however, is always recorded. This finding is in contrast to STX-bpc uncaging in hippocampal neurons, which completely shuts down the generation of single action potentials at 500 nM.

In embryonic DRG neurons, every phase of AP shape is altered by STX-bpc uncaging at concentrations ≥500 nM (**Figure 3D, Extended Data Figure 8**). Uncaging of 500 nM STX-bpc decreases AP amplitude by 15% (p = 0.005) and shifts threshold to more positive potentials by 9 mV (p = 0.02; **Extended Data Figure 8B– C**). The magnitude of the maximum rate of AP rise is also diminished by 49% (p = 0.0002) and AP fall by 28% (p = 0.05; **Extended Data Figure 8E–F**). Maximal AP acceleration is similarly reduced by 66% (p = 0.0001; **Extended Data Figure 8G**). In control experiments, AP shape is impervious to all concentrations of STX-bpc, with the exception of a very slight decrease in AP amplitude observed at 1000 nM (2%, p = 0.008; **Extended Data Figure 10**). Thus, prior to light exposure STX-bpc addition to DRGs is functionally invisible, even at 1 µM concentration.

### STX-bpc Performance in Mammalian Brain Tissue

To demonstrate the general utility of STX-bpc for controlling electrical excitability in tissue, we assessed the effects of uncaging this compound in acute cortical slices prepared from mice (**Figure 4**). These experiments were performing with a low power, LED light source (365 nm, < 20 mW/mm^2^). Action potential trains were evoked by 500 ms current steps of 50–150 pA and measured by whole-cell current clamp recording. A LED-only control experiment confirmed that UV light exposure alone, absent toxin or depolarization, did not result in AP generation (**Figure 4A–B**). Application of 250 nM STX-bpc showed minimal reduction in AP generation prior to light delivery; conversely, photo-uncaging resulted in rapid and efficient block of AP generation (**Figure 4A–B**).

Further analyses of AP waveforms in slice recordings were conducted to characterize the effects of uncaging on signal propagation (**Figure 4C–I**). Examination of the first and second time derivatives of individual fractionated APs reveals two peaks that appear in the initial rising phase of the action potential (**Figure 4C–D**). These peaks correspond to activation of Na_V_s located in the axon initial segment (AIS, first peak) and subsequent activation of somatic Na_V_s (second peak) as the AP backpropagates from its initiation point in the AIS to the soma and dendritic compartments.^35^ Such data allow us to assay the effects of the uncaged toxin on the inhibition of different subpopulations of Na_V_s within the axon and somatodendritic compartments. Following uncaging of 250 nM STX-bpc, the amplitude and maximum first and second time derivative values decrease in APs before sufficient Na_V_ block completely prevents AP initiation (**Figure 4D**). No difference in the relative attenuation of axonal vs somatodendritic responses (AIS/DS peak ratio) was detected, indicating Na_V_s in these subcompartments are blocked equivalently by uncaging of STX-bpc (**Figure 4H**).

To determine the duration of light exposure required to achieve complete block of AP generation in cortical neurons from 250 nM STX-bpc uncaging, we recorded from layer 4 cortical neurons (L4) while injecting a continual depolarizing current step to elicit a sustained train of action potentials (**Figure 4J**). In this experiment, the LED remained on until AP generation (**Figure 4J–L**) had completely ceased. Progressive attenuation of AP rate of rise and peak are apparent in the phase plot of a selected recording (**Figure 4K**). Histograms of each cell recording show that complete block of AP generation is achieved in most cells within seconds following exposure to LED light (**Figure 4L**).

To assess the effect of uncaging on cortical networks, we exposed cortical slices maintained at the air liquid interface to 250 nM STX-bpc while activating network responses through electrical stimulation within cortical layer 5 (L5) (**Extended Data Figure 12**). Network responses were recorded by linear multielectrode arrays arranged across all cortical layers. From local field potential recordings, current source density (CSD) calculations were performed to localize current sources and sinks in response to simulation (**Extended Data Figure 12**). Pharmacological dissection of the CSD response was used to identify CSD components that correspond to presynaptic and postsynaptic activity (**Extended Data Figure 12A**). Using this sensitive measure of cortical network activity, slices were exposed to steps of increasing light intensity until all activity was abolished (**Extended Data Figure 12B–D**). No significant separation between presynaptic and postsynaptic responses are observed even under conditions of partial Na_V_ block at intermediate light exposures (4–20 mW/mm^2^, **Extended Data Figure 12C**). This result indicates that the diverse cellular compartments and Na_V_ isoforms distributed throughout the cortical network are similarly inhibited upon photo-deprotection of STX-bpc.

### Dissection of Zebrafish Swimming by Focal Silencing of Neuronal Activity Using STX-bpc

To assess the performance of STX-bpc *in vivo*, we examined the utility of STX-bpc uncaging for the control of locomotion in larval zebrafish. For these experiments, we tracked tail movements in response to swim-eliciting stimulation of the inner ear before and after light exposure (**Figure 5A**). To validate *in vivo* silencing of neural activity with STX-bpc, we performed a single 50 µM STX-bpc injection into the hindbrain ventricle and focused uncaging light in the brainstem, verifying localization with UV-sensitive photo-activatable GFP (**Figure 5B**). Brief pulses of 365 nm light in the absence of toxin had no effect on behavior (data not shown), nor did injection of the caged toxin, but immediately after uncaging throughout the brainstem, stimulus responses were completely abolished, indicating silencing of brainstem neurons that transform sensation into swimming (**Figure 5C**). The duration and probability of stimulus response interference was dependent on the duration of uncaging, with 500 ms of light exposure abolishing responses for 10 seconds and preventing most responses for the subsequent 4 minutes (**Figure 5D**). Swimming responses returned within 5 minutes following irradiation, presumably due to dissociation of bound toxin. Uncaging from the available pool of STX-bpc can be repeated with recovery of block of locomotor function over several cycles (**Figure 5E**).

We next sought to make nuanced changes to swimming behavior through focal control of neuronal function. Head rotation and auditory stimuli are transduced by both ears, eliciting bilateral activity in the brainstem that provides descending input to the spinal cord for bending the tail.^36^ To manipulate swimming responses to our symmetrical stimulation of the ears, we precisely localized uncaging light to the left octavolateral hindbrain (**Figure 5F**) to unilaterally silence sensory neurons that encode the stimulus and ultimately drive swimming. After injection but before uncaging of STX-bpc, swimming responses were symmetrical, but after uncaging swimming was transiently and exclusively contraversive, bending to the unaffected side. The observed behavior suggests focal silencing inhibits the recruitment of muscles on the silenced side (**Figure 5G–5I, Extended Data Video 1**). Swimming responses were reliably evoked and consistently asymmetric immediately following uncaging, and leftward tail bends re-emerged within 30 seconds and gradually recovered until responses were again symmetrical 6 minutes later. Uncaging also directly elicited tail bends away from the inhibited side (**Figure 5H–5I, Extended Data Video 1**), consistent with acute disinhibition of the right brainstem through loss of commissural inhibition from the site of uncaging on the left. Importantly, spontaneous swims following uncaging of STX-bpc were still symmetrical (data not shown), in striking contrast to stimulus-evoked swims, suggesting focal release of **1** interfered with the sensorimotor transformation but not movement generation broadly. Thus, focal uncaging of STX-bpc enables high-precision control and tuning of Na_V_ current, electrical excitability, and finally, behavior, in a rapid, reproducible, and reversible manner.

## DISCUSSION

We have designed and validated STX-bpc **13** as a Na_V_-selective small molecule tool for the spatiotemporal manipulation of electrical transmission *in vitro* and *in vivo*. STX-bpc incorporates multiple carboxylate groups on a modified nitrobenzyl platform to destabilize toxin binding to Na_V_s.^23^ Na_V_ block following light-promoted deprotection of STX-bpc proceeds more rapidly (t_1/2_ ≤ 1.1 seconds at all concentrations tested; **Extended Data Figure 4, 5**) than laser-induced uncaging of STX-coumarin derivatives^23^ and other caged toxins previously described.^37^ The inertness of STX-bpc enables experiments over a sizable concentration window (≤ 500 nM), thus affording a large dynamic range to regulate AP signaling. Brief light pulses using either a 355 nm laser or low-cost LEDs are sufficient to promote inhibitor release from STX-bpc.

The efficacy of STX-bpc uncaging in dissociated cells, tissue slice, and zebrafish establishes the utility of this tool compound. Uncaging STX-bpc elicits dose-dependent decreases in AP firing rate in both dissociated hippocampal and DRG neurons, the latter in spite of the preponderance of STX-resistant Na_V_s (Na_V_1.8, 1.9) in sensory neurons—a result likely due to the predominant role of Na_V_1.7, a STX-sensitive channel in rat, for effecting threshold depolarization.^38^ In hippocampal neurons, a comparison of Na_V_ current block to AP frequency reduction is in keeping with our prior work^23^ (**Extended Data Figure 11A**). Our results show that AP firing rate is largely insensitive to a small percentage of Na_V_ block (≤ 11% inhibition of peak current),^33^ but dramatically reduced if inhibition of peak current ≥67%. AP firing rate in DRG neurons shows greater resistance to block by **1**, with >2-fold the concentration of STX-bpc required to elicit similar reductions in AP frequency (**Extended Data Figure 11B**). Analysis of the AP waveform in both dissociated neurons and acute cortical slice reveals that STX-bpc is inert prior to uncaging, but following deprotection, **1** impacts all Na_V_-dependent components of the AP (firing threshold, velocity, acceleration). Additionally, analysis of the AIS/DS peak ratio shows that STX-bpc uncaging exerts similar effects on both axonal and somatodendritic Na_V_s, establishing the potential of this tool to target sodium channels in subcellular compartments.^35^

The effectiveness of STX-bpc uncaging is highlighted for behavioral control *in vivo* in larval *D. rerio*. As in experiments performed with rodent neurons and tissue slice, STX-bpc remains functionally inert prior to LED-induced uncaging, at which point the released **1** rapidly and selectively abrogates activity. Application of STX-bpc to the entire nervous system has no apparent effect on behavior, permitting inducible and focal uncaging upon light exposure. Behavioral effects are restricted to the site of uncaging, as swimming manipulations are kinematically nuanced and context-specific. Furthermore, the rapid recovery from uncaging (∼5 minutes) coupled with replenishment of caged STX-bpc from the injected pool enables toggling of activity between silenced and active states and behavior from manipulated to intact. Resultant changes in zebrafish locomotion are reproducible, reversible, and spatiotemporally precise.

We have demonstrated that STX-bpc enables optically-induced control of electrical activity and neuronal silencing in cells, tissue, and zebrafish. This reagent operates absent the need for specialized equipment or extensive optimization and, most importantly, without genetic engineering. Accordingly, STX-bpc should serve as a valuable complement to available optogenetic methods for the control of electrical activity, cellular communication, and behavior, and may find application in non-model organisms for which efficient genetic manipulation methods are not yet optimized.

## METHODS

### Synthesis

Detailed preparations for all photocaged toxins are described in the extended data.

### Cell Culture

#### Chinese hamster ovary (CHO) cells stably expressing rat Na_V_ 1.2

Stably-expressing Na_V_1.2 CHO cells were generously provided by Dr. W. A. Catterall (University of Washington, Department of Pharmacology). Cells were grown on 10 cm tissue culture dishes in RPMI 1640 medium with L-glutamine (Thermo Fisher, Waltham, MA) and supplemented with 10% fetal bovine serum (ATCC, Manassas, VA), 50 U/mL penicillin-streptomycin (Thermo Fisher, Waltham, MA), and 0.2 µg/mL G418 (Sigma-Aldrich Co., St. Louis, MO). Cells were kept in a 37 °C, 5% carbon dioxide, 96% relative humidity incubator and passaged approximately every three days. Passaging of cells was accomplished by aspiration of media, washing with phosphate-buffered saline, treatment with 1 mL of trypsin-EDTA (0.05% trypsin, Millipore Sigma, Hayward, CA) until full dissociation of cells was observed (approximately five minutes), and dilution with 4 mL of growth medium. Cells were routinely passaged at 1 in 20 to 1 in 10 dilution.

#### Sprague Dawley rat embryonic day 18 hippocampal neurons

Prior to dissection, 5 mm diameter, 0.15 mm thick round glass coverslips (Warner Instruments, Hamden, CT) were coated overnight with 1 mg/mL poly-D-lysine hydrobromide (PDL, molecular weight 70,000–150,000 Da, Millipore Sigma, Hayward, CA) in 0.1 M, pH 8.5 borate buffer in a 37 °C, 5% carbon dioxide, 96% relative humidity incubator.

Hippocampi were dissected from embryonic day 18 fetuses into ice-cold Hibernate E (BrainBits, LLC, Springfield, IL) as previously described.^39^ Following dissection, cells were dissociated in 2 mL of trypsin-EDTA for 15 minutes in a 37 °C, 5% carbon dioxide, 96% relative humidity incubator. Subsequently, trypsinized cells were quenched with 10 mL of quenching medium (DMEM high glucose (Thermo Fisher, Waltham, MA) supplemented with 15% fetal bovine serum, 100 U/mL penicillin-streptomycin, and 1 mM MEM sodium pyruvate (Atlanta Biologicals, Flowery Branch, GA). Tissue was allowed to settle, supernatant was removed, and the tissue pellet was rinsed twice more with 10 mL of quenching medium. Cells were then manually dissociated into 2 mL of plating medium (DMEM supplemented with 10% FBS, 50 U/mL penicillin-streptomycin, and 1 mM MEM sodium pyruvate) by pipetting with a fire-polished 9” borosilicate glass Pasteur pipet (Fisher Scientific, Waltham, MA).

Cells were plated onto PDL-coated 5 mm glass coverslips in 35 mm tissue culture dishes containing 2 mL of plating medium at a density of 200,000 cells/dish (for voltage-clamp experiments) or 600,000 cells/dish (for current-clamp experiments). After 45 minutes, coverslips were transferred to new tissue culture dishes containing 2 mL of maintenance medium (neurobasal supplemented with 1x B-27 Supplement, 1x Glutamax, and 50 U/mL penicillin-streptomycin (Thermo Fisher, Waltham, MA)). Cells were fed every 3–4 days by changing 50% of the working medium.

#### Sprague Dawley rat embryonic day 15 dorsal root ganglia (DRG) neurons

Dissociated DRG cells were prepared in an analogous manner to hippocampal neurons, with the following exceptions. First, coverslips were coated with 10 µg/mL PDL and 2 µg/mL mouse laminin I (Bio-Techne, Minneapolis, MN) as previously described by Zuchero.^40^ Second, after treating trypsinized cells with quenching media, the working solution was centrifuged at 1000 x g for 30 s to yield a cell pellet. The supernatant was discarded, and the pellet was immediately dissociated into plating medium, as described above. Following dissociation, cells were plated in plating media at a density of 200,000 cells/dish for both voltage-and current-clamp experiments. After 2–3 hours, coverslips were transferred to new culture dishes containing 2 mL of DRG maintenance medium (neurobasal supplemented with 1x B-27 Supplement, 1x Glutamax, 50 U/mL penicillin-streptomycin, 5 mg/mL NaCl, 7.5 µg/mL 5-fluoro-2’-deoxyuridine, 17.5 µg/ml uridine, and 100 ng/mL HPLC-purified nerve growth factor (NGF) 2.5S, beta subunit (Cedarlane, Ontario, Canada)).^41^ Cells were fed every 2–4 days by changing 50% of the working medium.

#### Patch clamp electrophysiology on CHO cells and dissociated neurons

Data were measured using the patch-clamp technique in the whole-cell configuration with an Axon Axopatch 200B amplifier (Molecular Devices, San Jose, CA). The output of the patch-clamp amplifier was filtered with a built-in low-pass, four-pole Bessel filter having a cutoff frequency of 5 kHz for voltage-clamp recordings or 10 kHz for current-clamp recordings, and sampled at 100 kHz. Pulse stimulation and data acquisition used Molecular Devices Digidata 1322A or 1550B controlled with pCLAMP software version 10.4 or 11.1, respectively (Molecular Devices, San Jose, CA).

Borosilicate glass micropipettes (Sutter Instruments, Novato, CA) were fire-polished to a tip diameter yielding a resistance of 1.3–5.5 MΩ, for Na_V_1.2 CHO cells or neurons in slice, or 3.0–9.0 MΩ, for dissociated neurons, in the working solutions.

#### Voltage-clamp recordings

For Na_V_1.2 CHO and CHO-K1 cells, the internal solution was composed of 40 mM NaF, 1 mM EDTA, 20 mM HEPES, and 125 mM CsCl (pH 7.4 with CsOH); the external solution comprised 160 mM NaCl, 2 mM CaCl_2_, and 20 mM HEPES (pH 7.4 with CsOH). For E18, DIV 6–8 hippocampal neurons, the internal solution was composed of 114.5 mM gluconic acid, 114.5 mM CsOH, 2 mM NaCl, 8 mM CsCl, 10 mM MOPS, 4 mM EGTA, 4 mM MgATP, and 0.3 mM Na_2_GTP (pH 7.3 with CsOH, 240 mOsm with glucose), and the external solution was Hibernate E low fluorescence (BrainBits,

LLC, Springfield, IL). For E15, DIV 6–22 DRGs, the internal and external solutions were prepared as described by Cummins, *et al*.^42^ Currents were elicited by 10 ms step depolarizations from a holding potential (–100 mV for Na_V_1.2 CHO cells or –80 mV for dissociated neurons) to 0 mV at a rate of 0.5 Hz (unless otherwise noted). Leak currents were subtracted using a standard P/4 protocol of the same polarity (for Na_V_1.2 CHO cells) or opposite polarity (for dissociated neurons). Series resistance was compensated at 80–95% with a τ_lag_ of 20 or 35 ms for Na_V_1.2 CHO cells dissociated neurons, respectively. All measurements were recorded at room temperature (20– 25 °C). Data were normalized to control currents, plotted against toxin concentration, and analyzed using Prism 8 (GraphPad Software, LLC). Data were fit to concentration-response curves to obtain IC_50_ values and expressed as mean ± s.e.m. The number of observations (n) was ≥ 3 for all reported data. For IC_50_ measurements, cells were used if they exhibited currents greater than 1 nA upon depolarization and I/I_0_ was within 10% of initial current upon toxin washout.

#### Current-clamp recordings

Data were collected on hippocampal and DRG neurons firing action potential trains with frequencies greater than 5 Hz. For the E18, DIV 9–13 hippocampal neurons, the internal solution was composed of 130 mM CH_3_SO ^−^K^+^, 8 mM NaCl, 10 mM HEPES, 10 mM Na_2_phosphocreatine, 3 mM L-ascorbic acid, 4 mM MgATP, and 0.4 mM Na_2_GTP (pH 7.4 with KOH, 305 mOsm with glucose); the external solution comprised Hibernate E low fluorescence adjusted to 310 mOsm with 40 mM NaCl. For E15, DIV 6–8 DRG neurons, the internal and external solutions were prepared as described by Cummins, *et al*.^42^

Action potentials were elicited by four 500 ms current injections of 50–150 pA in hippocampal neurons or 250–550 pA in DRG cells, at a rate of 0.25 Hz. Series resistance was typically compensated at 90–95% with a τ_lag_ of 35 ms. All measurements were recorded at room temperature (20–25 °C). Data were analyzed using Clampfit (Molecular Devices, San Jose, CA) and Prism 8 (GraphPad Software, LLC). All data represent mean ± s.e.m. for n (the number of observations) ≥ 3. Statistical comparisons were performed as described in figure legends. Phase plots were graphed by plotting the first derivative of the AP against its recorded potential. AP threshold and onset rapidness were calculated as described by Lazarov, *et al*.^43^

### Patch clamp electrophysiology on mouse brain slice

#### Current-clamp recordings

All procedures were performed in accordance with the protocols approved by the Institutional Animal Care and Use Committee (IACUC) of Columbia University. P15–30 mice were quickly decapitated under isoflurane sedation. Coronal slices (300 µm) from somatosensory barrel cortex (S1bf). were prepared in ice-cold sucrose cutting solution containing 2.5 mM KCl, 10 mM MgCl_2_, 0.5 mM CaCl_2_, 1.25 mM NaH_2_PO_4_, 26 mM NaHCO_3_, 234 mM sucrose, 11 mM glucose using a vibratome (Leica VT1200). Slices were then transferred to artificial cerebral spinal fluid (aCSF) containing 126 mM NaCl, 26 mM NaHCO_3_, 2.5 mM KCl, 1.25 mM NaH_2_PO_4_, 2 mM CaCl_2_, 1 mM MgCl_2_, and 10 mM glucose bubbled with 95% O_2_ and 5% CO_2_ at 32 °C for 30 minutes. The slices were then maintained at room temperature until the experiment was performed.

Action potentials were recorded from layer 4 cortical neurons in aCSF at 32 °C using conventional whole-cell current clamp recording techniques. Neurons were visualized using infrared differential interference contrast (DIC) with a SliceScope Pro (Scientifica) microscope. Data were acquired using a Multiclamp 700B amplifier and a Digidata 1550B digitizer. Signals were recorded at 100 kHz sample rate and filtered with a 10 kHz low-pass Bessell filter using pClamp 11 software (all equipment from Molecular Devices). Patch pipettes were fabricated with a P-80C pipette puller (Sutter Instruments) using 1.5 mm outer diameter, 1.28 mm inner diameter filamented capillary glass (World Precision Instruments). The pipette resistance was 3–6 MΩ.

Pipette intracellular solution contained 127 mM K^+^ gluconate, 8 mM NaCl, 4 mM MgATP, 0.6 mM EGTA, 0.3 mM GTP, 10 HEPES, adjusted to pH 7.3–7.4 with KOH. Osmolarity was adjusted to 290–300 mOsm. Action potentials were evoked using four 500 ms duration current steps every 20 s. The amplitude of the step varied between neurons and was that which evoked half the maximum number of action potentials (60–180 pA). An action potential was defined as a transient depolarization that had a minimum rise rate > 10 mV/ms and reached a peak amplitude > 0 mV. During recordings, an offset current (< 150 pA) was manually injected to hold the cells at approximately −75 mV. Membrane potential values were not corrected for a –15 mV liquid junction potential. Only cells with a stable series resistance < 15 MΩ were used for analysis.

Quantification was carried out using custom-written scripts for Igor Pro v.9 (Wavemetrics) and R v.4 (www.R-project.org). Statistical comparisons were made using Dunnett’s test. P values < 0.05 were considered significant.

#### Extracellular multielectrode electrophysiology

To measure differences in cortical network activity following uncaging of STX, extracellular recordings were obtained using linear multielectrode arrays in mouse brain. Coronal slices (350 µm) were prepared as described above. Slices were placed in an interface recording chamber and maintained in aCSF at 34 ± 1 °C. Neuronal activity was recorded with commercially available linear array probes (Model: A16×1-2mm-100-177, NeuroNexus, Ann Arbor, MI, USA) with 16 contacts (spacing: 100 µm distance) spanning 1500 µm. Probes were slowly lowered across cortical layers. A concentric bipolar stimulating electrode (World Precision Instruments) was placed approximately 300 µm from the probe location. Data acquisition was performed through a digitizing board (SI-4, Tucker-Davis Technologies) connected to a real-time acquisition processor (RZ10x, Tucker-Davis Technologies) and PC workstation (WS-8, Tucker-Davis Technologies) running custom-written routines in Synapse (Tucker-Davis Technologies). Recordings were sampled at 24 kHz.

#### Current source density analyses

Current source density (CSD) analysis was performed to estimate the density of transmembrane current sources. Computations were performed in Python 3.7 using the Elephant 0.7.0 package, implementing the 1-dimensional electrode set-up kernel CSD method for a non-parametric CSD estimation from arbitrarily distributed electrodes.^44^ Cross-validation was performed to prevent over-fitting. CSD data were represented as an *m* × *n* matrix *C*. Current volumes, *Vol*(*C*), were calculated as in Equation 1 in which the absolute value of all CSD sink and source values in matrix *C* were summed.

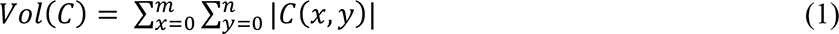

Data visualizations were created using Python Pandas, Scipy, Season, NumpPy and Matplotlib packages. Statistical significance was determined using a Tukey’s HSD test. A *p* < 0.05 was considered significant.

### UV light delivery

UV light was delivered to CHO cells and dissociated hippocampal and DRG neurons using a pulsed 355 nm UV laser (DPSS Lasers, Model 3501-100) directed through a 200 µm core optical fiber to a 200 µm core, 0.22 NA fiber optic cannula (Thorlabs, Newton, NJ) to the clamped cell. Unless otherwise noted, photo-uncaging was induced by 5 ms, 130 mW UV pulses activated immediately prior to the depolarization (or current injection) step. For all slice recordings, a pE-300 Ultra UV LED (Cool LED, Andover, UK) was used to deliver 365 nm light through a 20x objective (Olympus, Tokyo, JP).

### Focal inactivation of neural circuits in behaving zebrafish

Larval zebrafish at 4–7 days post-fertilization were anesthetized (0.025% MS-222, Syndel) and mounted in 2% low melting point agarose (Thermo Fisher Scientific 16520), dorsal-up, for injection of toxin into the hindbrain ventricle.^45^ Larvae of the *mitfa-/-* skin pigmentation mutant were used to visualize the nervous system.^46^ Larvae were visualized under a 20x water dipping objective (XLUMPLFLN20XW, Olympus) of an upright, epifluorescent microscope (BX51WI, Olympus). A microinjection needle (1.7 μm diameter tip) was prepared on a micropipette puller (P-97, Sutter Instrument) and backfilled with 50 μM STX-bpc for injection of a small bolus. Then, the micropipette tip was visually inserted into the ventricle using a micromanipulator (uMp-3, Sensapex). Toxin was pressure injected using an inline Openspritzer^47^ based on initial calibration to 45 psi and 10 msec pulse duration by visualizing the injection volume of 128 μM sulfrhodamine B (Sigma-Aldrich). Following toxin injection, the anesthetized larva was freed from agarose, allowed to recover from anesthesia in fresh E3, and mounted in a bead of agarose on the reverse side of the mirror of a galvanometer (GVS011, ThorLabs). The larva was aligned rostrocaudally with the axis of rotation. The embedded larva was immersed in E3 and had its tail freed with a no. 11 scalpel by removing an isosceles triangle of agarose with its apex at the rostral tail and its base caudal to the tip of the tail. The galvanometer mirror was affixed under the same 20x objective by mounting in a custom 3D-printed clamp. Rotational stimuli were delivered through the galvanometer as 5 cycles of 1 kHz sine waves at 10 deg amplitude. Swimming responses were recorded at 400 fps with an infrared camera (Blackfly S, BFS-U3-16S2M-CS, FLIR) and fixed magnification lens (InfiniStix 0.5X, Infinity Photo-Optical Company) focused on tails illuminated with 940 nm infrared LED array (Homyl). The vibration stimulus was delivered to the galvanometer in concert with uncaging light using analog outputs of a patch clamp amplifier and stimuli designed in SutterPatch software (dPatch, Sutter Instrument). The analog uncaging trigger was used to drive illumination through the microscope objective with a LED driver (Cyclops, OpenEphys) and collimated ultraviolet LED (M365LP1-C1, ThorLabs). The UV illumination was constrained with the microscope field diaphragm and targeted within the brain by translating the fish laterally with a manual stage (MXMS-115, Siskiyou Corporation). To evaluate the localization of UV light within the brain, zebrafish expressing photoactivatable GFP (*Tg:alpha-tubulin:C3PA-GFP*) were illuminated in the uncaging setup and then imaged on a confocal microscope (FV1000, Olympus).^48^ To compute tail deflection angles from videos of embedded larvae, each larva had a dedicated artificial neural network trained on 6 points along the midline of its tail using DeepLabCut.^49^ Angles were calculated in Matlab and absolute angle integrated to compute swimming vigor (Mathworks).

### Ethics oversight

All animal care and dissection procedures were performed in accordance with NIH guidelines for the use of experimental animals and approved by the Stanford Administrative Panel on Laboratory Animal Care (APLAC), the Columbia University Institutional Animal Care and Use Committee, or the University of Wisconsin-Madison Institutional Animal Care and Use Committee.

## DATA AVAILABILITY

The datasets used the current study are available from the corresponding authors on request. All synthetic characterization data generated during this study are included in the supplemental information files. Source data for all main and supplementary figures are provided as a Source Data file.

## CODE AVAILABILITY

Python code used for analysis of action potential properties in rodent brain slice is available at GitHub.com/MakinsonLab.

## ACKNOWLEDGEMENTS

This work was supported by a National Institutes of Health grant R21NS107003 to J.D. and by R01 GM117263-05 (J.D.). C.D.M was supported by the National Institutes of Health, National Institute of Neurological Disorders and Stroke R00NS104215, and National Institute of Mental Health DP2 MH132944. D.E.E. and N.M. were supported by the National Institute of General Medical Sciences of the National Institutes of Health under award R35GM146885. J.S. was supported by the NIH BRAIN Initiative grant F32H130023. The authors would like to thank David Schoppik for providing zebrafish lines and Rodrigo Zúñiga Mouret for equipment fabrication. A.V.E. was supported by the Stanford University Center for Molecular Analysis and Design (CMAD) as well as a Stanford Interdisciplinary Graduate Fellowship (SIGF) through the Stanford Bio-X Interdisciplinary Biosciences Institute. A.L.H. was supported by a Stanford Chemistry Undergraduate Summer Research Fellowship. C.L. was supported by the National Science Foundation Graduate Research Fellowships Program (GRFP) DGE-2036197. This research was performed using the computational resources and assistance of the UW-Madison Center For High Throughput Computing (CHTC) in the Department of Computer Sciences. The CHTC is supported by UW-Madison, the Advanced Computing Initiative, the Wisconsin Alumni Research Foundation, the Wisconsin Institutes for Discovery, and the National Science Foundation, and is an active member of the OSG Consortium, which is supported by the National Science Foundation and the U.S. Department of Energy’s Office of Science.

## AUTHOR CONTRIBUTIONS

A.V.E., C.D.M., D.E.E., and J.D. conceived the project. A.V.E. and J.D. designed photocaged STXs. A.V.E. and A.L.H. synthesized all photocaged STXs and chemical intermediates. A.V.E. collected potency and uncaging data against Na_V_1.2 CHO cells and singly dissociated hippocampal and DRG neurons. D.W., C.L., J.S., and C.D.M. performed all slice physiology experiments. N.M. and D.E.E. performed all work in *D. rerio*. All authors contributed to data analysis and the writing of the manuscript.

## REFERENCES

1. Kim TH, Schnitzer MJ. Fluorescence imaging of large-scale neural ensemble dynamics. Cell. 2022 Jan 6;185(1):9–41.

2. Jacobs J, Kahana MJ. Direct brain recordings fuel advances in cognitive electrophysiology. Trends Cogn Sci. 2010 Apr;14(4):162–71.

3. Deisseroth K. Optogenetics: 10 years of microbial opsins in neuroscience. Nat Neurosci. 2015 Sep;18(9):1213–25.

4. Yizhar O, Fenno LE, Davidson TJ, Mogri M, Deisseroth K. Optogenetics in neural systems. Neuron. 2011 Jul 14;71(1):9–34.

5. Boyden ES, Zhang F, Bamberg E, Nagel G, Deisseroth K. Millisecond-timescale, genetically targeted optical control of neural activity. Nat Neurosci. 2005 Sep;8(9):1263–8.

6. Aghayee A, Ashton R. Methods for Controlled Induction of Singular Rosette Cytoarchitecture Within Human Pluripotent Stem Cell-Derived Neural Multicellular Assemblies. Methods Mol Biol. 2021;2258:193–203.

7. Mahn M, Gibor L, Patil P, Cohen-Kashi Malina K, Oring S, Printz Y, Levy R, Lampl I, Yizhar O. High-efficiency optogenetic silencing with soma-targeted anion-conducting channelrhodopsins. Nat Commun. 2018 Oct 8;9(1):4125.

8. Miyashita T, Shao YR, Chung J, Pourzia O, Feldman DE. Long-term channelrhodopsin-2 (ChR2) expression can induce abnormal axonal morphology and targeting in cerebral cortex. Front Neural Circuits. 2013 Jan 31;7:8.

9. Hudry E, Vandenberghe LH. Therapeutic AAV Gene Transfer to the Nervous System: A Clinical Reality. Neuron. 2019 Apr 3;102(1):263.

10. Hamada S, Nagase M, Yoshizawa T, Hagiwara A, Isomura Y, Watabe AM, Ohtsuka T. An engineered channelrhodopsin optimized for axon terminal activation and circuit mapping. Commun Biol. 2021 Apr 12;4(1):461.

11. Zimmermann D, Zhou A, Kiesel M, Feldbauer K, Terpitz U, Haase W, Schneider-Hohendorf T, Bamberg E, Sukhorukov VL. Effects on capacitance by overexpression of membrane proteins. Biochem Biophys Res Commun. 2008 May 16;369(4):1022–6.

12. Messier JE, Chen H, Cai ZL, Xue M. Targeting light-gated chloride channels to neuronal somatodendritic domain reduces their excitatory effect in the axon. Elife. 2018 Aug 9;7:e38506.

13. Hayward RF, Brooks FP, Yang S, Gao S, Cohen AE. Diminishing neuronal acidification by channelrhodopsins with low proton conduction. bioRxiv [Preprint]. 2023 Feb 8:2023.02.07.527404. doi: 10.1101/2023.02.07.527404.

14. Shen Y, Campbell RE, Côté DC, Paquet ME. Challenges for Therapeutic Applications of Opsin-Based Optogenetic Tools in Humans. Front Neural Circuits. 2020 Jul 15;14:41.

15. Shen H, Liu D, Wu K, Lei J, Yan N. Structures of human Nav1.7 channel in complex with auxiliary subunits and animal toxins. Science. 2019 Mar 22;363(6433):1303-1308.

16. Shen H, Li Z, Jiang Y, Pan X, Wu J, Cristofori-Armstrong B, Smith JJ, Chin YKY, Lei J, Zhou Q, King GF, Yan N. Structural basis for the modulation of voltage-gated sodium channels by animal toxins. Science. 2018 Oct 19;362(6412):eaau2596.

17. Mulcahy JV, Pajouhesh H, Beckley JT, Delwig A, Du Bois J, Hunter JC. Challenges and Opportunities for Therapeutics Targeting the Voltage-Gated Sodium Channel Isoform NaV1.7. J Med Chem. 2019 Oct 10;62(19):8695–8710.

18. Penzotti JL, Fozzard HA, Lipkind GM, Dudley SC Jr. Differences in saxitoxin and tetrodotoxin binding revealed by mutagenesis of the Na+ channel outer vestibule. Biophys J. 1998 Dec;75(6):2647–57.

19. Tanaka JC, Doyle DD, Barr L. Sodium channels in vertebrate hearts. Three types of saxitoxin binding sites in heart. Biochim Biophys Acta. 1984 Aug 22;775(2):203–14.

20. Gordon D, Zlotkin E, Catterall WA. The binding of an insect-selective neurotoxin and saxitoxin to insect neuronal membranes. Biochim. Biophys. Acta. 1985; 821, 130–136.

21. Sattelle DB, Pelhate M, Hue B. Pharmacological properties of axonal sodium channels in the cockroach Periplaneta americana L. I. Selective block by synthetic saxitoxin. J Exp Biol. 1979 Dec;83:41–8.

22. Alonso E, Alfonso A, Vieytes MR, Botana LM. Evaluation of toxicity equivalent factors of paralytic shellfish poisoning toxins in seven human sodium channels types by an automated high throughput electrophysiology system. Arch Toxicol. 2016 Feb;90(2):479–88.

23. Elleman AV, Devienne G, Makinson CD, Haynes AL, Huguenard JR, Du Bois J. Precise spatiotemporal control of voltage-gated sodium channels by photocaged saxitoxin. Nat Commun. 2021 Jul 7;12(1):4171.

24. Furuta T, Wang SS, Dantzker JL, Dore TM, Bybee WJ, Callaway EM, Denk W, Tsien RY. Brominated 7-hydroxycoumarin-4-ylmethyls: photolabile protecting groups with biologically useful cross-sections for two photon photolysis. Proc Natl Acad Sci U S A. 1999 Feb 16;96(4):1193–200.

25. Klán P, Šolomek T, Bochet CG, Blanc A, Givens R, Rubina M, Popik V, Kostikov A, Wirz J. Photoremovable protecting groups in chemistry and biology: reaction mechanisms and efficacy. Chem Rev. 2013 Jan 9;113(1):119–91.

26. Beier M, Hoheisel JD. Production by quantitative photolithographic synthesis of individually quality checked DNA microarrays. Nucleic Acids Res. 2000 Feb 15;28(4):E11.

27. Specht A, Goeldner M. 1-(o-Nitrophenyl)-2,2,2-trifluoroethyl ether derivatives as stable and efficient photoremovable alcohol-protecting groups. Angew Chem Int Ed Engl. 2004 Apr 2;43(15):2008-12.

28. Aujard I, Benbrahim C, Gouget M, Ruel O, Baudin JB, Neveu P, Jullien L. o-nitrobenzyl photolabile protecting groups with red-shifted absorption: syntheses and uncaging cross-sections for one-and two-photon excitation. Chemistry. 2006 Sep 6;12(26):6865–79.

29. Hansch C, Leo A, Taft RW. A Survey of Hammett Substituent Constants and Resonance and Field Parameters. Chem. Rev. 1991;91:165–195.

30. Fleming JJ, Du Bois J. A synthesis of (+)-saxitoxin. J Am Chem Soc. 2006 Mar 29;128(12):3926–7.

31. Mulcahy JV, Walker JR, Merit JE, Whitehead A, Du Bois J. Synthesis of the Paralytic Shellfish Poisons (+)-Gonyautoxin 2, (+)-Gonyautoxin 3, and (+)-11,11-Dihydroxysaxitoxin. J Am Chem Soc. 2016 May 11;138(18):5994-6001.

32. Berroy P, Viriot ML, Carré MC. Photolabile group for 5′-OH protection of nucleosides: Synthesis and photodeprotection rate. Sensors Actuators, B Chem. 2001;74:186–189.

33. Madeja M. Do neurons have a reserve of sodium channels for the generation of action potentials? A study on acutely isolated CA1 neurons from the guinea-pig hippocampus. Eur J Neurosci. 2000 Jan;12(1):1-7.

34. Naundorf B, Wolf F, Volgushev M. Unique features of action potential initiation in cortical neurons. Nature. 2006 Apr 20;440(7087):1060-3.

35. Bean BP. The action potential in mammalian central neurons. Nat Rev Neurosci. 2007 Jun;8(6):451–65.

36. Privat M, Romano SA, Pietri T, Jouary A, Boulanger-Weill J, Elbaz N, Duchemin A, Soares D, Sumbre G. Sensorimotor Transformations in the Zebrafish Auditory System. Curr Biol. 2019 Dec 2;29(23):4010–4023.e4.

37. Montnach J, Blömer LA, Lopez L, Filipis L, Meudal H, Lafoux A, Nicolas S, Chu D, Caumes C, Béroud R, Jopling C, Bosmans F, Huchet C, Landon C, Canepari M, De Waard M. In vivo spatiotemporal control of voltage-gated ion channels by using photoactivatable peptidic toxins. Nat Commun. 2022 Jan 20;13(1):417.

38. Rush AM, Cummins TR, Waxman SG. Multiple sodium channels and their roles in electrogenesis within dorsal root ganglion neurons. J Physiol. 2007 Feb 15;579(Pt 1):1–14.

39. Fath T, Ke YD, Gunning P, Götz J, Ittner LM. Primary support cultures of hippocampal and substantia nigra neurons. Nat Protoc. 2009;4(1):78–85.

40. Zuchero JB. Purification of dorsal root ganglion neurons from rat by immunopanning. Cold Spring Harb Protoc. 2014 Aug 1;2014(8):826–38.

41. Newberry K, Wang S, Hoque N, Kiss L, Ahlijanian MK, Herrington J, Graef JD. Development of a spontaneously active dorsal root ganglia assay using multiwell multielectrode arrays. J Neurophysiol. 2016 Jun 1;115(6):3217–28.

42. Cummins TR, Rush AM, Estacion M, Dib-Hajj SD, Waxman SG. Voltage-clamp and current-clamp recordings from mammalian DRG neurons. Nat Protoc. 2009;4(8):1103–12.

43. Lazarov E, Dannemeyer M, Feulner B, Enderlein J, Gutnick MJ, Wolf F, Neef A. An axon initial segment is required for temporal precision in action potential encoding by neuronal populations. Sci Adv. 2018 Nov 28;4(11):eaau8621.

44. Potworowski J, Jakuczun W, Lȩski S, Wójcik D. Kernel current source density method. Neural Comput. 2012 Feb;24(2):541–75.

45. Horstick EJ, Tabor KM, Jordan DC, Burgess HA. Genetic Ablation, Sensitization, and Isolation of Neurons Using Nitroreductase and Tetrodotoxin-Insensitive Channels. Methods Mol Biol. 2016;1451:355–66.

46. Lister JA, Robertson CP, Lepage T, Johnson SL, Raible DW. nacre encodes a zebrafish microphthalmia-related protein that regulates neural-crest-derived pigment cell fate. Development. 1999 Sep;126(17):3757–67.

47. Forman CJ, Tomes H, Mbobo B, Burman RJ, Jacobs M, Baden T, Raimondo JV. Openspritzer: an open hardware pressure ejection system for reliably delivering picolitre volumes. Sci Rep. 2017 May 19;7(1):2188.

48. Ahrens MB, Li JM, Orger MB, Robson DN, Schier AF, Engert F, Portugues R. Brain-wide neuronal dynamics during motor adaptation in zebrafish. Nature. 2012 May 9;485(7399):471-7.

49. Mathis A, Mamidanna P, Cury KM, Abe T, Murthy VN, Mathis MW, Bethge M. DeepLabCut: markerless pose estimation of user-defined body parts with deep learning. Nat Neurosci. 2018 Sep;21(9):1281–1289.

