## Supporting information for "Behavioral control through the direct, focal silencing of neuronal activity"

### Extended Data

1. Extended Data Figures
2. Extended Data Tables
3. Chemical Procedures
4. $^{1}$H NMR Spectra

### 1. Extended Data Figures

**Extended Data Figure 1.** Full panel of photocaged saxitoxins.

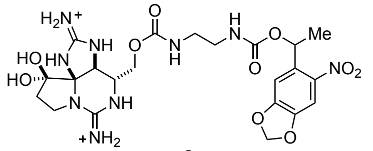

**2**

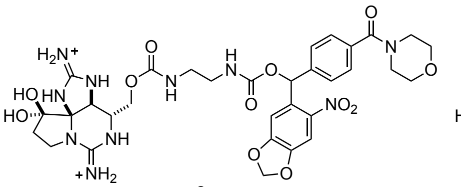

**3**

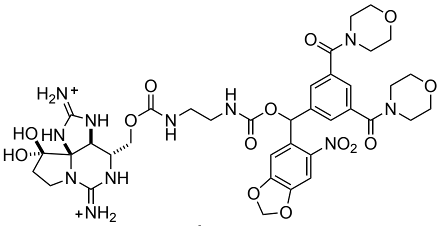

**4**

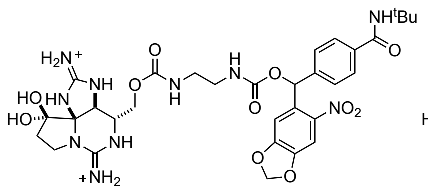

**5**

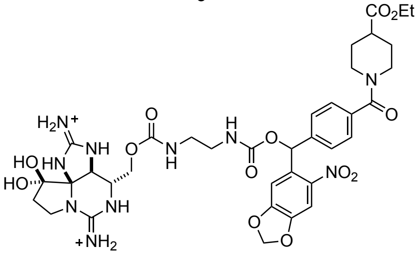

**6**

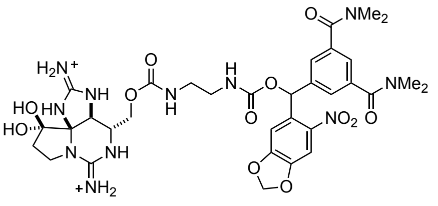

**7**

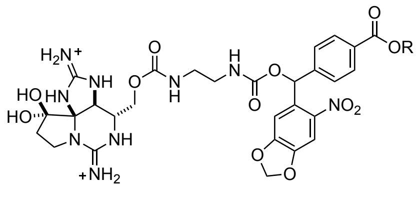

**8**

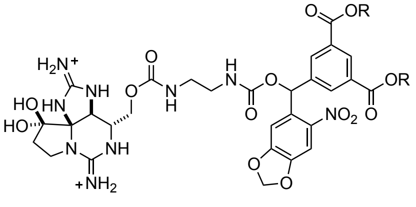

**9**

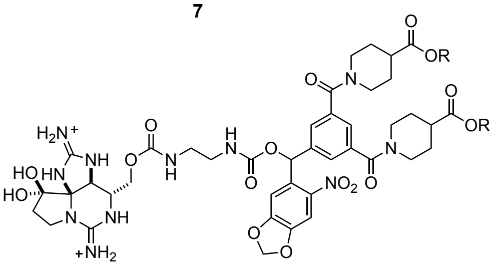

**10**

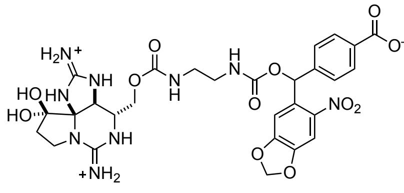

**11**

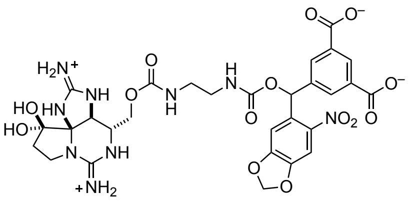

**12**

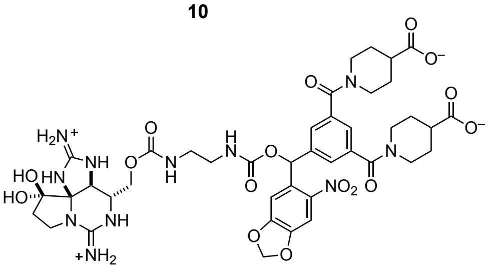

**13**

R = $CH_2CH{=}CH_2$

**Extended Data Figure 2.** Carboxylate-derived photocaged STXs (**11**–**13**) exhibit the lowest potencies and highest uncaging efficiencies. Data collected against rNa$_V$1.2 expressed in CHO cells.

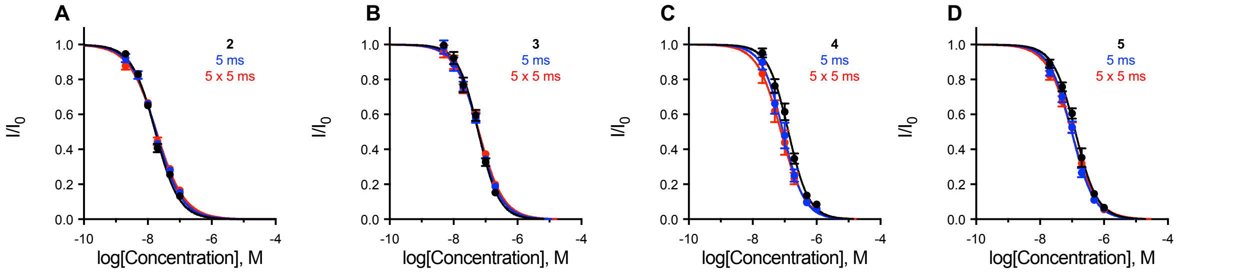


| | 2<br>IC50 (nM), Hill Coefficient | 3<br>IC50 (nM), Hill Coefficient | 4<br>IC50 (nM), Hill Coefficient | 5<br>IC50 (nM), Hill Coefficient |
|---|---|---|---|---|
| **Start** | 17.3 ± 0.7; –1.17 ± 0.05 | 59.8 ± 3.4; –1.35 ± 0.10 | 132.1 ± 7.3; –1.35 ± 0.10 | 114.2 ± 6.5; –1.30 ± 0.10 |
| **5 ms** | 18.4 ± 0.8; –1.08 ± 0.05 | 59.9 ± 3.3; –1.20 ± 0.08 | 90.0 ± 7.3; –1.31 ± 0.14 | 113.6 ± 6.9; –1.23 ± 0.09 |
| **5 x 5 ms** | 18.9 ± 0.9; –1.00 ± 0.05 | 62.7 ± 3.5; –1.14 ± 0.07 | 77.4 ± 6.9; –1.17 ± 0.13 | 120.0 ± 6.6; –1.11 ± 0.07 |

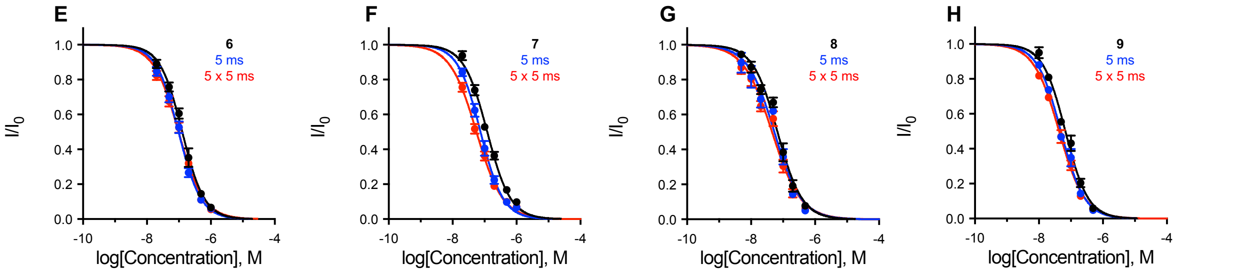


| | 6<br>IC50 (nM), Hill Coefficient | 7<br>IC50 (nM), Hill Coefficient | 8<br>IC50 (nM), Hill Coefficient | 9<br>IC50 (nM), Hill Coefficient |
|---|---|---|---|---|
| **Start** | 128.1 ± 7.2; –1.25 ± 0.09 | 121.9 ± 5.5; –1.19 ± 0.06 | 67.6 ± 4.9; –1.13 ± 0.09 | 67.6 ± 3.6; –1.23 ± 0.08 |
| **5 ms** | 98.1 ± 4.8; –1.24 ± 0.08 | 75.1 ± 3.9; –1.22 ± 0.08 | 53.9 ± 5.3; –1.03 ± 0.10 | 49.0 ± 2.2; –1.18 ± 0.06 |
| **5 x 5 ms** | 98.5 ± 4.6; –1.11 ± 0.06 | 56.0 ± 2.7; –1.05 ± 0.06 | 45.7 ± 3.9; –0.97 ± 0.08 | 43.1 ± 2.0; –1.09 ± 0.06 |

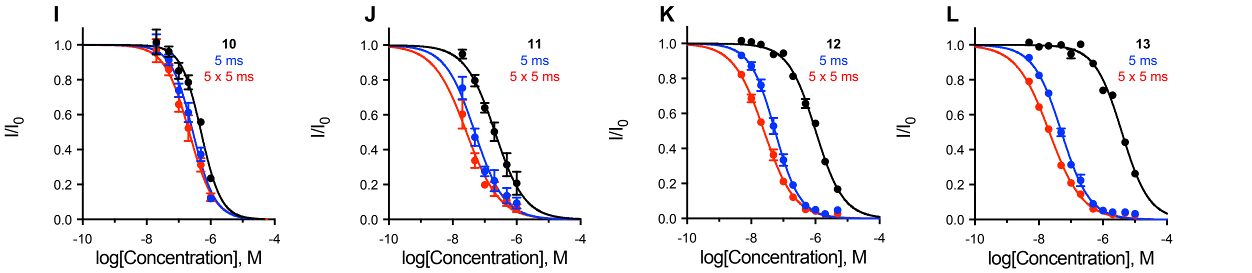


| | 10<br>IC50 (nM), Hill Coefficient | 11<br>IC50 (nM), Hill Coefficient | 12<br>IC50 (nM), Hill Coefficient | 13<br>IC50 (nM), Hill Coefficient |
|---|---|---|---|---|
| **Start** | 507.2 ± 39.0; –1.38 ± 0.15 | 211.3 ± 25.7; –0.93 ± 0.12 | 1024.9 ± 38.6; –1.01 ± 0.04 | 3919.4 ± 172.6; –1.02 ± 0.05 |
| **5 ms** | 281.2 ± 22.3; –1.26 ± 0.12 | 49.5 ± 7.2; –0.93 ± 0.12 | 55.5 ± 2.7; –1.11 ± 0.05 | 51.0 ± 1.9; –0.98 ± 0.03 |
| **5 x 5 ms** | 214.9 ± 20.2; –1.12 ± 0.12 | 27.1 ± 4.6; –0.84 ± 0.11 | 25.1 ± 0.9; –0.91 ± 0.03 | 21.3 ± 0.7; –0.84 ± 0.02 |

(**A**)–(**L**) Electrophysiological characterization of photocaged STXs **2–13** against Na$_V$1.2 CHO. Initial $IC_{50}$ in black; apparent $IC_{50}$ following 5 ms applied laser (blue) and 5 x 5 ms applied laser (red, n = 3–9).

**Extended Data Figure 3.** Photocaged STXs **11–13** have similar absorbance spectra to STX MeNPOC **2**.

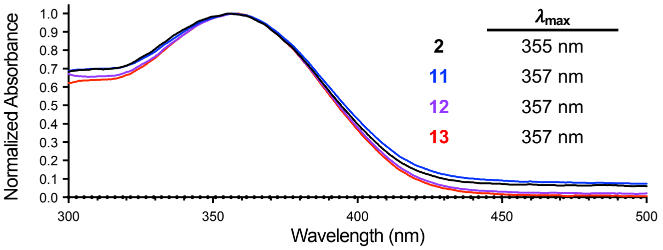


UV/Vis spectra of 2 mM photocaged STXs in water, collected from 2 µL samples on a Thermo NanoDrop One.

**Extended Data Figure 4.** Laser-induced uncaging of **13** effects rapid, concentration-dependent Nav block of Nav1.2 (CHO).

**A**

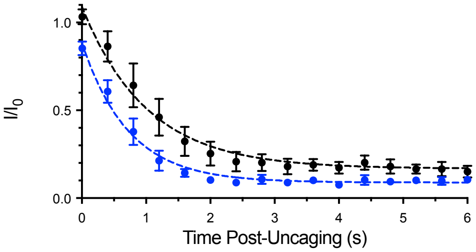


**B**

| [13] | $\tau$ (s) | $t_{1/2}$ (s) | $R^2$ |
|---|---|---|---|
| **200 nM** | 1.01 ± 0.10 | 0.73 ± 0.05 | 0.8367 |
| **500 nM** | 0.70 ± 0.07 | 0.50 ± 0.03 | 0.9132 |

(**A**) Time course of uncaging **13** against Nav1.2 (CHO). Laser was applied at t = 0 s. Data was subjected to one phase decay exponential regression (n = 4). Curves are significantly different ($p < 1 \times 10^{-6}$; extra sum-of-squares F test). (**B**) Table of parameters derived from data plotted in (A).

**Extended Data Figure 5.** Laser-induced uncaging of **13** effects rapid, concentration-dependent Nav block in dissociated hippocampal neurons.

**A**

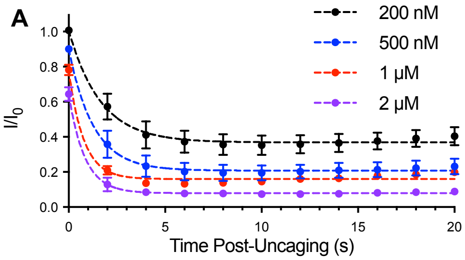


**B**

| [13] | $\tau$ (s) | $t_{1/2}$ (s) | $R^2$ |
|---|---|---|---|
| **200 nM** | 1.64 ± 0.45 | 1.14 ± 0.32 | 0.7759 |
| **500 nM** | 1.28 ± 0.31 | 0.89 ± 0.21 | 0.8473 |
| **1 µM** | 0.77 ± 0.15 | 0.53 ± 0.10 | 0.9483 |
| **2 µM** | 0.83 ± 0.15 | 0.57 ± 0.10 | 0.9575 |

(**A**) Time course of uncaging **13** against dissociated embryonic hippocampal neurons. Laser was applied at t = 0 s. Data was subjected to one phase decay exponential regression (n = 4–5). Curves are significantly different (200 nM vs 500 nM, $p < 1 \times 10^{-6}$; 500 nM vs 1 µM, $p = 5 \times 10^{-6}$; 1 µM vs 2 µM, $p < 1 \times 10^{-6}$; extra sum-of-squares F test). (**B**) Table of parameters derived from data plotted in (A).

**Extended Data Figure 6.** Application of up to 500 nM **13** against dissociated embryonic hippocampal neurons has no effect on AP parameters prior to uncaging.

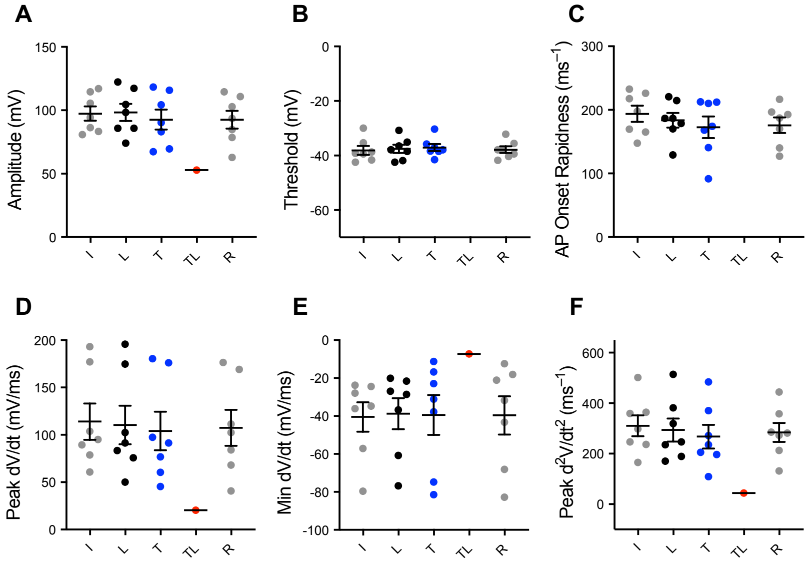


All data calculated from first AP in current step 2. Traces labeled: initial (I), laser applied (L), 500 nM **13** applied (T), 500 nM **13** and laser applied (TL), recovered (R) after wash-off. (**A**) AP amplitude is unchanged by 500 nM **13** or laser application alone. (**B**) AP threshold is unchanged by 500 nM **13** or laser application alone. (**C**) AP onset rapidness is unchanged by 500 nM **13** or laser application alone. (**D**) AP peak rate of rise is unchanged by 500 nM **13** or laser application alone. (**E**) AP peak rate of fall is unchanged by 500 nM **13** or laser application alone. (**F**) Maximal AP acceleration is unchanged by 500 nM **13** or laser application alone (n = 7). (No p values significant, one-way ANOVA with Tukey's correction).

**Extended Data Figure 7.** Uncaging of 200 nM **13** against dissociated embryonic hippocampal neurons reduces AP amplitude and onset rapidness as well as peak rate of rise and maximal acceleration.

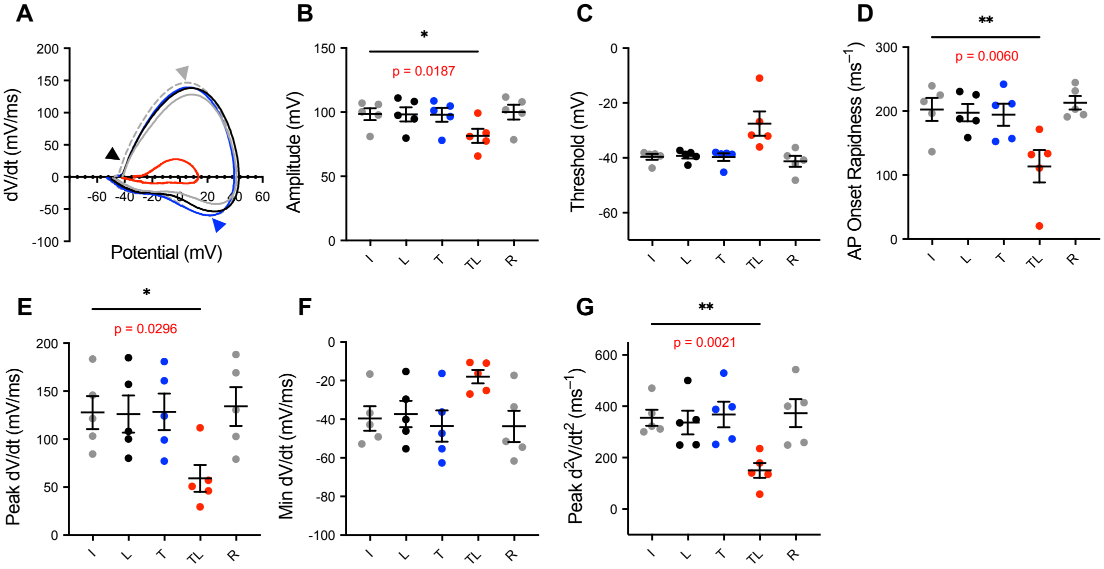


All data calculated from first action potential in current step 2. (**A**) Representative phase plot depicting uncaging of 200 nM **13**. Traces colored/labeled: initial (I), grey; laser applied (L), black; 200 nM **13** applied (T), blue; 200 nM **13** and laser applied (TL), red; and recovered (R) after wash-off, dashed grey. Difference between x-intercepts, amplitude. Black arrow: x-value, threshold; derivative, AP onset rapidness. Grey arrow: y-value, peak dV/dt. Blue arrow: y-value, min dV/dt. (**B**) Reduction in AP amplitude following application and uncaging of **13**. (**C**) Non-significant increase in AP threshold following application and uncaging of **13**. (**D**) Reduction in AP onset rapidness following application and uncaging of **13**. (**E**) Reduction in AP peak rate of rise following application and uncaging of **13**. (**F**) Non-significant increase in AP peak rate of fall following application and uncaging of **13**. (**G**) Reduction in maximal AP acceleration following application and uncaging of **13** (n = 5). (*p<0.05, **p<0.01, one-way ANOVA with Tukey's correction).

**Extended Data Figure 8.** Uncaging of 500 nM **13** against dissociated embryonic dorsal root ganglia neurons reduces AP amplitude, peak rate of rise, and maximal acceleration and increases AP threshold and peak rate of fall.

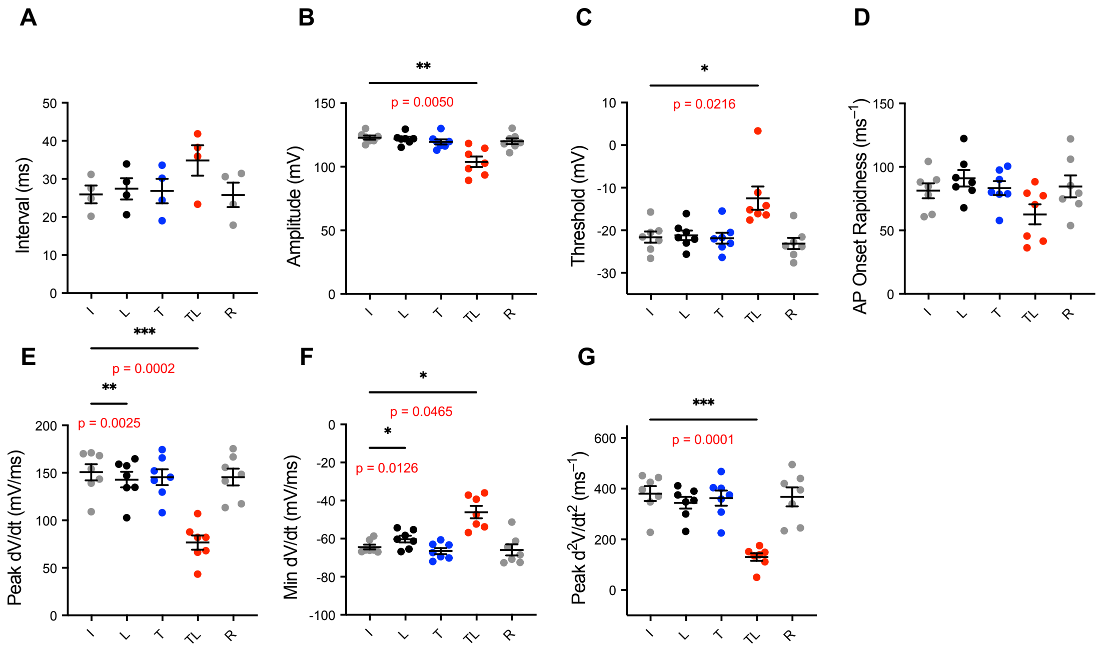


All data calculated from first AP in current step 2. Traces labeled: initial (I), laser applied (L), 500 nM **13** applied (T), 500 nM **13** and laser applied (TL), recovered (R) after wash-off. (**A**) Non-significant increase in interval between first and second AP following application and uncaging of **13**. (**B**) Reduction in AP amplitude following application and uncaging of **13**. (**C**) Increase in AP threshold following application and uncaging of **13**. (**D**) No significant change in AP onset rapidness following application and uncaging of **13**. (**E**) Reduction in AP peak rate of rise following application and uncaging of **13**. (**F**) Increase in AP peak rate of fall following application and uncaging of **13**. (**G**) Reduction in maximal AP acceleration following application and uncaging of **13** (n = 4–7). (*p<0.05, **p<0.01, ***p<0.001 one-way ANOVA with Tukey's correction).

**Extended Data Figure 9.** Uncaging **13** against dissociated embryonic dorsal root ganglia cells affects AP onset rapidness, peak rate of rise, peak rate of fall, and maximal acceleration in a dose-dependent manner.

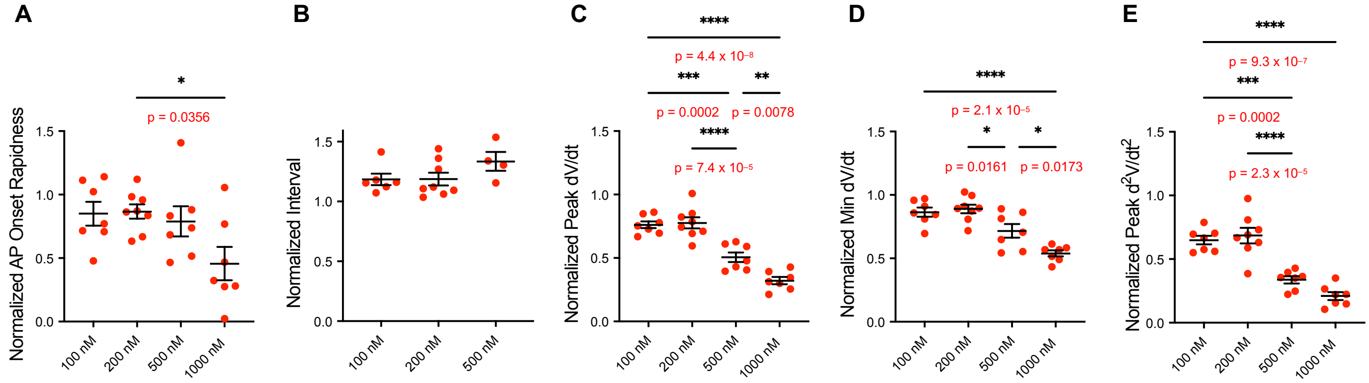


All data calculated from first AP in current step 2 following application of **13** and laser pulse. (**A**) Dose-dependent reduction in AP onset rapidness following application and uncaging of **13** at listed concentrations. (**B**) Interval between first and second AP trends upward (non-significant) with increasing concentration of uncaged **13**. (**C**) Dose-dependent reduction in AP peak rate of rise following application and uncaging of **13** at listed concentrations. Unlisted significant p-values: 200 nM vs 1000 nM, $p = 1.3 \times 10^{-8}$. (**D**) Dose-dependent reduction in AP peak rate of fall following uncaging of **13** at listed concentrations. Unlisted significant p-values: 200 nM vs 1000 nM, $p = 3.9 \times 10^{-6}$. (**E**) Dose-dependent reduction in maximal AP acceleration following application and uncaging of **13** at listed concentrations. Unlisted significant p-values: 200 nM vs 1000 nM, $p = 1.3 \times 10^{-7}$ (n = 4–8). (*$p<0.05$, **$p<0.01$, ***$p<0.001$, ****$p<0.0001$ one-way ANOVA with Tukey's correction).

**Extended Data Figure 10.** Application of 1.0 μM **13** to dissociated embryonic dorsal root ganglia produces a small reduction in AP amplitude prior to uncaging, but exerts no significant effect on any other measured AP parameter.

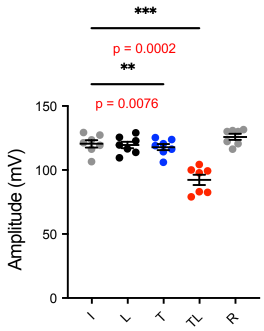


Action potential amplitudes calculated from first action potential in current step 2. Traces labeled: initial (I), laser applied (L), 1000 nM **13** applied (T), 1000 nM **13** and laser applied (TL), recovered (R) after wash-off (n = 7). (**$p<0.01$, ***$p<0.001$ one-way ANOVA with Tukey's correction).

**Extended Data Figure 11.** Uncaging of **13** exerts an outsized effect on AP firing rate, as compared to current block. Dorsal root ganglia neurons are less sensitive to **13** uncaging than hippocampal neurons.

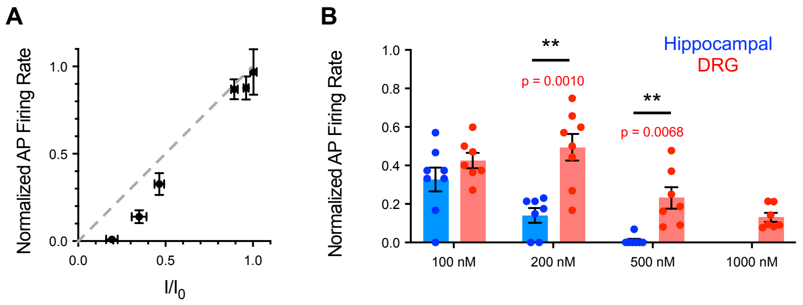


(A) Correlation between current block (x-axis) and AP firing rate (y-axis) of embryonic hippocampal neurons after uncaging of **13** (n = 5–8). (**p<0.01, ***p<0.001 one-way ANOVA with Tukey's correction). (**B**) Comparison between normalized AP firing rate in dissociated hippocampal and DRG neurons after uncaging **13** at specific concentrations (n = 7–8). (**p<0.01, multiple unpaired t-tests).

**Extended Data Figure 12.** Silencing of cortical network activity by STX-bpc **13**.

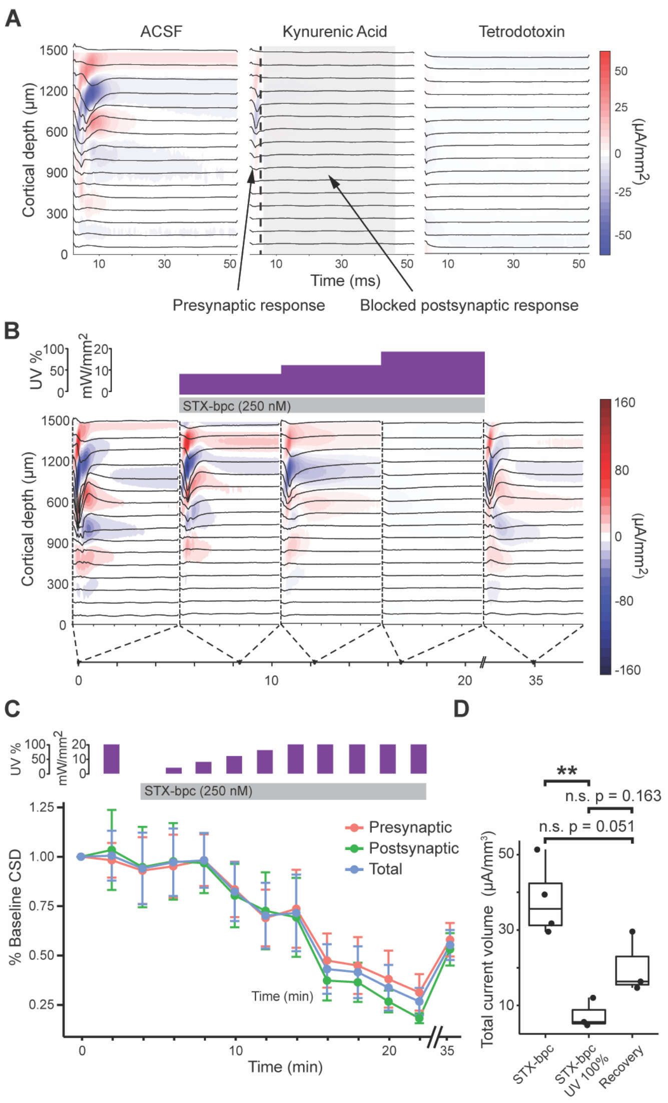


(**A**) Example of electrically evoked LFP and corresponding current source density (CSD) map showing components of the cortical network responses and their sensitivity to sequential blockade of glutamate receptors by kynurenic (1 mM) and blockade of $Na_V$s by TTX (1.0 µM). Application of kynurenic abolished postsynaptic activity (shown in grey) whereas presynaptic activity was abolished by TTX. The red (current source) and blue (current sink) shading represent the calculated CSD. The CSD values are represented by the scale on the right. (**B**) A representative example of cortical local field potential (LFP) and CSD responses at baseline and in the presence of 250 nM **13** following exposure to different intensities of UV light (as indicated by the purple bars at the top of the panel). A timeline is shown along the bottom of the panel, which indicates the point within the experiment when the recording was made. (**C**) Baseline normalized total CSD volume and CSD volume decomposed into presynaptic and postsynaptic components in the presence of UV light exposure and incrementing UV light intensities with **13** (as indicated by the purple bars at the top of the panel). A timeline is shown along the bottom of the panel, which indicates the point within the experiment when the recording was made. The shading relates to the amplitude of the CSD and corresponds to the scale bar shown on the right. (**D**) Quantification of the CSD following selected UV/**13** treatments. The summation of all current sink and source volumes within a 50 ms window after the stimulation are calculated for each slice. n = 3-4, ** $p<0.01$, (One-way ANOVA $p = 0.004$, df = 7, f = 13.6; Tukey Post-hoc). Each point represents a different slice, 1-2 slices per animal.

**Extended Data Video 1.** Uncaging of STX-bpc **13** in left brainstem biases sensory responses of larval zebrafish. Larval zebrafish swimming responses at 0.2x speed to three symmetrical vibration stimuli ('Stim.') presented before and three presented after uncaging of **13** in the left octavolateral hindbrain. The uncaging light is visible to the camera and elicits rightward tail bends. Note the unilateral deflections of the tail to stimuli after but not before uncaging. Periods between stimulus presentations are cropped and points along the tail are superimposed after posthoc tracking.

### 2. Extended Data Tables

**Extended Data Figure 1.** Comparison of Generation 1 photocaged STX apparent $IC_{50}$s pre- and post-uncaging.

| | Before uncaging | | After 5 x 5 ms 355 nm light | | Comparison | |
|---|---|---|---|---|---|---|
| | $IC_{50}$ (nM) | Hill Coefficient | $IC_{50}$ (nM) | Hill Coefficient | p-value | Fold Change |
| **1** | 14.4 ± 0.3 | –0.94 ± 0.02 | | | | |
| **2** | 17.3 ± 0.7 | –1.17 ± 0.05 | 18.9 ± 0.9 | –1.00 ± 0.05 | 0.0303 | 0.9 |
| **3** | 59.8 ± 3.3 | –1.35 ± 0.10 | 62.7 ± 3.5 | –1.14 ± 0.07 | 0.1873 | 1 |
| **4** | 132.1 ± 7.3 | –1.35 ± 0.10 | 77.4 ± 6.9 | –1.17 ± 0.13 | $<1 \times 10^{-6}$ | 1.7 |
| **5** | 114.2 ± 6.5 | –1.30 ± 0.10 | 120.0 ± 6.6 | –1.11 ± 0.07 | 0.2672 | 1 |
| **6** | 128.1 ± 7.2 | –1.25 ± 0.09 | 98.5 ± 4.6 | –1.11 ± 0.06 | 0.0017 | 1.3 |
| **7** | 121.9 ± 5.5 | –1.19 ± 0.06 | 56.0 ± 2.7 | –1.05 ± 0.06 | $<1 \times 10^{-6}$ | 2.2 |

Color-coded according to Figure 1 (note: compounds **5–7** are not shown in **Fig. 1**). See **Extended Data Figure 1** for structures.

**Extended Data Figure 2.** Comparison of Generation 2 photocaged STX apparent $IC_{50}$s pre- and post-uncaging.

| | Before uncaging | | After 5 x 5 ms 355 nm light | | Comparison | |
|---|---|---|---|---|---|---|
| | $IC_{50}$ (nM) | Hill Coefficient | $IC_{50}$ (nM) | Hill Coefficient | p-value | Fold Change |
| **1** | 14.4 ± 0.3 | –0.94 ± 0.02 | | | | |
| **8** | 67.6 ± 4.9 | –1.13 ± 0.09 | 45.7 ± 3.9 | –0.97 ± 0.08 | 0.0027 | 1.5 |
| **9** | 67.6 ± 3.6 | –1.23 ± 0.08 | 43.1 ± 2.0 | –1.09 ± 0.06 | $<1 \times 10^{-6}$ | 1.6 |
| **10** | 507.2 ± 39.0 | –1.38 ± 0.15 | 214.9 ± 20.2 | –1.12 ± 0.12 | $<1 \times 10^{-6}$ | 2.4 |
| **11** | 211.3 ± 25.7 | –0.93 ± 0.12 | 27.1 ± 4.6 | –0.84 ± 0.11 | $<1 \times 10^{-6}$ | 7.8 |
| **12** | 1024.9 ± 38.6 | –1.01 ± 0.04 | 25.1 ± 0.9 | –0.91 ± 0.03 | $<1 \times 10^{-6}$ | 40.8 |
| **13** | **3919.4 ± 172.6** | **–1.02 ± 0.05** | **21.3 ± 0.7** | **–0.84 ± 0.02** | $\mathbf{<1 \times 10^{-6}}$ | **184** |

Color-coded according to Figure 1. See **Extended Data Figure 1** for structures.

### 3. Chemical Procedures:

**General:** All reagents were obtained commercially unless otherwise noted. Organic solutions were concentrated under reduced pressure by rotary evaporation. Anhydrous $CH_2Cl_2$ and HPLC-grade $CH_3CN$ were obtained from commercial suppliers and used as is. *N,N*-Dimethylformamide (DMF) was passed through two columns of activated alumina prior to use. Triethylamine was distilled from calcium hydride.

Product purification was accomplished using forced-flow chromatography on Silicycle ultrapure silica gel (40–63 μm). Semi-preparative high-performance liquid chromatography (HPLC) was performed on a Varian ProStar model 210. Thin layer chromatography was performed with EM Science silica gel 60 $F_{254}$ plates (250 μm). Visualization of the developed chromatogram was accomplished by fluorescence quenching. High-resolution mass spectra were obtained from the Vincent Coates Foundation Mass Spectrometry Laboratory at Stanford University. Samples were analyzed with LC/ESI-MS by direct injection onto a Waters Acquity UPLC and Thermo Fisher Exactive mass spectrometer scanning m/z 100–2000. Methanol was used as the LC mobile phase at a flow rate of 0.175 mL/min. UV/Vis spectra were recorded on a Thermo NanoDrop One.

Saxitoxin derivatives were quantified by $^1$H NMR spectroscopy on a Varian Inova 600 MHz NMR instrument using distilled DMF as an internal standard. A relaxation delay (d1) of 20 s and an acquisition time (at) of 10 s were used for spectral acquisition. The concentration of the toxin derivative was determined by integration of $^1$H signals corresponding to toxin against those from a fixed concentration of the DMF standard.

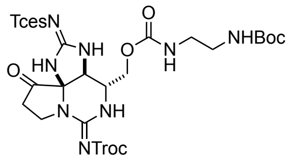


**Triprotected saxitoxin N21-ethylamine (14).** To an ice-cold solution of Tces- and Troc-protected decarbamoyl saxitoxin[1,2] (39 mg, 0.06 mmol) in 2.5 mL of THF was added 1,1'-carbonyldimidazole (25 mg, 0.157 mmol, 2.5 equiv). After 5 min, the reaction was warmed to room temperature and stirred for 4 h. Following this time, the reaction was quenched by the addition of 2.9 mL of saturated aqueous $NH_4Cl$. The biphasic mixture was stirred vigorously for 5 min then diluted with 10 mL of THF and transferred to a separatory funnel. The organic phase was collected and the aqueous phase was extracted with 3 x 10 mL of THF. The combined extracts were dried over $Na_2SO_4$, filtered, and concentrated under reduced pressure to a yellow foam. This material was dissolved in 2.5 mL of THF and to this solution was added *N*-Boc-1,2-ethylenediamine (50 μL, 0.314 mmol, 5.0 equiv). The pale yellow solution was stirred for 3 h following which time 2.9 mL of 0.5 M aqueous HCl was added. The biphasic mixture was stirred vigorously for 5 min then diluted with 10 mL of EtOAc and transferred to a separatory funnel. The organic phase was collected and the aqueous phase was extracted with 3 x 10 mL of EtOAc. The combined extracts were dried over $Na_2SO_4$, filtered, and concentrated under reduced pressure to a yellow foam. Purification of this material by chromatography on silica gel (gradient elution: 10:0→7:3 $CH_2Cl_2$/acetone) afforded carbamate **1** (30 mg, 59%) as a white powder. Boc, *tert*-butoxycarbonyl; Tces, 2,2,2-trichloroethoxysulfonyl; Troc, 2,2,2-trichloroethoxycarbonyl.

TLC (7:3 $CH_2Cl_2$/acetone): $R_f$ = 0.51

$^1$H NMR (600 MHz, $CD_3CN$) 8.84 (br s, 1H), 7.30 (br s, 1H), 7.06 (br s, 1H), 5.71 (br s, 1H), 5.42 (br s, 1H), 4.90 (d, *J* = 12.3 Hz, 1H), 4.69 (d, *J* = 12.3 Hz, 1H), 4.60 (s, 2H), 4.47 (s, 1H), 4.14–3.97 (m, 3H), 3.79–3.88 (m, 1H), 3.71–3.58 (m, 1H), 3.15–2.02 (m, 4H), 2.79–2.69 (m, 2H), 1.40 (s, 9H) ppm

$^{13}C$ NMR (125 MHz, $CD_3CN$) δ 206.0, 162.5, 161.6, 158.9, 157.1, 156.7, 96.7, 94.9, 79.4, 78.8, 75.6, 75.3, 63.8, 59.7, 54.4, 41.9, 41.5, 40.7, 33.7, 28.5 ppm

IR (thin film) ν 3310, 2929, 1706, 1586, 1530, 1391, 1230, 1179 $cm^{-1}$

HRMS (ESI$^+$): $[MH]^+$ calcd for $C_{22}H_{30}Cl_6N_8O_{10}S$, 809.0010; found, 809.0004

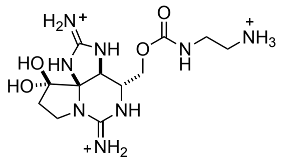

**Saxitoxin N21-ethylamine (1).** To a solution of carbamate **14** (15 mg, 18.5 μmol) in 4.6 mL of a 3:1 $MeOH/H_2O$ was added 212 μL of trifluoroacetic acid. The mixture was stirred for 30 min then $PdCl_2$ (1.13 mg, 9.2 μmol, 0.5 equiv) was added. The suspension was sparged with a gentle stream of $N_2$ for 5 min and then $H_2$ for 5 min. The flask was fitted with a balloon of $H_2$ (1 atm) and the contents stirred for 3.5 h. Following this time, the mixture was filtered through a Fisher 0.2 μm PTFE syringe filter. The flask and filter were rinsed with 4 mL of MeOH and 7.4 mL of 1.0 N aqueous HCl, and the combined filtrates were concentrated under reduced pressure. The isolated residue was redissolved in 2.3 mL of 1.0 N aqueous HCl and the mixture was stirred for 1 h, then frozen and lyophilized to remove all volatiles. The product was purified by reversed-phase HPLC (Silicycle SiliaChrom AQ C18, 5 μm, 10 x 250 mm column, eluting with a gradient flow of 10→20% $CH_3CN$ in 10 mM aqueous $C_3F_7CO_2H$ over 60 min, 214 nm UV detection). At a flow rate of 4 mL/min, **1** had a retention time of 22–30 min and was isolated as a white powder following lyophilization (6.13 μmol, 33%, $^1H$ NMR quantitation). This compound has been characterized previously.[3,4]

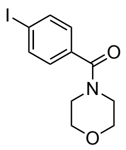

**(4-Iodophenyl)(morpholino)methanone (15).** To a solution of 4-iodobenzoic acid (248 mg, 1.0 mmol) in 6.0 mL of a 5:1 mixture of $THF/CH_2Cl_2$ was added successively 1-ethyl-3-(3-dimethylaminopropyl)carbodiimide (230 mg, 1.2 mmol, 1.2 equiv), 1-hyroxybenzotriazole hydrate (185 mg, 1.2 mmol, 1.2 equiv), and morpholine (173 μL, 2.0 mmol, 2.0 equiv). The reaction mixture was stirred for 24 h and then concentrated under reduced pressure. Purification of this material by chromatography on silica gel (gradient elution: 10:0→9:1 $CH_2Cl_2$/acetone) afforded **14** (286 mg, 90%) as a white foam.

TLC (9:1 $CH_2Cl_2$/acetone): $R_f = 0.66$

$^1H$ NMR (500 MHz, $CDCl_3$) δ 7.75 (d, $J$ = 8.3 Hz, 2H), 7.13 (d, $J$ = 8.3 Hz, 2H), 3.71 (br s, 4H), 3.64 (br s, 2H), 3.43 (br s, 2H) ppm

$^{13}C$ NMR (125 MHz, $CDCl_3$) δ 169.6, 137.8, 134.8, 128.9, 96.2, 66.9, 48.4, 42.6 ppm

IR (thin film) ν 2854, 1640, 1587, 1457, 1254, 1112 $cm^{-1}$

HRMS (ESI$^+$): $[MH]^+$ calcd for $C_{11}H_{12}INO_2$, 317.9985; found, 317.9972

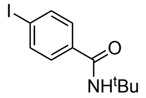

***N*-(*tert*-Butyl)-4-iodobenzamide (16).** This compound was prepared in an analogous manner to **15** starting from 4-iodobenzoic acid (740 mg, 4.8 mmol) and substituting *tert*-butylamine for morpholine. Purification

by chromatography on silica gel (gradient elution: 100:0→97:3 $CH_2Cl_2$/MeOH) afforded **15** (1.06 g, 87%) as a white powder.

TLC (7:3 hexanes/EtOAc): $R_f = 0.62$

$^1$H NMR (500 MHz, $CDCl_3$) δ 7.63 (d, $J$ = 8.5 Hz, 2H), 7.36 (d, $J$ = 8.5 Hz, 2H), 6.17 (br s, 1H), 1.40 (s, 9H) ppm

$^{13}$C NMR (125 MHz, $CDCl_3$) δ 166.1, 137.4, 135.3, 128.4, 97.8, 51.7, 28.8 ppm

IR (thin film) ν 3317, 1634, 1539, 1449, 1320, 1216, 1006 $cm^{-1}$

HRMS (ESI$^+$): $[MH]^+$ calcd for $C_{11}H_{14}INO$, 304.0193; found, 304.0181

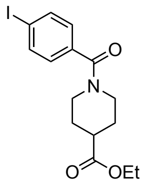


**Ethyl 1-(4-iodobenzoyl)piperidine-4-carboxylate (17).** This compound was prepared in an analogous manner to **15** starting from 4-iodobenzoic acid (500 mg, 2.0 mmol) and substituting ethyl isonipecotate for morpholine. Purification by chromatography on silica gel (gradient elution: 5:0→4:1 $CH_2Cl_2$/acetone) afforded **17** (745 mg, 94%) as a clear oil.

TLC (9:1 $CH_2Cl_2$/acetone): $R_f = 0.73$

$^1$H NMR (500 MHz, $CDCl_3$) δ 7.65 (d, $J$ = 8.3 Hz, 2H), 7.05 (d, $J$ = 8.3 Hz, 2H), 4.38 (br s, 1H), 4.06 (q, $J$ = 7.1 Hz, 2H), 3.60 (br s, 1H), 2.96 (br s, 2H), 2.48 (tt, $J$ = 10.7, 4.0 Hz, 1H), 1.91 (br s, 1H), 1.77 (br s, 1H), 1.61 (br s, 2H), 1.16 (t, $J$ = 7.1 Hz, 3H) ppm

$^{13}$C NMR (125 MHz, $CDCl_3$) δ 173.8, 169.2, 137.5, 135.3, 128.5, 95.6, 60.5, 46.8, 41.4, 40.7, 28.3, 27.7, 14.1 ppm

IR (thin film) ν 2954, 2860, 1728, 1630, 1437, 1315, 1181, 1146, 1041, 1003 $cm^{-1}$

HRMS (ESI$^+$): $[MH]^+$ calcd for $C_{15}H_{18}INO_3$, 388.0404; found, 388.0393

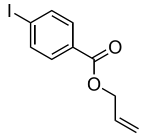


**Allyl 4-iodobenzoate (18).** To a solution of 4-iodobenzoic acid (850 mg, 3.4 mmol) in 34 mL of *N,N*-dimethylformamide was added *i*-$Pr_2NEt$ base (720 µL, 4.1 mmol, 1.2 equiv) and allyl bromide (360 µL, 4.1 mmol, 1.2 equiv). The reaction mixture was stirred for 16 h then diluted with 200 mL of $CH_2Cl_2$ and transferred to a separatory funnel containing 200 mL of saturated aqueous $NaHCO_3$. The organic layer was collected and the aqueous layer was extracted with 2 x 200 mL of $CH_2Cl_2$. The organic fraction was dried over $Na_2SO_4$, filtered, and concentrated under reduced pressure to a clear oil. Purification of this material by chromatography on silica gel (gradient elution: 10:0→8:2 hexanes/EtOAc) afforded **18** (780 mg, 79%) as a clear oil.

TLC (9:1 hexanes/EtOAc): $R_f = 0.65$

$^1$H NMR (500 MHz, $CDCl_3$) δ 7.77–7.70 (m, 4H), 6.04–5.95 (m, 1H), 5.37 (dd, $J$ = 17.2, 1.5 Hz, 1H), 5.26 (dd, $J$ = 10.4, 1.3 Hz, 1H), 4.48 (d, $J$ = 5.6 Hz, 2H) ppm

$^{13}$C NMR (125 MHz, $CDCl_3$) δ 165.6, 137.7, 132.0, 131.0, 129.6, 118.5, 100.9, 65.7 ppm

IR (thin film) ν 2943, 1724, 1587, 1393, 1267, 1177, 1102, 1008 cm$^{-1}$

HRMS (ESI$^+$): [MH]$^+$ calcd for $C_{10}H_9IO_2$, 288.9720; found, 288.9724

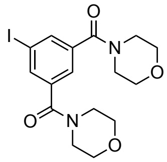

**(5-Iodo-1,3-phenylene)bis(morpholinomethanone) (19).** To a solution of 5-iodoisophthalic acid (500 mg, 1.7 mmol) in 12 mL of a 5:1 mixture of THF/$CH_2Cl_2$ was added successively 1-ethyl-3-(3-dimethylaminopropyl)carbodiimide (788 mg, 4.1 mmol, 2.4 equiv), 1-hyroxybenzotriazole hydrate (633 mg, 4.1 mmol, 2.4 equiv), and morpholine (591 µL, 6.8 mmol, 4.0 equiv). The reaction mixture was stirred for 24 h and then concentrated under reduced pressure. Purification of this material by chromatography on silica gel (gradient elution: 5:0→3:2 $CH_2Cl_2$/acetone) afforded **19** (620 mg, 84%) as a white foam.

TLC (4:1 $CH_2Cl_2$/acetone): $R_f = 0.42$

$^1$H NMR (500 MHz, $CDCl_3$) δ 7.70 (d, $J$ = 1.5 Hz, 2H), 7.27 (t, $J$ = 1.5 Hz, 1H), 3.64 (br s, 8H), 3.52 (br s, 4H), 3.31 (br s, 3.31) ppm

$^{13}$C NMR (125 MHz, $CDCl_3$) δ 167.3, 137.5, 136.9, 124.5, 94.2, 66.5 (2), 48.0, 42.5 ppm

IR (thin film) ν 2857, 1634, 1439, 1410, 1274, 1114, 1036 cm$^{-1}$

HRMS (ESI$^+$): [MH]$^+$ calcd for $C_{16}H_{19}IN_2O_4$, 431.0462; found, 431.0451

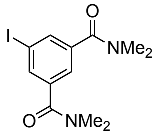

**5-Iodo-$N^1$,$N^1$,$N^3$,$N^3$-tetramethylisophthalamide (20).** This compound was prepared in an analogous manner to **19** starting from 5-iodoisophthalic acid (500 mg, 1.7 mmol) and substituting 2.0 M dimethylamine in THF (2.5 mL, 6.9 mmol, 4.0 equiv) for morpholine. Purification by chromatography on silica gel (gradient elution: 5:0→2:3 $CH_2Cl_2$/acetone) afforded **20** (445 mg, 75%) as a white powder.

TLC (4:1 $CH_2Cl_2$/acetone): $R_f = 0.44$

$^1$H NMR (500 MHz, $CDCl_3$) δ 7.77 (d, $J$ = 1.4 Hz, 2H), 7.37 (t, $J$ = 1.5 Hz, 1H), 3.05 (br s, 6H), 2.95 (br s, 6H) ppm

$^{13}$C NMR (125 MHz, $CDCl_3$) δ 169.0, 138.4, 136.9, 124.8, 124.7, 94.1, 39.6, 35.4 ppm

IR (thin film) ν 3482, 2932, 2360, 1635, 1506, 1394, 1270, 1187, 1103 cm$^{-1}$

HRMS (ESI$^+$): [MH]$^+$ calcd for $C_{12}H_{15}IN_2O_2$, 347.0251; found, 347.0239

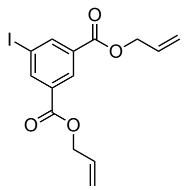

**Diallyl 5-iodoisophthalate (21).** This compound was prepared in an analogous manner to **18** starting from 5-iodoisophthalic acid (1.0 g, 3.4 mmol) using 2.4 equivalents of *i*-$Pr_2NEt$ and allyl bromide. Purification by chromatography on silica gel (gradient elution: 10:0→8:2 $CH_2Cl_2$/acetone) afforded **21** (1.04 g, 81%) as a clear oil.

TLC (9:1 hexanes/EtOAc): $R_f = 0.55$

$^1$H NMR (500 MHz, $CDCl_3$) δ 8.55 (t, *J* = 1.6 Hz, 1H), 8.45 (d, *J* = 1.5 Hz, 2H), 6.03–5.93 (m, 2H), 5.37 (dd, *J* = 17.1, 1.5 Hz, 2H), 5.26 (dd, *J* = 10.4, 1.3 Hz, 2H), 4.78 (d, *J* = 5.8 Hz, 4H) ppm

$^{13}$C NMR (125 MHz, $CDCl_3$) δ 164.0, 142.6, 132.4, 131.8, 130.1, 119.2, 93.6, 66.4 ppm

IR (thin film) ν 3081, 1727, 1361, 1300, 1232, 1137, 985, 934 cm$^{-1}$

HRMS (ESI$^+$): [MH]$^+$ calcd for $C_{14}H_{13}IO_4$, 372.9931; found, 372.9933

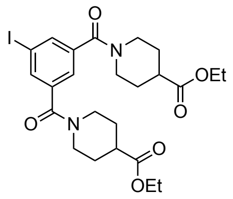


**Diethyl 1,1'-(5-iodoisophthaloyl)bis(piperidine-4-carboxylate) (22).** This compound was prepared in an analogous manner to **19** starting from 5-iodoisophthalic acid (876 mg, 3.0 mmol) and substituting ethyl isonipecotate for morpholine. Purification by chromatography on silica gel (gradient elution: 5:0→4:1 $CH_2Cl_2$/acetone) afforded **22** (1.27 g, 75%) as a clear oil.

TLC (4:1 $CH_2Cl_2$/acetone): $R_f = 0.59$

$^1$H NMR (500 MHz, $CDCl_3$) δ 7.67 (d, *J* = 1.5 Hz, 2H), 7.22 (t, *J* = 1.5 Hz, 1H), 4.34 (br s, 2H), 4.04 (q, *J* = 7.1 Hz, 4H), 3.55 (br s, 2H), 2.97 (br s, 4H), 2.46 (tt, *J* = 10.5, 4.0 Hz, 2H), 1.91 (br s, 2H), 1.77 (br s, 2H), 1.64 (br s, 2H), 1.55 (br s, 2H), 1.15 (t, *J* = 7.1 Hz, 6H) ppm

$^{13}$C NMR (125 MHz, $CDCl_3$) δ 173.7, 167.4, 138.0, 136.5, 123.9, 94.3, 60.5, 46.8, 41.4, 40.6, 28.3, 27.6, 14.1 ppm

IR (thin film) ν 2955, 2861, 1728, 1635, 1446, 1410, 1316, 1273, 1182, 1041 cm$^{-1}$

HRMS (ESI$^+$): [MH]$^+$ calcd for $C_{24}H_{31}IN_2O_6$, 571.1300; found, 571.1294

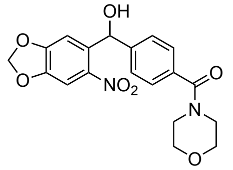


**(4-(Hydroxy(6-nitrobenzo[*d*][1,3]dioxol-5-yl)methyl)phenyl)(morpholino)methanone (23).** To a –40 °C solution of *i*-PrMgCl•LiCl (1.5 mL of 1.3 M in THF, 1.95 mmol, 1.3 equiv) was added dropwise a solution of **15** (476 mg, 1.5 mmol) in 7.5 mL of THF. The reaction mixture was stirred for 2 h at –40 °C. Following this time, a solution of 6-nitropiperonal (293 mg, 1.5 mmol) in 6.0 mL THF was added dropwise. The reaction mixture was warmed to room temperature over 2 h and quenched by the addition of 5 mL of saturated aqueous $NH_4Cl$. The solution was transferred to a separatory funnel containing 10 mL of EtOAc. The organic layer was collected and the aqueous layer was extracted with 2 x 10 mL of EtOAc. The organic fractions were combined, dried over $Na_2SO_4$, filtered, and concentrated under reduced pressure. Purification of this material by chromatography on silica gel (gradient elution: 10:0→9:1 $CH_2Cl_2$/acetone) afforded **23** (306 mg, 53%) as a light yellow powder.

TLC (8:2 $CH_2Cl_2$/acetone): $R_f = 0.39$

$^1$H NMR (500 MHz, $CDCl_3$) δ 7.49 (s, 1H), 7.38 (d, *J* = 8.1 Hz, 2H), 7.33 (d, *J* = 8.3 Hz, 2H), 7.14 (s, 1H), 6.41 (s, 1H), 6.11 (s, 2H), 3.73 (br s, 4H), 3.62 (br s, 2H), 3.42 (br s, 2H), 2.81 (br s, 1H) ppm

$^{13}C$ NMR (125 MHz, $CDCl_3$) δ 170.3, 152.5, 147.5, 143.9, 142.2, 136.2, 134.7, 127.4, 127.2, 108.4, 105.6, 103.3, 70.8, 67.0 (2), 48.4, 42.9 ppm

IR (thin film) ν 3374, 2917, 1617, 1521, 1483, 1436, 1333, 1259, 1114, 1035 $cm^{-1}$

HRMS ($ESI^+$): $[MH]^+$ calcd for $C_{19}H_{18}N_2O_7$, 387.1187; found, 387.1176

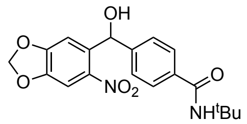


***N*-(*tert*-Butyl)-4-(hydroxy(6-nitrobenzo[*d*][1,3]dioxol-5-yl)methyl)benzamide (24).** This compound was prepared in an analogous manner to **23** starting from **16** (305 mg, 0.82 mmol). Purification by chromatography on silica gel (gradient elution: 10:0→7:3 hexanes/EtOAc) afforded **24** (200 mg, 48%) as a light yellow powder.

TLC (7:3 hexanes/EtOAc): $R_f = 0.11$

$^1H$ NMR (500 MHz, $CDCl_3$) δ 7.64 (d, $J$ = 8.4 Hz, 2H), 7.49 (s, 1H), 7.38 (d, $J$ = 7.9 Hz, 2H), 7.08 (s, 1H), 6.43 (s, 1H), 6.11 (d, $J$ = 1.2 Hz, 2H), 5.90 (br s, 1H), 1.46 (s, 9H) ppm

$^{13}C$ NMR (125 MHz, $CDCl_3$) δ 166.9, 152.4, 147.5, 145.0, 142.3, 136.1, 135.4, 127.1, 127.0, 108.5, 105.7, 103.2, 71.0, 51.9, 29.0 ppm

IR (thin film) ν 3335, 2922, 1636, 1521, 1483, 1332, 1259, 1036 $cm^{-1}$

HRMS ($ESI^+$): $[MH]^+$ calcd for $C_{19}H_{20}N_2O_6$, 373.1394; found, 373.1383

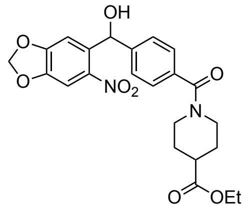


**Ethyl 1-(4-(hydroxy(6-nitrobenzo[*d*][1,3]dioxol-5-yl)methyl)benzoyl)piperidine-4-carboxylate (25).** This compound was prepared in an analogous manner to **23** starting from **17** (581 mg, 1.5 mmol), but at a higher reaction concentration using 3.0 mL of THF to dissolve both **17** and 6-nitropiperonal. Purification by chromatography on silica gel (gradient elution: 10:0→9:1 $CH_2Cl_2$/acetone) afforded **25** (295 mg, 43%) as a yellow foam.

TLC (9:1 $CH_2Cl_2$/acetone): $R_f = 0.38$

$^1H$ NMR (500 MHz, $CDCl_3$) δ 7.41 (s, 1H), 7.28 (d, $J$ = 8.2 Hz, 2H), 7.21 (d, $J$ = 8.2 Hz, 2H), 7.19 (s, 1H), 6.34 (s, 1H), 6.06 (d, $J$ = 1.2 Hz, 2H), 4.39 (br s, 1H), 4.25 (br s, 1H), 4.10 (q, $J$ = 7.1 Hz, 2H), 3.65 (br s, 1H), 2.98 (br s, 2H), 2.51 (tt, $J$ = 10.7, 4.0 Hz, 1H), 1.94 (br s, 1H), 1.80 (br s, 1H), 1.64 (br s, 2H), 1.21 (t, $J$ = 7.1 Hz, 3H) ppm

$^{13}C$ NMR (125 MHz, $CDCl_3$) δ 174.2, 170.3, 152.3, 147.1, 144.2, 141.8, 136.7, 134.7, 127.1, 126.9, 108.2, 105.2, 103.1, 70.3, 60.7, 47.1, 41.7, 40.9, 28.5, 28.0, 14.2 ppm

IR (thin film) ν 3372, 2927, 1728, 1616, 1522, 1483, 1447, 1365, 1332, 1258, 1183, 1148, 1038, 930 $cm^{-1}$

HRMS ($ESI^+$): $[MH]^+$ calcd for $C_{23}H_{24}N_2O_8$, 457.1605; found, 457.1592

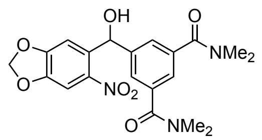

**5-(Hydroxy(6-nitrobenzo[*d*][1,3]dioxol-5-yl)methyl)-*N*$^1$,*N*$^1$,*N*$^3$,*N*$^3$-tetramethylisophthalamide (26).** This compound was prepared in an analogous manner to **23** starting from **20** (242 mg, 0.7 mmol), but at a higher reaction concentration using 1.5 mL of THF to dissolve **20** and 2 mL of THF to dissolve 6-nitropiperonal. Purification by chromatography on silica gel (gradient elution: 2:0→1:1 $CH_2Cl_2$/acetone) afforded **26** (216 mg, 74%) as a yellow foam.

TLC (4:1 $CH_2Cl_2$/acetone): $R_f = 0.19$

$^1$H NMR (500 MHz, $CDCl_3$) δ 7.40 (s, 1H), 7.34 (d, $J$ = 1.5 Hz, 2H), 7.28 (s, 1H), 7.26 (d, $J$ = 1.6 Hz, 1H), 6.35 (s, 1H), 6.08 (d, $J$ = 1.2 Hz, 2H), 4.88 (br s, 1H), 3.02 (br s, 6H), 2.88 (br s, 6H) ppm

$^{13}$C NMR (125 MHz, $CDCl_3$) δ 170.7, 152.5, 147.3, 143.6, 141.7, 136.5, 136.1, 127.2, 127.0, 125.0, 124.8, 108.2, 108.0, 105.3, 105.1, 103.3, 103.1, 102.9, 70.3, 70.1, 39.8, 39.6, 35.6, 35.4 ppm

IR (thin film) ν 3358, 2360, 1617, 1506, 1484, 1396, 1334, 1259, 1034 cm$^{-1}$

HRMS (ESI$^+$): [MH]$^+$ calcd for $C_{20}H_{21}N_3O_7$, 416.1452; found, 416.1441

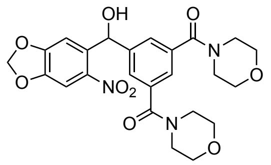

**(5-(Hydroxy(6-nitrobenzo[*d*][1,3]dioxol-5-yl)methyl)-1,3-phenylene)bis(morpholinomethanone) (27).** This compound was prepared in an analogous manner to **26** starting from **19** (300 mg, 0.7 mmol). Purification by chromatography on silica gel (gradient elution: 2:0→1:1 $CH_2Cl_2$/acetone) afforded **27** (235 mg, 67%) as a yellow foam.

TLC (4:1 $CH_2Cl_2$/acetone): $R_f = 0.11$

$^1$H NMR (500 MHz, $CDCl_3$) δ 7.42 (s, 1H), 7.36 (d, $J$ = 1.5 Hz, 2H), 7.29 (t, $J$ = 1.6 Hz, 1H), 7.27 (s, 1H), 6.34 (s, 1H), 6.11 (d, $J$ = 1.1 Hz, 2H), 4.17 (br s, 1H), 3.72 (br s, 8H), 3.57 (br s, 4H), 3.33 (br s, 4H) ppm

$^{13}$C NMR (125 MHz, $CDCl_3$) δ 169.4, 152.6, 147.5, 143.7, 141.9, 136.0, 135.7, 127.4, 125.1, 108.0, 105.4, 103.3, 70.3, 66.8 (2), 48.3, 42.7 ppm

IR (thin film) ν 3384, 2969, 2916, 2859, 1628, 1521, 1483, 1426, 1334, 1252, 1115, 1036 cm$^{-1}$

HRMS (ESI$^+$): [MH]$^+$ calcd for $C_{24}H_{25}N_3O_9$, 500.1664; found, 500.1651

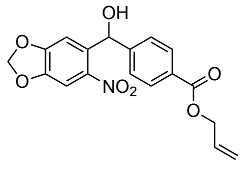

**Allyl 4-(hydroxy(6-nitrobenzo[*d*][1,3]dioxol-5-yl)methyl)benzoate (28).** This compound was prepared in an analogous manner to **26** starting from **18** (387 mg, 1.3 mmol). Purification by chromatography on silica gel (gradient elution: 20:0→19:1 $CH_2Cl_2$/acetone) afforded **28** (181 mg, 38%) as a yellow solid.

TLC (19:1 $CH_2Cl_2$/acetone): $R_f = 0.67$

$^{1}H$ NMR (500 MHz, $CDCl_3$) δ 8.01 (d, *J* = 8.5 Hz, 2H), 7.48 (s, 1H), 7.43 (d, *J* = 8.4 Hz, 2H), 7.05 (s, 1H), 6.46 (s, 1H), 6.11 (d, *J* = 1.2 Hz, 2H), 6.07–5.97 (m, 1H), 5.40 (dd, *J* = 17.2, 1.5 Hz, 1H), 5.28 (dd, *J* = 10.5, 1.3 Hz, 1H), 4.80 (d, *J* = 5.6 Hz, 2H), 3.12 (br s, 1H) ppm

$^{13}C$ NMR (125 MHz, $CDCl_3$) δ 166.1, 152.5, 147.6, 146.7, 142.5, 135.7, 132.3, 130.0, 129.8, 126.9, 118.4, 108.5, 105.7, 103.3, 71.0, 65.7 ppm

IR (thin film) ν 3444, 2917, 1718, 1611, 1521, 1504, 1483, 1421, 1361, 1332, 1261, 1118, 1035, 928 $cm^{-1}$

HRMS ($ESI^+$): $[MH]^+$ calcd for $C_{18}H_{15}NO_7$, 358.0921; found, 358.0908

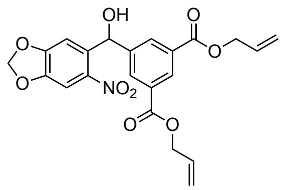


**Diallyl 5-(hydroxy(6-nitrobenzo[*d*][1,3]dioxol-5-yl)methyl)isophthalate (29).** This compound was prepared in an analogous manner to **26** starting from **21** (500 mg, 1.3 mmol). Purification by chromatography on silica gel (gradient elution: 20:0→19:1 $CH_2Cl_2$/acetone) afforded **29** (217 mg, 37%) as a yellow foam.

TLC (19:1 $CH_2Cl_2$/acetone): $R_f = 0.68$

$^{1}H$ NMR (500 MHz, $CDCl_3$) δ 8.56 (t, *J* = 1.6 Hz, 1H), 8.20 (d, *J* = 1.6 Hz, 2H), 7.46 (s, 1H), 7.10 (s, 1H), 6.48 (s, 1H), 6.10 (d, *J* = 1.3 Hz, 2H), 6.01 (dddd, *J* = 17.3, 10.3, 5.7, 1.2 Hz, 2H), 5.39 (dd, *J* = 17.2, 1.5 Hz, 2H), 5.28 (dd, *J* = 10.4, 1.3 Hz, 2H), 4.80 (d, *J* = 5.7 Hz, 4H), 3.56 (br s, 1H) ppm

$^{13}C$ NMR (125 MHz, $CDCl_3$) δ 165.3, 152.5, 147.6, 143.1, 142.1, 135.4, 132.4, 131.9, 130.8, 130.1, 118.7, 108.2, 105.5, 103.2, 70.5, 66.1 ppm

IR (thin film) ν 3483, 3086, 2916, 1724, 1523, 1484, 1333, 1260, 1235, 1183, 1036, 986 $cm^{-1}$

HRMS ($ESI^+$): $[MNa]^+$ calcd for $C_{22}H_{19}NO_9$, 464.0952; found, 464.0939

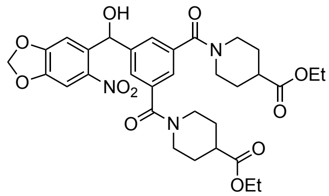


**Diethyl 1,1'-(5-(hydroxy(6-nitrobenzo[*d*][1,3]dioxol-5-yl)methyl)isophthaloyl)bis(piperidine-4-carboxylate) (30).** This compound was prepared in an analogous manner to **26** starting from **22** (570 mg, 1.0 mmol) using 1.2 equiv of 6-nitropiperonal. Purification by chromatography on silica gel (gradient elution: 2:0→1:1 $CH_2Cl_2$/acetone) afforded **30** (227 mg, 35%) as a light yellow foam.

TLC (4:1 $CH_2Cl_2$/acetone): $R_f = 0.15$

$^{1}H$ NMR (500 MHz, $CDCl_3$) δ 7.49 (s, 1H), 7.43 (s, 2H), 7.32 (s, 1H), 7.19 (s, 1H), 6.43 (d, *J* = 3.9 Hz, 1H), 6.14 (s, 2H), 4.48 (br s, 2H), 4.16 (q, *J* = 7.1 Hz, 4H), 3.64 (br s, 2H), 3.09 (d, *J* = 4.5 Hz, 1H), 3.04 (br s, 4H), 2.56 (tt, *J* = 10.1, 5.3 Hz, 2H), 2.01 (br s, 2H), 1.84 (br s, 2H), 1.76 (br s, 2H), 1.65 (br s, 2H), 1.27 (t, *J* = 7.1 Hz, 6H) ppm

$^{13}C$ NMR (125 MHz, $CDCl_3$) δ 174.1, 169.4, 152.6, 147.5, 143.5, 142.0, 136.3, 136.1, 126.9, 124.6, 108.1, 105.4, 103.3, 70.5, 60.8, 47.1, 41.7, 41.0, 31.0, 28.6, 27.9 ppm

IR (thin film) ν 3346, 2930, 2863, 2244, 1727, 1620, 1521, 1482, 1448, 1317, 1257, 1179, 1039, 918 $cm^{-1}$

HRMS ($ESI^+$): $[MH]^+$ calcd for $C_{32}H_{37}N_3O_{11}$, 640.2501; found, 640.2486

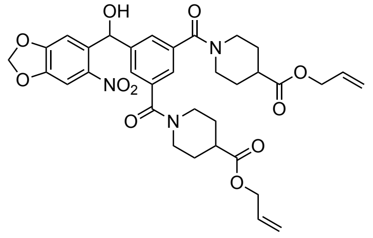

**Diallyl 1,1'-(5-(hydroxy(6-nitrobenzo[*d*][1,3]dioxol-5-yl)methyl)isophthaloyl)bis(piperidine-4-carboxylate) (31).** To a solution of **30** (117 mg, 0.18 mmol) in 1.8 mL of a 1:1 1,4-dioxane/water mixture was added LiOH•$H_2O$ (30 mg, 0.73 mmol, 4.0 equiv). The reaction mixture was stirred for 15 min then slowly diluted with 9 mL of 1.0 N HCl and transferred to a separatory funnel containing 10 mL of EtOAc. The organic layer was collected and the aqueous layer was extracted with 2 x 10 mL of EtOAc. The organic fractions were combined, dried over $Na_2SO_4$, filtered, and concentrated under reduced pressure to a white foam. This material was redissolved in 1.8 mL of DMF and to this solution were added successively *i*-$Pr_2NEt$ (80 µL, 0.46 mmol, 2.5 equiv) and allyl bromide (40 µL, 0.46 mmol, 2.5 equiv). The reaction mixture was stirred for 18 h. Following this time, the reaction was quenched by the addition of 50 mL of saturated aqueous $NaHCO_3$ and transferred to a separatory funnel containing 50 mL of EtOAc. The organic layer was collected and the aqueous layer was extracted with 2 x 50 mL of EtOAc. The organic fractions were combined, dried over $Na_2SO_4$, filtered, and concentrated under reduced pressure to a yellow oil. Purification of this material by chromatography on silica gel (gradient elution: 1:0→0:1 $CH_2Cl_2$/acetone) afforded **31** (67 mg, 54%) as a light yellow foam.

TLC (8:2 $CH_2Cl_2$/acetone): $R_f = 0.19$

$^1$H NMR (500 MHz, $CDCl_3$) δ 7.42 (s, 1H), 7.35 (s, 2H), 7.25 (s, 2H), 6.37 (d, *J* = 4.4 Hz, 1H), 6.10 (s, 2H), 5.88 (ddt, *J* = 16.2, 10.6, 5.6 Hz, 2H), 5.29 (dd, *J* = 17.2, 1.6 Hz, 2H), 5.22 (dd, *J* = 10.5, 1.4 Hz, 2H), 4.57 (dd, *J* = 5.8, 1.6 Hz, 4H), 4.45 (s, 1H), 4.42 (br s, 2H), 3.60 (br s, 2H), 4.99 (br s, 4H), 2.57 (ddt, *J* = 10.8, 8.5, 3.9 Hz, 2H), 2.00 (br s, 2H), 1.81 (br s, 2H), 1.73 (br s, 2H), 1.63 (br s, 1H), 1.56 (br s, 1H) ppm

$^{13}$C NMR (125 MHz, $CDCl_3$) δ 173.7, 169.3, 152.5, 147.5, 143.6, 141.9, 136.3, 136.2, 132.0, 127.0, 124.6, 118.5, 108.1, 105.4, 103.2, 70.4, 65.4, 47.1, 41.7, 41.0, 28.6, 27.9 ppm

IR (thin film) ν 3347, 2930, 2863, 2246, 1732, 1622, 1521, 1483, 1317, 1259, 1175, 1036 $cm^{-1}$

HRMS (ESI$^+$): $[MH]^+$ calcd for $C_{34}H_{37}N_3O_{11}$, 664.2501 found, 664.2505

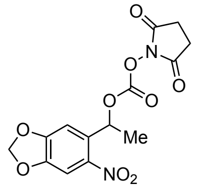

**(1-(6-Nitrobenzo[*d*][1,3]dioxol-5-yl)ethyl) succinimid-N-yl carbonate (32).** To a solution of 1-(6-nitro-1,3-benzodioxol-5-yl)ethanol (15 mg, 70 µmol) in 355 µL of *N,N*-dimethylformamide was added *N,N'*-disuccinimidyl carbonate (36 mg, 140 µmol, 2.0 equiv) and $Et_3N$ (19.8 µL, 140 µmol, 2.0 equiv). The reaction mixture was stirred for 22 h then diluted with 5 mL of $CH_2Cl_2$ and transferred to a separatory funnel containing 5 mL of saturated aqueous $NH_4Cl$. The organic layer was collected and the aqueous layer was extracted with 3 x 5 mL of $CH_2Cl_2$. The organic fractions were combined, dried over $Na_2SO_4$, filtered, and concentrated under reduced pressure. Purification of this material by chromatography on silica gel (gradient elution: 1:0→0:1 hexanes/EtOAc) afforded **32** (29 mg, 99%) as a yellow powder.

TLC (1:1 hexanes/EtOAc): $R_f = 0.32$

$^1$H NMR (500 MHz, $CD_3CN$) δ 7.49 (s, 1H), 7.16 (s, 1H), 6.30 (q, *J* = 6.4 Hz, 1H), 6.18 (s, 2H), 2.74 (s, 4H), 1.71 (d, *J* = 6.4 Hz, 3H) ppm

$^{13}C$ NMR (125 MHz, $CD_3CN$) δ 170.5, 153.8, 151.8, 149.1, 143.0, 133.0, 106.5, 105.8, 105.1, 76.8, 26.2, 21.7 ppm

IR (thin film) ν 1789, 1742, 1487, 1506, 1340, 1261, 1235, 1065, 1034 $cm^{-1}$

HRMS ($ESI^+$): $[MNa]^+$ calcd for $C_{14}H_{12}N_2O_9$, 375.0435; found, 375.0423

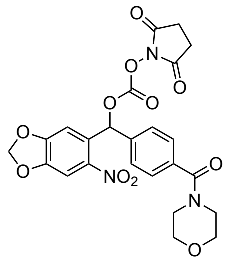


**((4-(Morpholine-4-carbonyl)phenyl)(6-nitrobenzo[*d*][1,3]dioxol-5-yl)methyl) succinimid-N-yl carbonate (33).** To a solution of **23** (50 mg, 0.13 mmol) in 3.2 mL of *N,N*-dimethylformamide was added *N,N'*-disuccinimidyl carbonate (66 mg, 0.26 mmol, 2.0 equiv) and $Et_3N$ (18 µL, 0.13 mmol). The reaction mixture was stirred for 22 h then diluted with 20 mL of EtOAc and transferred to a separatory funnel containing 20 mL of saturated aqueous $NH_4Cl$. The organic layer was collected and washed successively with 1 x 20 mL of saturated aqueous NaCl and 2 x 20 mL of half-saturated aqueous NaCl. The organic fraction was dried over $Na_2SO_4$, filtered, and concentrated under reduced pressure. Purification of this material by chromatography on silica gel (gradient elution: 10:0→9:1 $CH_2Cl_2$/acetone) afforded **33** (30 mg, 44%) as a yellow foam.

TLC (4:1 $CH_2Cl_2$/acetone): $R_f = 0.53$

$^1H$ NMR (500 MHz, $CD_3CN$) δ 7.58 (s, 1H), 7.55 (s, 1H), 7.46 (d, $J = 8.2$ Hz, 2H), 7.41 (d, $J = 8.3$ Hz, 2H), 7.19 (s, 1H), 6.18 (d, $J = 6.7$ Hz, 2H), 3.75 (br s, 4H), 3.62 (br s, 2H), 3.42 (br s, 2H), 2.81 (s, 4H) ppm

$^{13}C$ NMR (125 MHz, $CDCl_3$) δ 169.7, 168.4, 152.9, 150.8, 148.5, 142.1, 138.0, 136.3, 129.9, 128.1, 127.7, 107.2, 106.0, 103.7, 79.1, 67.0 (2), 48.2, 42.8, 25.6 ppm

IR (thin film) ν 2921, 1789, 1743, 1631, 1525, 1486, 1428, 1337, 1263, 1225, 1114, 1034 $cm^{-1}$

HRMS ($ESI^+$): $[MH]^+$ calcd for $C_{24}H_{21}N_3O_{11}$, 528.1249; found, 528.1248

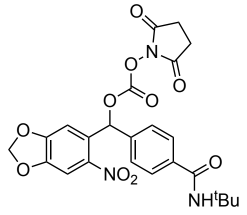


**(4-(*tert*-Butylcarbamoyl)phenyl)(6-nitrobenzo[*d*][1,3]dioxol-5-yl)methyl succinimid-N-yl carbonate (34).** This compound was prepared in an analogous manner to **33** starting from **24** (75 mg, 0.2 mmol) using $CH_3CN$ in place of *N,N*-dimethylformamide. Purification by chromatography on silica gel (gradient elution: 100:0→93:7 $CH_2Cl_2$/acetone) afforded **34** (58 mg, 56%) as a neon yellow film.

TLC (1:1 $CH_2Cl_2$/acetone): $R_f = 0.38$

$^1H$ NMR (500 MHz, $CD_3CN$) δ 7.71 (d, $J = 8.5$ Hz, 2H), 7.58 (s, 1H), 7.54 (s, 1H), 7.46 (d, $J = 8.2$ Hz, 2H), 7.18 (s, 1H), 6.18 (dd, $J = 8.1$, 1.2 Hz, 2H), 5.90 (br s, 1H), 2.81 (s, 4H), 1.46 (s, 9H) ppm

$^{13}C$ NMR (125 MHz, $CDCl_3$) δ 168.5, 166.4, 152.9, 150.8, 148.4, 142.1, 139.0, 136.8, 129.8, 127.8, 127.3, 107.2, 106.0, 103.7, 79.1, 51.9, 28.9, 25.5 ppm

IR (thin film) ν 3397, 2971, 1790, 1742, 1655, 1526, 1506, 1337, 1266, 1222, 1035 $cm^{-1}$

HRMS (ESI$^+$): $[MH]^+$ calcd for $C_{24}H_{23}N_3O_{10}$, 514.1456; found, 514.1419

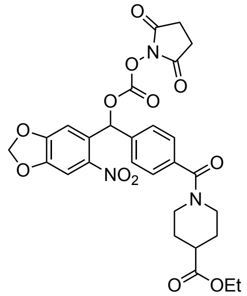

**(4-(4-(Ethylcarbonoyl)piperidine-N-carbonyl)phenyl)(6-nitrobenzo[*d*][1,3]dioxol-5-yl)methyl succinimid-N-yl carbonate (35).** This compound was prepared in an analogous manner to **34** starting from **25** (50 mg, 0.11 mmol). Purification by chromatography on silica gel (gradient elution: 10:0→9:1 $CH_2Cl_2$/acetone) afforded **35** (28 mg, 43%) as a yellow film.

TLC (9:1 $CH_2Cl_2$/acetone): $R_f = 0.36$

$^1$H NMR (500 MHz, $CD_3CN$) δ 7.59 (s, 1H), 7.55 (s, 1H), 7.45 (d, *J* = 8.1 Hz, 2H), 7.40 (d, *J* = 7.9 Hz, 2H), 7.18 (s, 1H), 6.18 (d, *J* = 8.8, 2H), 4.50 (br s, 1H), 4.15 (q, *J* = 7.1 Hz, 2H), 3.70 (br s, 1H), 3.05 (br s, 2H), 2.81 (s, 4H), 2.57 (tt, *J* = 10.7, 4.0 Hz, 1H), 2.03 (br s, 1H), 1.86 (br s, 1H), 1.76 (br s, 1H), 1.67 (br s, 1H), 1.26 (t, *J* = 7.1 Hz, 3H) ppm

$^{13}$C NMR (125 MHz, $CDCl_3$) δ 174.2, 169.7, 168.4, 152.9, 150.8, 148.4, 137.0, 130.1, 130.0, 128.0, 127.4, 127.1, 107.3, 106.0, 103.7, 79.2, 60.8, 47.0, 41.7, 41.1, 28.6, 28.0, 25.6, 20.3, 14.3 ppm

IR (thin film) ν 2927, 1789, 1742, 1629, 1507, 1486, 1428, 1337, 1267, 1224, 1037, 917 cm$^{-1}$

HRMS (ESI$^+$): $[MH]^+$ calcd for $C_{28}H_{27}N_3O_{12}$, 598.1667; found, 598.1667

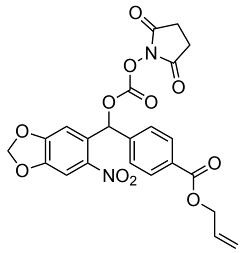

**(4-(Allylcarbonoyl)phenyl)(6-nitrobenzo[*d*][1,3]dioxol-5-yl)methyl succinimid-N-yl carbonate (36).** This compound was prepared in an analogous manner to **34** starting from **28** (50 mg, 0.14 mmol), quenching the reaction after 5 h. Purification by chromatography on silica gel (gradient elution: 100:0→99:1 $CH_2Cl_2$/acetone) afforded **36** (58 mg, 83%) as a yellow film.

TLC (1:1 hexanes/EtOAc): $R_f = 0.27$

$^1$H NMR (400 MHz, $CD_3CN$) δ 8.05 (d, *J* = 8.4 Hz, 2H), 7.59 (d, *J* = 3.3 Hz, 2H), 7.57 (s, 1H), 7.44 (s, 1H), 7.16 (s, 1H), 6.19 (dd, *J* = 9.4, 1.0 Hz, 2H), 6.06 (dddd, *J* = 17.2, 10.8, 5.5, 1.2 Hz, 1H), 5.40 (dd, *J* = 17.3, 1.6 Hz, 1H), 5.27 (dd, *J* = 10.5, 1.4 Hz, 1H), 4.79 (d, *J* = 1.5 Hz, 2H), 2.75 (s, 4H) ppm

$^{13}$C NMR (125 MHz, $CD_3CN$) δ 170.5, 166.2, 154.0, 151.8, 149.7, 143.6, 142.4, 133.6, 131.9, 130.7, 129.9, 128.7, 108.0, 106.4, 105.3, 79.5, 66.4, 26.3 ppm

IR (thin film) ν 2921, 1790, 1743, 1507, 1487, 1337, 1269, 1226, 1102, 1035, 949 cm$^{-1}$

HRMS (ESI$^+$): $[MNa]^+$ calcd for $C_{23}H_{18}N_2O_{11}$, 499.0983; found, 521.0782

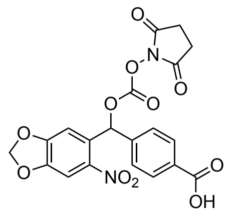

**(4-(Carbonoyl)phenyl)(6-nitrobenzo[*d*][1,3]dioxol-5-yl)methyl succinimid-N-yl carbonate (37).** To a solution of **36** (30 mg, 0.06 mmol) in 7.6 mL of a 1:1 THF/$CH_2Cl_2$ mixture was added dimedone (8.5 mg, 0.06 mmol) and tetrakis(triphenylphosphine)palladium (10.5 mg, 0.01 mmol, 0.15 equiv). The reaction mixture was stirred for 30 min; subsequent purification of this material by chromatography on silica gel (gradient elution: 1:0→0:1 $CH_2Cl_2$/acetone with 0.01% $CF_3CO_2H$) afforded **37** (13 mg, 48%) as a yellow film.

TLC (1:1 $CH_2Cl_2$/acetone): $R_f = 0.05$

$^1H$ NMR (500 MHz, $d_6$-acetone) δ 8.09 (d, $J$ = 8.1 Hz, 2H), 7.66 (d, $J$ = 8.2 Hz, 2H), 7.64 (s, 1H), 7.56 (s, 1H), 7.24 (s, 1H), 6.33 (dd, $J$ = 11.6, 1.0 Hz, 2H), 2.88 (s, 4H) ppm

$^{13}C$ NMR (125 MHz, $d_6$-acetone) δ 170.0, 153.9, 151.8, 149.6, 143.5, 142.2, 130.9, 130.2, 128.5, 107.9, 106.2, 105.2, 79.4, 55.5, 30.7, 26.2 ppm

IR (thin film) ν 2927, 1790, 1713, 1508, 1488, 1426, 1338, 1206, 1141, 1035 $cm^{-1}$

HRMS ($ESI^-$): $[M–H]^-$ calcd for $C_{20}H_{14}N_2O_{11}$, 457.0525; found, 457.0528

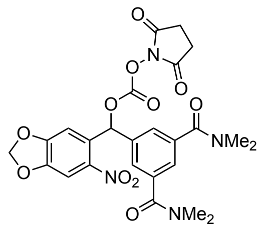

**(3,5-Bis(dimethylcarbamoyl)phenyl)(6-nitrobenzo[*d*][1,3]dioxol-5-yl)methyl succinimid-N-yl carbonate (38).** This compound was prepared in an analogous manner to **34** starting from **26** (42 mg, 0.1 mmol), quenching the reaction after 4 h. Purification by chromatography on silica gel (gradient elution: 9:1→5:5 $CH_2Cl_2$/acetone) afforded **38** (29 mg, 52%) as a neon yellow film.

TLC (3:2 $CH_2Cl_2$/acetone): $R_f = 0.28$

$^1H$ NMR (500 MHz, $CD_3CN$) δ 7.59 (s, 1H), 7.50 (d, $J$ = 1.4 Hz, 2H), 7.41 (t, $J$ = 1.5 Hz, 1H), 7.40 (s, 1H), 7.25 (s, 1H), 6.20 (dd, $J$ = 10.8, 1.0 Hz, 2H), 3.00 (s, 6H), 2.87 (s, 6H), 2.74 (s, 4H) ppm

$^{13}C$ NMR (125 MHz, $CD_3CN$) δ 170.5, 170.2, 154.1, 151.7, 149.7, 143.3, 138.6, 137.9, 130.0, 127.9, 127.1, 107.8, 106.4, 105.3, 79.7, 39.7, 35.3, 26.2 ppm

IR (thin film) ν 2928, 1789, 1743, 1634, 1507, 1395, 1339, 1265, 1225, 1093, 1035 $cm^{-1}$

HRMS ($ESI^+$): $[MH]^+$ calcd for $C_{25}H_{24}N_4O_{11}$, 557.1514; found, 557.1508

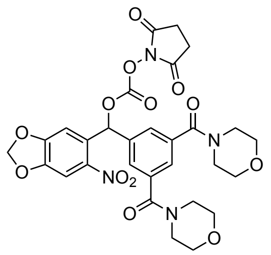

**(3,5-Di(morpholine-4-carbonyl)phenyl)(6-nitrobenzo[*d*][1,3]dioxol-5-yl)methyl succinimid-N-yl carbonate (39).** This compound was prepared in an analogous manner to **34** starting from **27** (50 mg, 0.1 mmol), quenching the reaction after 4 h. Purification by chromatography on silica gel (gradient elution: 9:1→5:5 $CH_2Cl_2$/acetone) afforded **39** (33 mg, 51%) as a neon yellow film.

TLC (3:2 $CH_2Cl_2$/acetone): $R_f = 0.34$

$^1$H NMR (500 MHz, $CD_3CN$) δ 7.60 (s, 1H), 7.52 (d, $J = 1.3$ Hz, 2H), 7.43 (t, $J = 1.5$ Hz, 1H), 7.40 (s, 1H), 7.26 (s, 1H), 6.21 (dd, $J = 7.7$, 1.0 Hz, 2H), 3.65 (br s, 8H), 3.53 (br s, 4H), 3.29 (br s, 4H), 2.74 (s, 4H) ppm

$^{13}$C NMR (125 MHz, $CD_3CN$) δ 170.4, 169.0, 154.0, 151.6, 149.6, 143.2, 138.1, 137.8, 129.9, 128.3, 127.4, 107.7, 106.4, 105.3, 79.5, 67.1(2), 48.8, 43.2, 26.2 ppm

IR (thin film) ν 2922, 2857, 1789, 1743, 1633, 1526, 1487, 1427, 1338, 1269, 1226, 1114 $cm^{-1}$

HRMS (ESI$^+$): $[MH]^+$ calcd for $C_{29}H_{28}N_4O_{13}$, 641.1726; found, 641.1726

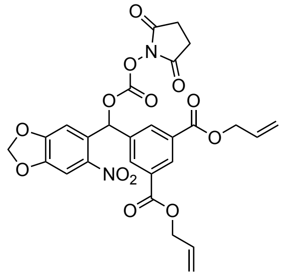

**(3,5-Di(allylcarbonoyl)phenyl)(6-nitrobenzo[d][1,3]dioxol-5-yl)methyl succinimid-N-yl carbonate (40).** This compound was prepared in an analogous manner to **34** starting from **29** (47 mg, 0.1 mmol). Purification by chromatography on silica gel (gradient elution: 50:0→49:1 $CH_2Cl_2$/acetone) afforded **40** (24 mg, 38%) as a yellow film.

TLC (1:1 hexanes/EtOAc): $R_f = 0.36$

$^1$H NMR (500 MHz, $CD_3CN$) δ 8.58 (t, $J = 1.6$ Hz, 1H), 8.29 (d, $J = 1.6$ Hz, 2H), 7.60 (s, 1H), 7.47 (s, 1H), 7.21 (s, 1H), 6.21 (d, $J = 10.0$ Hz, 2H), 6.12–6.02 (m, 2H), 5.41 (dd, $J = 17.3$, 1.5 Hz, 2H), 5.30 (dd, $J = 10.6$, 1.4 Hz, 2H), 4.83 (dd, $J = 5.6$, 1.5 Hz, 4H), 2.75 (s, 4H) ppm

$^{13}$C NMR (125 MHz, $CD_3CN$) δ 170.4, 165.5, 154.1, 151.7, 149.9, 143.6, 139.1, 133.4, 133.3, 132.5, 131.5, 129.7, 118.8, 107.9, 106.5, 105.4, 79.2, 66.9, 26.3 ppm

IR (thin film) ν 3084, 2947, 1791, 1744, 1507, 1488, 1426, 1370, 1337, 1266, 1232, 1091, 1036, 987, 914 $cm^{-1}$

HRMS (ESI$^+$): $[MNa]^+$ calcd for $C_{27}H_{22}N_2O_{13}$, 605.1014; found, 605.1006

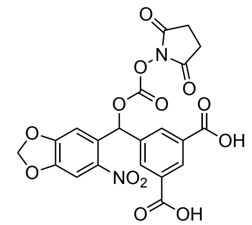


**(3,5-Dicarbonoylphenyl)(6-nitrobenzo[d][1,3]dioxol-5-yl)methyl succinimid-N-yl carbonate (41).** To a solution of **40** (50 mg, 0.06 mmol) in 7.6 mL of a 1:1 THF/$CH_2Cl_2$ mixture was added dimedone (17 mg, 0.12 mmol) and tetrakis(triphenylphosphine)palladium (21 mg, 0.02 mmol, 0.3 equiv). The reaction mixture was stirred for 30 minutes then purified by chromatography on silica gel (gradient elution: 1:0→0:1 $CH_2Cl_2$/acetone with 0.01% $CF_3CO_2H$). The product was concentrated under reduced pressure and washed with 5 x 2 mL $CHCl_3$ to afford **41** (22 mg, 73%) as a yellow solid.

TLC (1:1 $CH_2Cl_2$/acetone): $R_f$ = 0.00

$^1$H NMR (500 MHz, $d_6$-acetone) δ 8.69 (s, 1H), 8.39 (s, 2H), 7.67 (s, 1H), 7.62 (s, 1H), 7.33 (s, 1H), 6.34 (d, $J$ = 7.2 Hz, 2H), 2.88 (s, 4H) ppm

$^{13}$C NMR (125 MHz, $d_6$-acetone) δ 170.0, 166.8, 154.1, 151.7, 149.8, 143.7, 139.0, 133.6, 132.9, 132.1, 130.1, 107.8, 106.4, 105.3, 30.7, 26.3 ppm

IR (thin film) ν 2921, 1791, 1739, 1507, 1337, 1266, 1202, 1035, 909 cm$^{-1}$

HRMS (ESI$^-$): [M–H]$^-$ calcd for $C_{21}H_{14}N_2O_{13}$, 501.0423; found, 501.0423

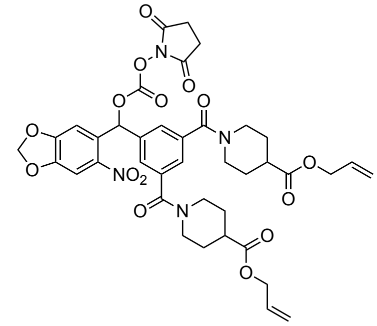


**(3,5-Di(4-(allylcarbonoyl)piperidine-N-carbonyl)phenyl)(6-nitrobenzo[*d*][1,3]dioxol-5-yl)methyl succinimid-N-yl carbonate (42).** This compound was prepared in an analogous manner to **34** starting from **31** (50 mg, 0.075 mmol), quenching the reaction after 5 h. Purification by chromatography on silica gel (gradient elution: 5:0→4:1 $CH_2Cl_2$/acetone) afforded **42** (38 mg, 63%) as a yellow film.

TLC (4:1 $CH_2Cl_2$/acetone): $R_f$ = 0.58

$^1$H NMR (400 MHz, $CDCl_3$) δ 7.57 (s, 1H), 7.48 (s, 1H), 7.47 (s, 2H), 7.42 (s, 1H), 7.23 (s, 1H), 6.19 (d, $J$ = 2.1 Hz, 2H), 5.90 (ddt, $J$ = 17.0, 10.2, 5.7 Hz, 2H), 5.30 (dd, $J$ = 17.3, 1.5 Hz, 2H), 5.23 (dd, $J$ = 10.4, 1.3 Hz, 2H), 4.48 (dd, $J$ = 5.7, 1.4 Hz, 4H), 4.47 (br s, 2H), 3.62 (br s, 2H), 3.05 (br s, 4H), 2.80 (s, 4H), 2.60 (tt, $J$ = 10.2, 4.0 Hz, 2H), 2.03 (br s, 2H), 1.87 (br s, 2H), 1.77 (br s, 2H), 1.61 (br s, 2H) ppm

$^{13}$C NMR (125 MHz, $CDCl_3$) δ 173.7, 168.7, 168.3, 153.1, 150.7, 148.6, 142.0, 137.2, 137.1, 132.1, 129.4, 127.7, 126.5, 118.5, 107.0, 106.1, 103.8, 78.9, 65.4, 47.2, 41.7, 41.0, 28.6, 27.8, 25.5 ppm

IR (thin film) ν 2930, 2863, 1789, 1742, 1633, 1526, 1487, 1428, 1373, 1316, 1268, 1226, 1036, 927 cm$^{-1}$

HRMS (ESI$^+$): [MH]$^+$ calcd for $C_{39}H_{40}N_4O_{15}$, 805.2563; found, 805.2550

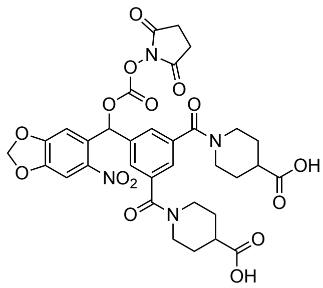

**(3,5-Di(4-carbonoylpiperidine-N-carbonyl)phenyl)(6-nitrobenzo[*d*][1,3]dioxol-5-yl)methyl succinimid-N-yl carbonate (43).** This compound was prepared in an analogous manner to **41** starting from **42** (30 mg, 0.04 mmol). Purification by chromatography on silica gel (gradient elution: 1:0→0:1 $CH_2Cl_2$/acetone with 0.01% $CF_3CO_2H$) afforded **43** (12 mg, 44%) as a yellow foam.

TLC (1:1 $CH_2Cl_2$/acetone): $R_f = 0.00$

$^1$H NMR (300 MHz, $d_6$-acetone) δ 7.99 (s, 1H), 7.59 (s, 2H), 7.50 (s, 1H), 7.47 (s, 1H), 7.32 (s, 1H), 6.30 (d, *J* = 5.9 Hz, 2H), 4.38 (br s, 2H), 3.61 (br s, 2H), 3.11 (br s, 2H), 2.98 (br s, 2H), 2.84 (s, 4H), 2.67–2.54 (m, 2H), 1.91 (br s, 4H), 1.61 (d, *J* = 13.4 Hz, 4H) ppm

$^{13}$C NMR (125 MHz, $d_6$-acetone) δ 175.7, 170.0, 168.9, 154.0, 151.8, 149.6, 138.4, 130.2, 128.1, 127.3, 107.8, 106.3, 105.2, 79.5, 55.5, 47.6, 42.1, 41.3, 30.7, 28.8, 28.7, 26.3 ppm

IR (thin film) ν 2931, 1789, 1742, 1615, 1507, 1487, 1431, 1338, 1267, 1224, 1092, 1033 $cm^{-1}$

HRMS ($ESI^+$): $[MH]^+$ calcd for $C_{33}H_{32}N_4O_{15}$, 725.1937; found, 725.1927

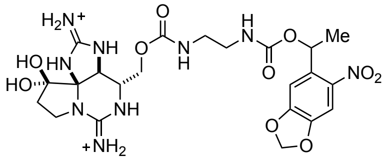

**(1-(6-Nitrobenzo[*d*][1,3]dioxol-5-yl)ethyl) saxitoxin N21-ethylcarbamate (2).** To an ice-cold solution of **1** (1.15 μmol) in 115 μL of pH 8.5 aqueous phosphate buffer (0.1 M $Na_2HPO_4$/$Na_3PO_4$) was added a solution of **32** (0.6 mg, 1.72 μmol, 1.5 equiv) in 115 μL of $CH_3CN$. The reaction flask was stoppered and placed in a sonication bath for 30 seconds. The flask was then wrapped in foil and the contents stirred for 6 h. Following this time, the reaction was quenched by the addition of 11.5 μL of 1.0 M aqueous $CF_3CO_2H$. The reaction mixture was diluted with 1.75 mL of a 9:1 10 mM aqueous $CF_3CO_2H$/$CH_3CN$ solution and filtered through a Fisher 0.22 μm PTFE filter. The product was purified by reversed-phase HPLC (Silicycle SiliaChrom AQ C18, 5 μm, 10 x 250 mm column, eluting with a gradient flow of 10→50% $CH_3CN$ in 10 mM aqueous $CF_3CO_2H$ over 80 min, 214 nm UV detection). At a flow rate of 4 mL/min, **2** had a retention time of 30–32 min and was isolated as a white powder following lyophilization (0.52 μmol, 45%, $^1$H NMR quantitation).

$^1$H NMR (600 MHz, $D_2O$, single diastereomer) δ 7.94 (s, 1H), 7.57 (s, 1H), 7.15 (s, 1H), 6.17 (s, 2H), 6.11 (q, *J* = 6.4 Hz, 1H), 4.67 (s, 1H), 4.28 (t, *J* = 10.8 Hz, 1H), 4.03–3.93 (m, 1H), 3.86–3.69 (m, 2H), 3.57 (t, *J* = 8.7 Hz, 1H), 3.37–3.10 (m, 4H), 2.45–2.26 (m, 1H), 1.60 (d, *J* = 6.5 Hz, 3H) ppm

HRMS ($ESI^+$): $[MH]^+$ calcd for $C_{22}H_{29}N_9O_{10}$, 580.2110; found, 580.2110

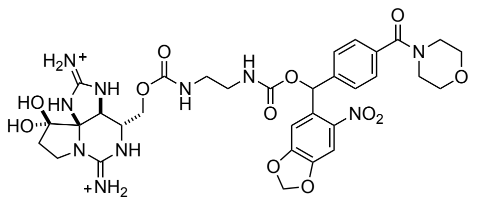

**((4-(Morpholine-4-carbonyl)phenyl)(6-nitrobenzo[*d*][1,3]dioxol-5-yl)methyl) saxitoxin N21-ethylcarbamate (3).** This compound was prepared in an analogous manner to **2** starting from **1** (1.04 µmol) and **33** (1.6 mg, 3.12 µmol, 3.0 equiv), quenching the reaction after 5.5 h. At a flow rate of 4 mL/min (gradient flow of 10→50% $CH_3CN$ in 10 mM aqueous $CF_3CO_2H$ over 80 min, 214 nm UV detection), **3** had a retention time of 28–31 min and was isolated as a white powder following lyophilization (0.46 µmol, 45%, $^1H$ NMR quantitation).

$^1H$ NMR (600 MHz, $D_2O$, single diastereomer) δ 7.67 (s, 1H), 7.56 (d, $J$ = 8.1 Hz, 2H), 7.50 (d, $J$ = 8.0 Hz, 2H), 7.31 (s, 1H), 7.12 (s, 1H), 6.20 (s, 2H), 4.70 (s, 1H), 4.29 (t, $J$ = 10.9 Hz, 1H), 3.95 (d, $J$ = 6.8 Hz, 1H), 3.86 (s, 2H), 3.84–3.82 (m, 1H), 3.80 (s, 2H), 3.70 (s, 2H), 3.69–3.63 (m, 1H), 3.59–3.54 (m, 1H), 3.50 (s, 2H), 3.38–3.29 (m, 1H), 3.27–3.12 (m, 3H), 2.47–2.32 (m, 1H) ppm

HRMS (ESI$^+$): [MH]$^+$ calcd for $C_{32}H_{38}N_{10}O_{12}$, 755.2743; found, 755.2727

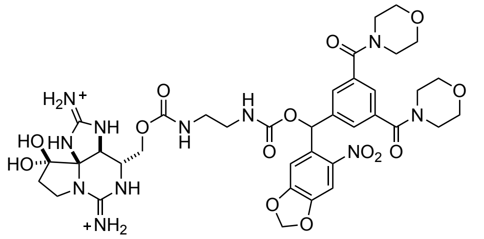

**(3,5-Di(morpholine-4-carbonyl)phenyl)(6-nitrobenzo[*d*][1,3]dioxol-5-yl)methyl axitoxin N21-ethylcarbamate (4).** This compound was prepared in an analogous manner to **8** starting from **1** (1.41 µmol) and **39** (2.8 mg, 3.10 µmol, 2.2 equiv), quenching the reaction after 4.5 h. At a flow rate of 4 mL/min (gradient flow of 0→40% $CH_3CN$ in 10 mM aqueous $CF_3CO_2H$ over 80 min, 214 nm UV detection), **4** had a retention time of 48–51 min and was isolated as a white powder following lyophilization (0.88 µmol, 62%, $^1H$ NMR quantitation).

$^1H$ NMR (600 MHz, $D_2O$, mixture of diastereomers) δ 7.70–7.64 (m, 1H), 7.61–7.55 (m, 2H), 7.54–7.49 (m, 1H), 7.35–7.27 (m, 1H), 7.26–7.17 (m, 1H), 6.26–6.19 (m, 2H), 4.74–4.67 (m, 1H), 4.35–4.18 (m, 1H), 4.04–3.89 (m, 1H), 3.84 (d, $J$ = 5.0 Hz, 4H), 3.83–3.80 (m, 2H), 3.79 (d, $J$ = 4.9 Hz, 4H), 3.68 (d, $J$ = 5.5 Hz, 4H), 3.64–3.51 (m, 1H), 3.44 (d, $J$ = 4.7 Hz, 4H), 3.39–3.33 (m, 1H), 3.30–3.11 (m, 3H), 2.48–2.31 (m, 2H) ppm

HRMS (ESI$^+$): [MH]$^+$ calcd for $C_{37}H_{45}N_{11}O_{14}$, 868.3220; found, 868.3217

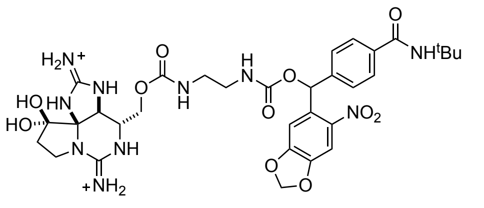

**(4-(*tert*-Butylcarbamoyl)phenyl)(6-nitrobenzo[*d*][1,3]dioxol-5-yl)methyl saxitoxin N21-ethylcarbamate (5).** This compound was prepared in an analogous manner to **2** starting from **1** (1.2 µmol) and **34** (2.5 mg, 3.6 µmol, 3.0 equiv), replacing $CH_3CN$ with DMSO and quenching the reaction after 4 h. At a flow rate of 4 mL/min (gradient flow of 10→50% $CH_3CN$ in 10 mM aqueous $CF_3CO_2H$ over 80 min, 214 nm

UV detection), **5** had a retention time of 48 min and was isolated as a white powder following lyophilization (0.31 µmol, 26%, $^1$H NMR quantitation).

$^1$H NMR (600 MHz, $D_2O$, mixture of diastereomers) δ 7.75–7.70 (m, 2H), 7.70–7.65 (m, 1H), 7.55–7.48 (m, 2H), 7.32–7.27 (m, 1H), 7.22–7.12 (m, 1H), 6.25–6.17 (m, 2H), 4.71–4.65 (m, 1H), 4.28 (t, *J* = 10.6 Hz, 1H), 3.98–3.89 (m, 1H), 3.84–3.65 (m, 2H), 3.60–3.46 (m, 1H), 3.40–3.28 (m, 1H), 3.28–3.17 (m, 3H), 2.48–2.30 (m, 2H), 1.45 (s, 9H) ppm

HRMS (ESI$^+$): $[MH]^+$ calcd for $C_{32}H_{40}N_{10}O_{11}$, 741.2951; found, 741.2946

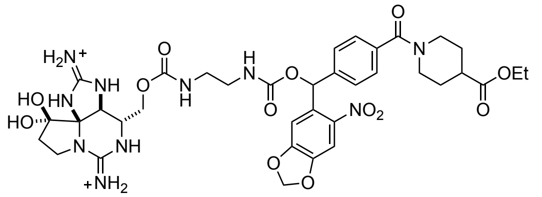


**(4-(4-(Ethylcarbonoyl)piperidine-N-carbonyl)phenyl)(6-nitrobenzo[*d*][1,3]dioxol-5-yl)methyl saxitoxin N21-ethylcarbamate (6).** This compound was prepared in an analogous manner to **2** starting from **1** (1.37 µmol) and **3.37** (2.5 mg, 4.11 µmol, 3.0 equiv), quenching the reaction after 4.5 h. At a flow rate of 4 mL/min (gradient flow of 10→50% $CH_3CN$ in 10 mM aqueous $CF_3CO_2H$ over 80 min, 214 nm UV detection), **6** had a retention time of 42–44 min and was isolated as a white powder following lyophilization (0.24 µmol, 18%, $^1$H NMR quantitation).

$^1$H NMR (600 MHz, $D_2O$, mixture of diastereomers) δ 7.57–7.49 (m, 1H), 7.43–7.36 (m, 2H), 7.34–7.27 (m, 2H), 7.17–7.10 (m, 1H), 7.09–6.92 (m, 1H), 6.10–6.02 (m, 2H), 4.51 (s, 1H), 4.31–4.24 (m, 1H), 4.16 (t, *J* = 11.0 Hz, 1H), 4.03 (q, *J* = 6.9 Hz, 2H), 3.73 (d, *J* = 11.8 Hz, 1H), 3.64 (t, *J* = 10.2 Hz, 1H), 3.58–3.52 (m, 2H), 3.36 (d, *J* = 9.3 Hz, 1H), 3.28–3.14 (m, 1H), 3.12–3.02 (m, 3H), 3.00–2.91 (m, 2H), 2.66–2.58 (m, 1H), 2.31–2.15 (m, 2H), 1.99–1.91 (m, 1H), 1.74 (d, *J* = 13.4 Hz, 1H), 1.65–1.54 (m, 1H), 1.53–1.41 (m, 1H), 1.11 (t, *J* = 7.3 Hz, 3H) ppm

HRMS (ESI$^+$): $[MH]^+$ calcd for $C_{36}H_{44}N_{10}O_{13}$, 825.3162; found, 825.3145

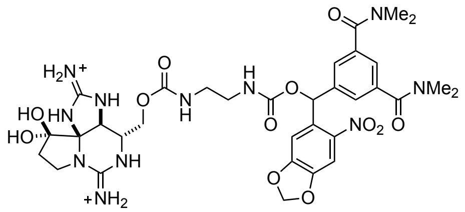


**(3,5-Bis(dimethylcarbamoyl)phenyl)(6-nitrobenzo[*d*][1,3]dioxol-5-yl)methyl saxitoxin N21-ethylcarbamate (7).** This compound was prepared in an analogous manner to **8** starting from **1** (1.5 µmol) and **38** (1.7 mg, 3.0 µmol, 2.0 equiv), quenching the reaction after 4.5 h. At a flow rate of 4 mL/min (gradient flow of 0→40% $CH_3CN$ in 10 mM aqueous $CF_3CO_2H$ over 80 min, 214 nm UV detection), **7** had a retention time of 49–51 min and was isolated as a white powder following lyophilization (0.43 µmol, 29%, $^1$H NMR quantitation).

$^1$H NMR (600 MHz, $D_2O$, single diastereomer) δ 7.67 (s, 1H), 7.58 (s, 2H), 7.51 (s, 1H), 7.31 (s, 1H), 7.18 (s, 1H), 6.21 (s, 2H), 4.71 (s, 1H), 4.30–4.19 (m, 1H), 4.03–3.93 (m, 1H), 3.89–3.72 (m, 2H), 3.61–3.50 (m, 1H), 3.32–3.26 (m, 1H), 3.26–3.16 (m, 3H), 3.11 (s, 6H), 2.97 (s, 6H), 2.48–2.31 (m, 2H) ppm

HRMS (ESI$^+$): $[MH]^+$ calcd for $C_{33}H_{41}N_{11}O_{12}$, 784.3009; found, 784.2995

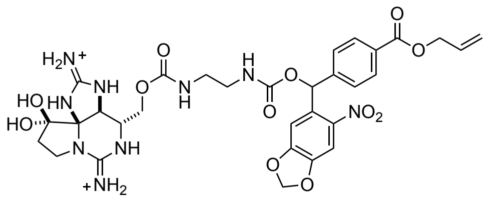

**(4-(Allylcarbonoyl)phenyl)(6-nitrobenzo[*d*][1,3]dioxol-5-yl)methyl saxitoxin N21-ethylcarbamate (8).** This compound was prepared in an analogous manner to **2** starting from **1** (1.23 µmol) in pH 9.5 aqueous phosphate buffer (0.1 M $NaHCO_3$/$Na_2CO_3$) and **3.38** (1.2 mg, 2.47 µmol, 2.0 equiv) in $CH_3CN$, quenching the reaction after 4 h. At a flow rate of 4 mL/min (gradient flow of 10→60% $CH_3CN$ in 10 mM aqueous $CF_3CO_2H$ over 50 min, 214 nm UV detection), **8** had a retention time of 30–31 min and was isolated as a white powder following lyophilization (0.11 µmol, 9%, $^1$H NMR quantitation).

$^1$H NMR (600 MHz, $D_2O$, mixture of diastereomers) δ 8.13–8.06 (m, 2H), 7.71–7.65 (m, 1H), 7.63–7.55 (m, 2H), 7.37–7.30 (m, 1H), 7.20–7.13 (m, 1H), 6.25–6.17 (m, 2H), 6.15–6.07 (m, 1H), 5.46 (d, *J* = 16.7 Hz, 1H), 5.36 (d, *J* = 11.1 Hz, 1H), 4.89 (d, *J* = 5.7 Hz, 2H), 4.71–4.66 (m, 1H), 4.32–4.22 (m, 1H), 4.01–3.89 (m, 1H), 3.86–3.68 (m, 2H), 3.63–3.48 (m, 1H), 3.44–3.28 (m, 1H), 3.27–3.12 (m, 3H), 2.48–2.32 (m, 2H) ppm

HRMS (ESI$^+$): [MH]$^+$ calcd for $C_{31}H_{35}N_9O_{12}$, 726.2478; found, 726.2477

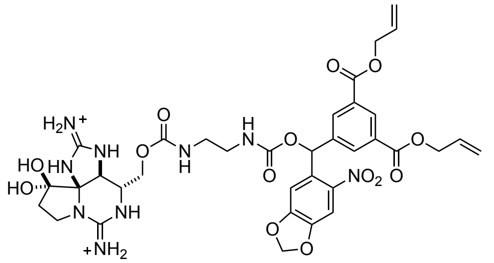

**(3,5-Di(allylcarbonoyl)phenyl)(6-nitrobenzo[d][1,3]dioxol-5-yl)methyl saxitoxin N21-ethylcarbamate (9).** This compound was prepared in an analogous manner to **8** starting from **1** (1.53 µmol) and **40** (1.8 mg, 3.06 µmol, 2.0 equiv), quenching the reaction after 4.5 h. At a flow rate of 4 mL/min (gradient flow of 10→60% $CH_3CN$ in 10 mM aqueous $CF_3CO_2H$ over 50 min, 214 nm UV detection), **9** had a retention time of 34–36 min and was isolated as a white powder following lyophilization (0.15 µmol, 10%, $^1$H NMR quantitation).

$^1$H NMR (600 MHz, $D_2O$, mixture of diastereomers) δ 8.61 (s, 1H), 8.31–8.24 (m, 2H), 7.71–7.63 (m, 1H), 7.38–7.29 (m, 1H), 7.21–7.11 (m, 1H), 6.25–6.18 (m, 2H), 6.11 (ddt, *J* = 16.5, 10.8, 5.6 Hz, 2H), 5.44 (d, *J* = 17.3 Hz, 2H), 5.36 (d, *J* = 10.5 Hz, 2H), 4.90 (d, *J* = 5.8 Hz, 4H), 4.72–4.66 (m, 1H), 4.34–4.18 (m, 1H), 4.01–3.89 (m, 1H), 3.85–3.66 (m, 2H), 3.63–3.50 (m, 1H), 3.42–3.28 (m, 1H), 3.28–3.11 (m, 3H), 2.47–2.32 (m, 2H) ppm

HRMS (ESI$^+$): [MH]$^+$ calcd for $C_{35}H_{39}N_9O_{14}$, 810.2689; found, 810.2682

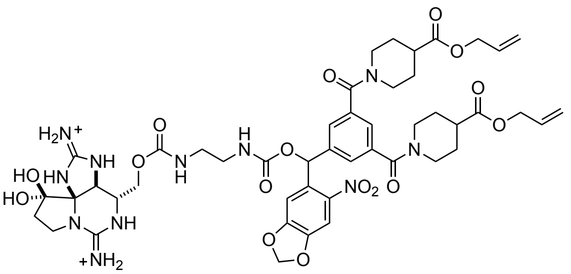

**(3,5-Di(4-(allylcarbonoyl)piperidine-N-carbonyl)phenyl)(6-nitrobenzo[*d*][1,3]dioxol-5-yl)methyl saxitoxin N21-ethylcarbamate (10).** This compound was prepared in an analogous manner to **8** starting from **1** (0.83 µmol) and **42** (1.3 mg, 1.66 µmol, 2.0 equiv), quenching the reaction after 4.5 h. The reaction mixture was diluted with 1.8 mL of a 7:3 10 mM aqueous $CF_3CO_2H/CH_3CN$ solution and filtered through a Fisher 0.22 µm PTFE filter prior to purification by reversed-phase HPLC. At a flow rate of 4 mL/min (gradient flow of 10→60% $CH_3CN$ in 10 mM aqueous $CF_3CO_2H$ over 50 min, 214 nm UV detection), **10** had a retention time of 31–33 min and was isolated as a white powder following lyophilization (0.29 µmol, 35%, $^1$H NMR quantitation).

$^1$H NMR (600 MHz, $D_2O$, mixture of diastereomers) δ 7.71–7.63 (m, 1H), 7.56–7.49 (m, 2H), 7.47 (s, 1H), 7.37–7.28 (m, 1H), 7.27–7.09 (m, 1H), 6.27–6.16 (m, 2H), 6.00 (ddt, *J* = 16.5, 10.9, 5.7 Hz, 2H), 5.36 (d, *J* = 17.4 Hz, 2H), 5.32 (d, *J* = 10.5 Hz, 2H), 4.73–4.70 (m, 1H), 4.68 (d, *J* = 5.6 Hz, 4H), 4.47–4.39 (m, 2H), 4.32–4.20 (m, 1H), 4.04–3.89 (m, 1H), 3.86–3.71 (m, 2H), 3.67–3.48 (m, 3H), 3.33–3.14 (m, 6H), 3.14–3.06 (m, 2H), 2.86–2.79 (m, 2H), 2.48–2.32 (m, 2H), 2.12 (d, *J* = 13.5 Hz, 2H), 1.95–1.85 (m, 2H), 1.81–1.70 (m, 2H), 1.66–1.45 (m, 2H) ppm

HRMS (ESI$^+$): $[MH]^+$ calcd for $C_{47}H_{57}N_{11}O_{16}$, 1032.4058; found, 1032.4020

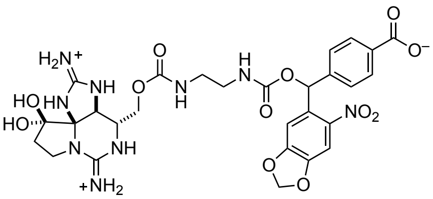

**(4-(Carbonoyl)phenyl)(6-nitrobenzo[*d*][1,3]dioxol-5-yl)methyl saxitoxin N21-ethylcarbamate (11).** This compound was prepared in an analogous manner to **8** starting from **1** (2.28 µmol) and **37** (2.1 mg, 4.55 µmol, 2.0 equiv), quenching the reaction after 4.5 h. At a flow rate of 4 mL/min (gradient flow of 10→40% $CH_3CN$ in 10 mM aqueous $CF_3CO_2H$ over 60 min, 214 nm UV detection), **11** had a retention time of 28–32 min and was isolated as a white powder following lyophilization (0.57 µmol, 25%, $^1$H NMR quantitation).

$^1$H NMR (600 MHz, $D_2O$, mixture of diastereomers) δ 8.00–7.93 (m, 2H), 7.70–7.62 (m, 1H), 7.57–7.48 (m, 2H), 7.34–7.27 (m, 1H), 7.19–7.05 (m, 1H), 6.25–6.16 (m, 2H), 4.48 (s, 1H), 4.32–4.21 (m, 1H), 4.06–3.85 (m, 1H), 3.85–3.71 (m, 1H), 3.68–3.42 (m, 2H), 3.36–3.21 (m, 3H), 3.20–3.11 (m, 1H), 2.48–2.30 (m, 2H) ppm

HRMS (ESI$^+$): $[MH]^+$ calcd for $C_{28}H_{31}N_9O_{12}$, 686.2165; found, 686.2165

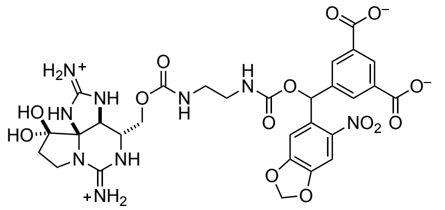

**(3,5-Dicarbonoylphenyl)(6-nitrobenzo[d][1,3]dioxol-5-yl)methyl saxitoxin N21-ethylcarbamate (12).** This compound was prepared in an analogous manner to **8** starting from **1** (2.38 µmol) and **41** (2.4 mg, 4.75 µmol, 2.0 equiv), quenching the reaction after 4 h. At a flow rate of 4 mL/min (gradient flow of 0→40% $CH_3CN$ in 10 mM aqueous $CF_3CO_2H$ over 80 min, 214 nm UV detection), **12** had a retention time of 44–47 min and was isolated as a white powder following lyophilization (1.13 µmol, 47%, $^1$H NMR quantitation).

$^1$H NMR (600 MHz, $D_2O$, single diastereomer) δ 8.19 (s, 2H), 7.95 (s, 1H), 7.66 (s, 1H), 7.36 (s, 1H), 7.07 (s, 1H), 6.19 (s, 2H), 4.64 (s, 1H), 4.24 (t, $J$ = 11.0 Hz, 1H), 3.88–3.85 (m, 1H), 3.81–3.75 (m, 1H), 3.57–3.53 (m, 1H), 3.51–3.41 (m, 2H), 3.32–3.21 (m, 2H), 3.19–3.09 (m, 1H), 2.33 (t, $J$ = 10.5 Hz, 1H) ppm

HRMS (ESI$^+$): [MH]$^+$ calcd for $C_{29}H_{31}N_9O_{14}$, 730.2063; found, 730.2062

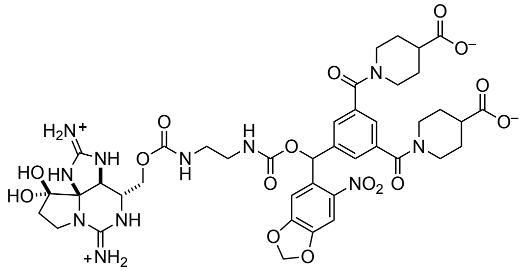

**3,5-Di(4-carbonoylpiperidine-N-carbonyl)phenyl)(6-nitrobenzo[*d*][1,3]dioxol-5-yl)methyl saxitoxin N21-ethylcarbamate (13).** This compound was prepared in an analogous manner to **8** starting from **1** (1.92 µmol) and **43** (2.8 mg, 3.84 µmol, 2.0 equiv), quenching the reaction after 4 h. At a flow rate of 4 mL/min (gradient flow of 0→40% $CH_3CN$ in 10 mM aqueous $CF_3CO_2H$ over 80 min, 214 nm UV detection), **13** had a retention time of 46–48 min and was isolated as a white powder following lyophilization (1.08 µmol, 56%, $^1$H NMR quantitation).

$^1$H NMR (600 MHz, $D_2O$, mixture of diastereomers) δ 7.70–7.63 (m, 1H), 7.58–7.50 (m, 2H), 7.48 (s, 1H), 7.36–7.28 (m, 1H), 7.27–7.12 (m, 1H), 6.26–6.17 (m, 2H), 4.74–4.67 (m, 1H), 4.49–4.39 (m, 2H), 4.31–4.18 (m, 1H), 4.04–3.87 (m, 1H), 3.85–3.71 (m, 2H), 3.66–3.49 (m, 3H), 3.37–3.13 (m, 6H), 3.13–3.05 (m, 2H), 2.80–2.69 (m, 2H), 2.48–2.32 (m, 2H), 2.13–2.05 (m, 2H), 1.94–1.81 (m, 2H), 1.78–1.66 (m, 2H), 1.62–1.42 (m, 2H) ppm

HRMS (ESI$^+$): [MH]$^+$ calcd for $C_{41}H_{49}N_{11}O_{16}$, 952.3432; found, 952.3409

### References


1. Thomas-Tran, R. Systematic Mapping of the Voltage-Gated Sodium Channel Outer Pore Using Modified Saxitoxins. Ph.D. Dissertation, Stanford University, Stanford, CA **2016**.
2. Mulcahy, J. V., Walker, J. R., Merit, J. E., Whitehead, A. & Du Bois, J. Synthesis of the Paralytic Shellfish Poisons (+)-Gonyautoxin 2, (+)-Gonyautoxin 3, and (+)-11,11-Dihydroxysaxitoxin. *J. Am. Chem. Soc.* **138**, 5994–6001 (2016).
3. Parsons, W. H. Saxitoxin-Derived Probes for the Study of Voltage-Gated Sodium Channels in Living Cells and Animals. Ph.D. Dissertation, Stanford University, Stanford, CA **2013**.
4. Parsons, W. H. & Du Bois, J. Maleimide conjugates of saxitoxin as covalent inhibitors of voltage-gated sodium channels. *J. Am. Chem. Soc.* **135**, 10582–10585 (2013).

### 4. [1]H NMR Spectra

#### Triprotected saxitoxin N21-ethylamine (14)

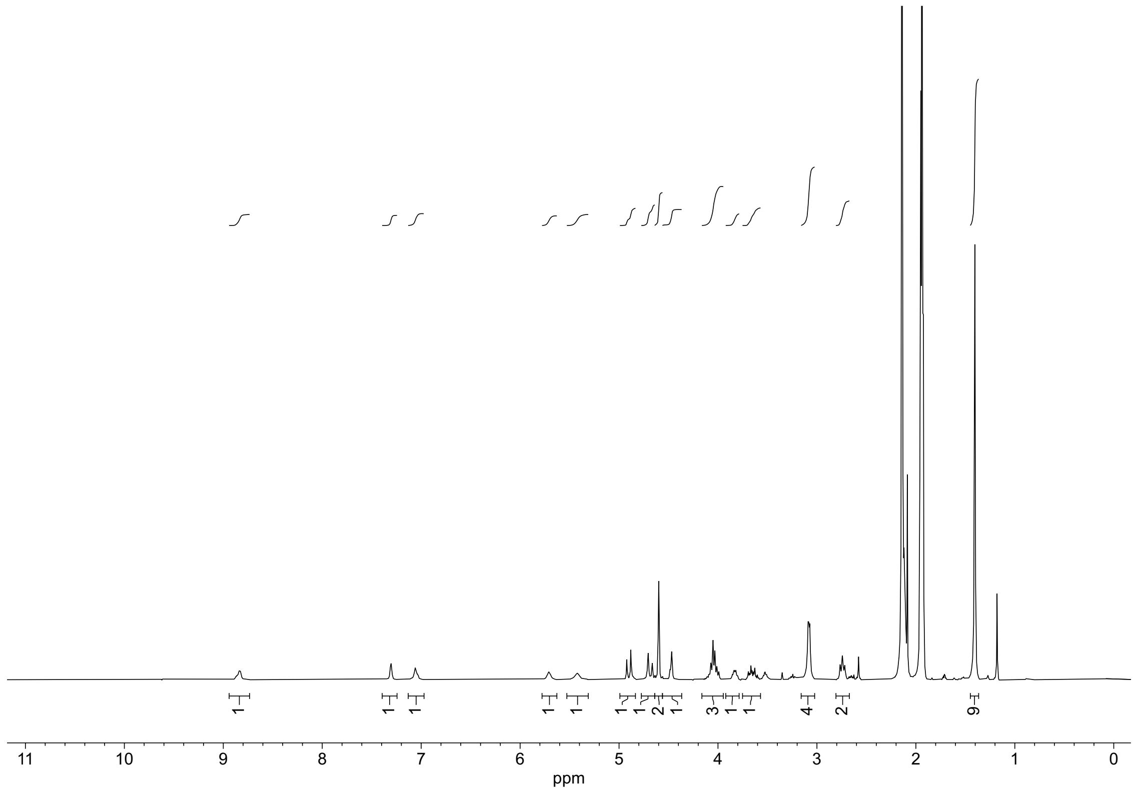


#### (4-Iodophenyl)(morpholino)methanone (15)

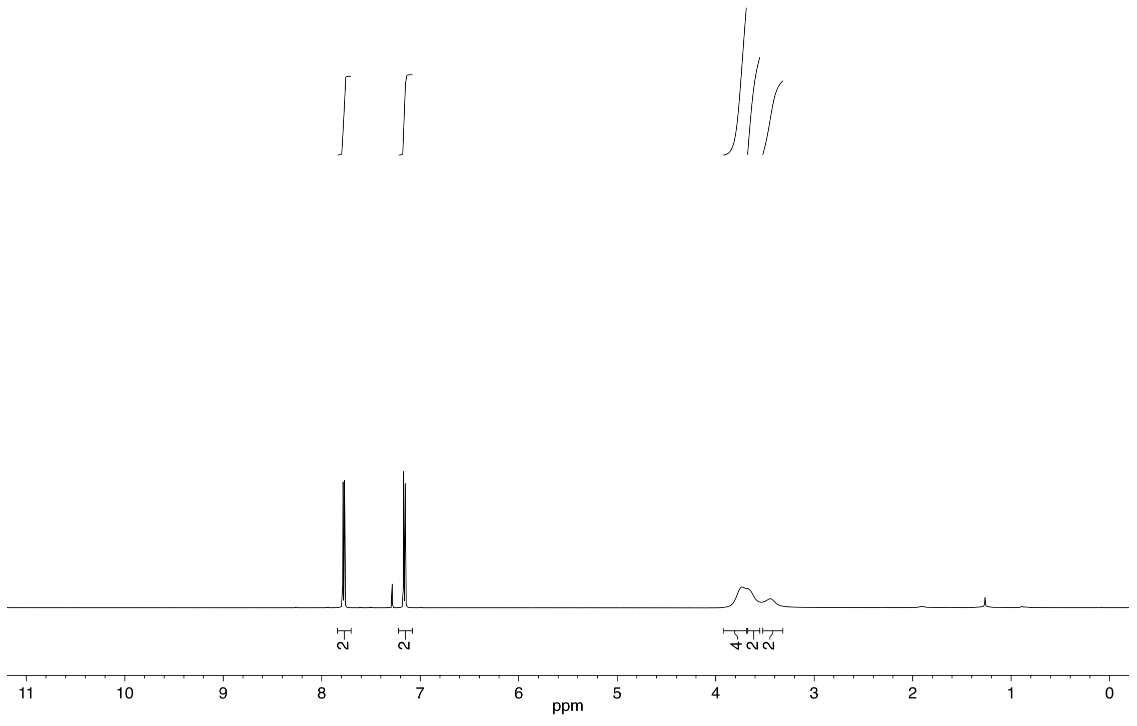

#### *N*-(*tert*-Butyl)-4-iodobenzamide (16)

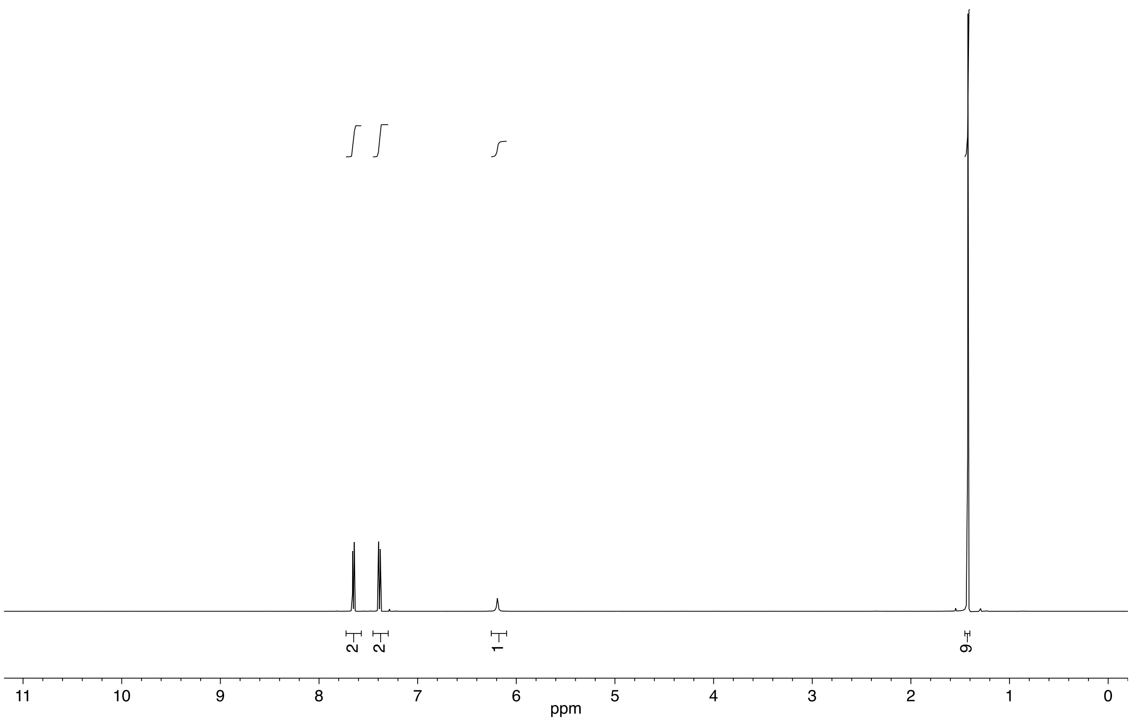


#### Ethyl 1-(4-iodobenzoyl)piperidine-4-carboxylate (17)

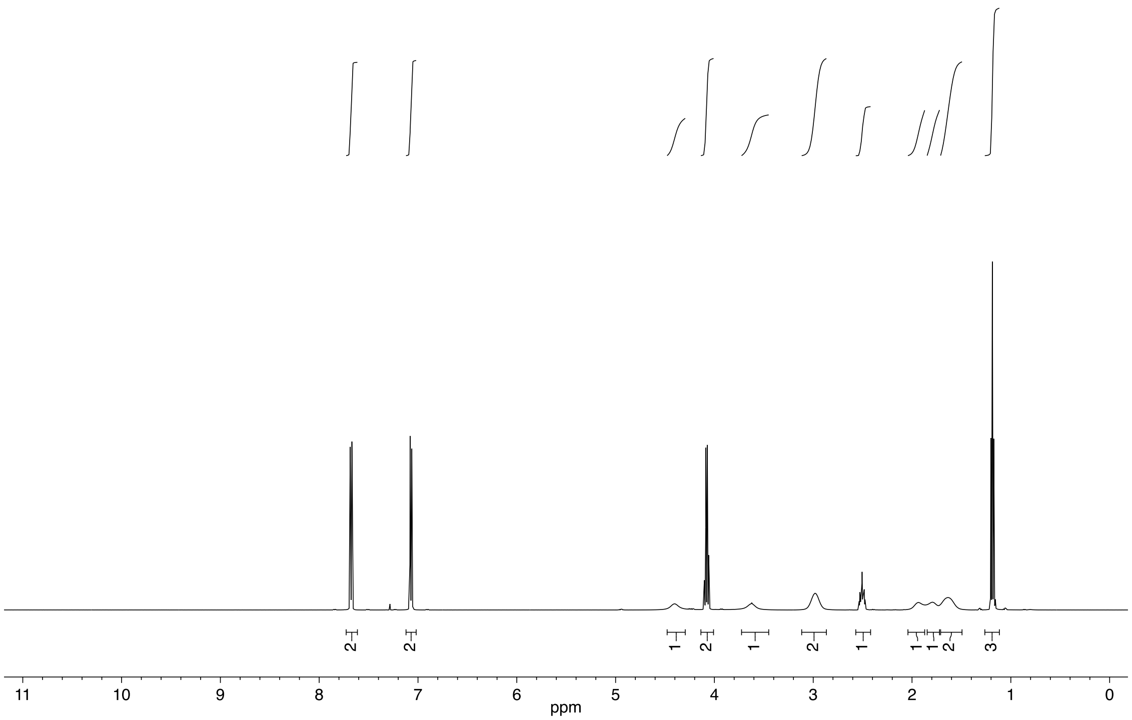

**Allyl 4-iodobenzoate (18)**

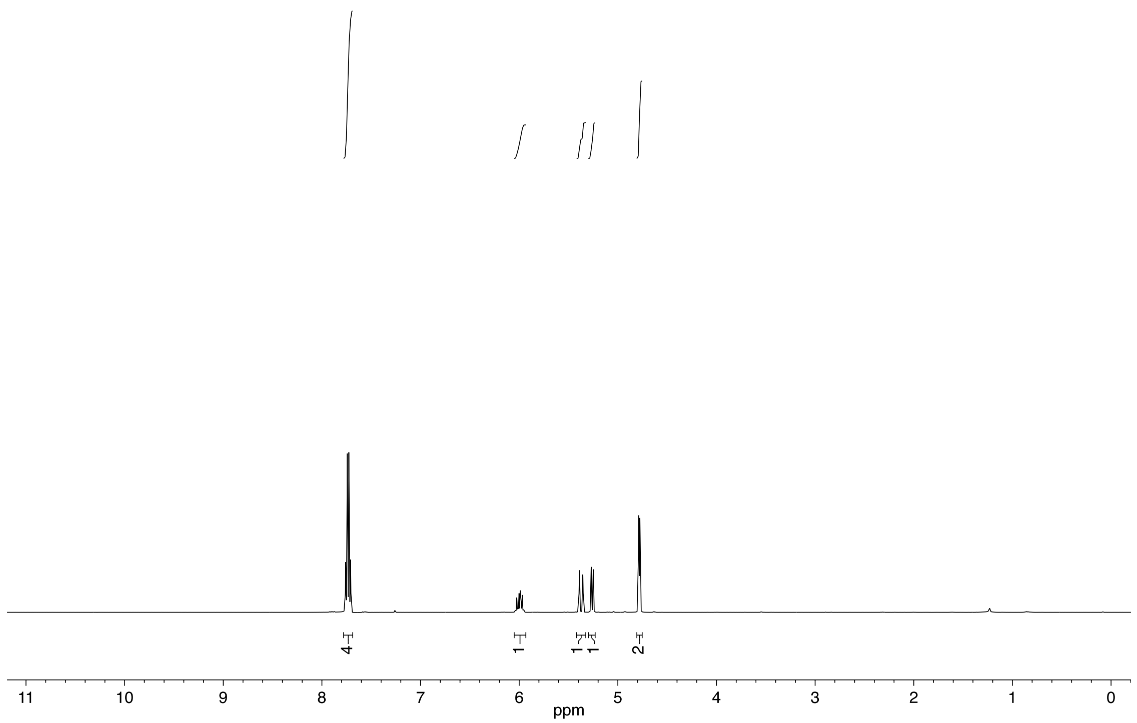


**(5-Iodo-1,3-phenylene)bis(morpholinomethanone) (19)**

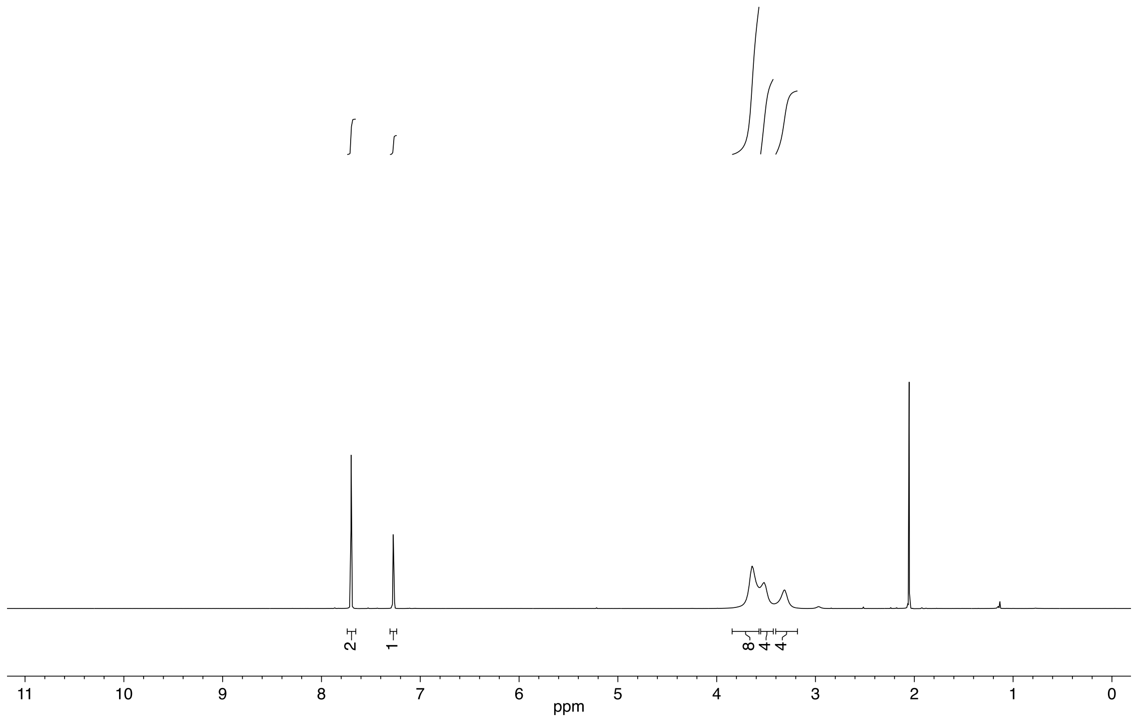

**5-Iodo-$N^1$,$N^1$,$N^3$,$N^3$-tetramethylisophthalamide (20)**

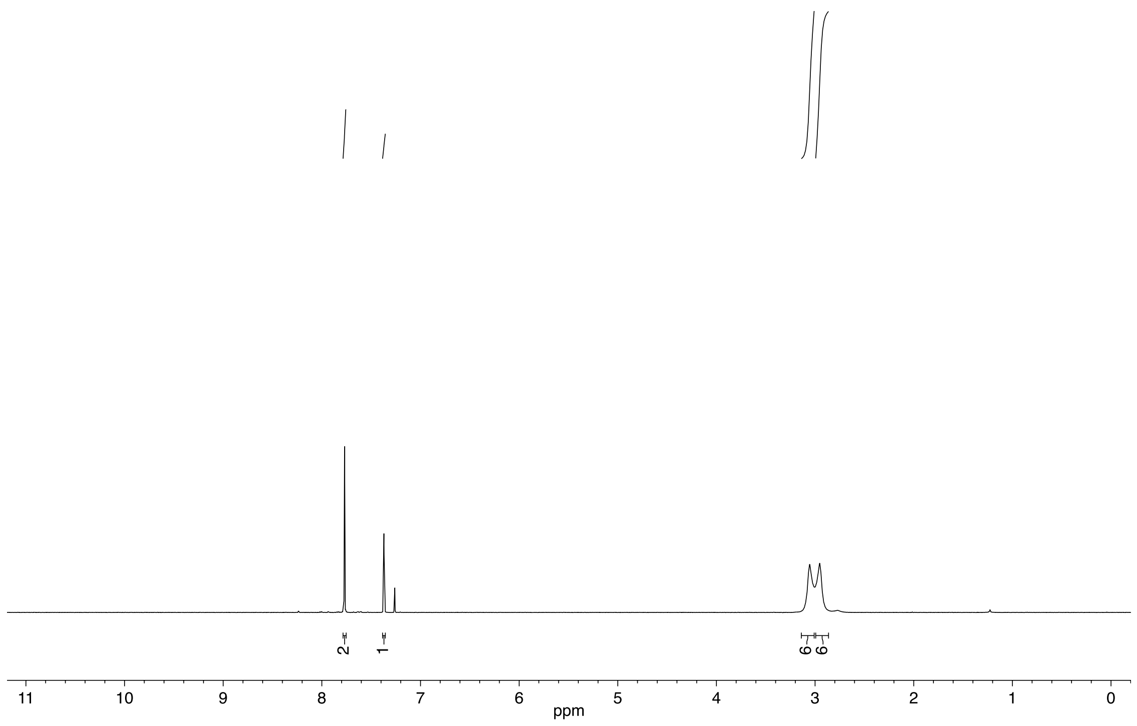

**Diallyl 5-iodoisophthalate (21)**

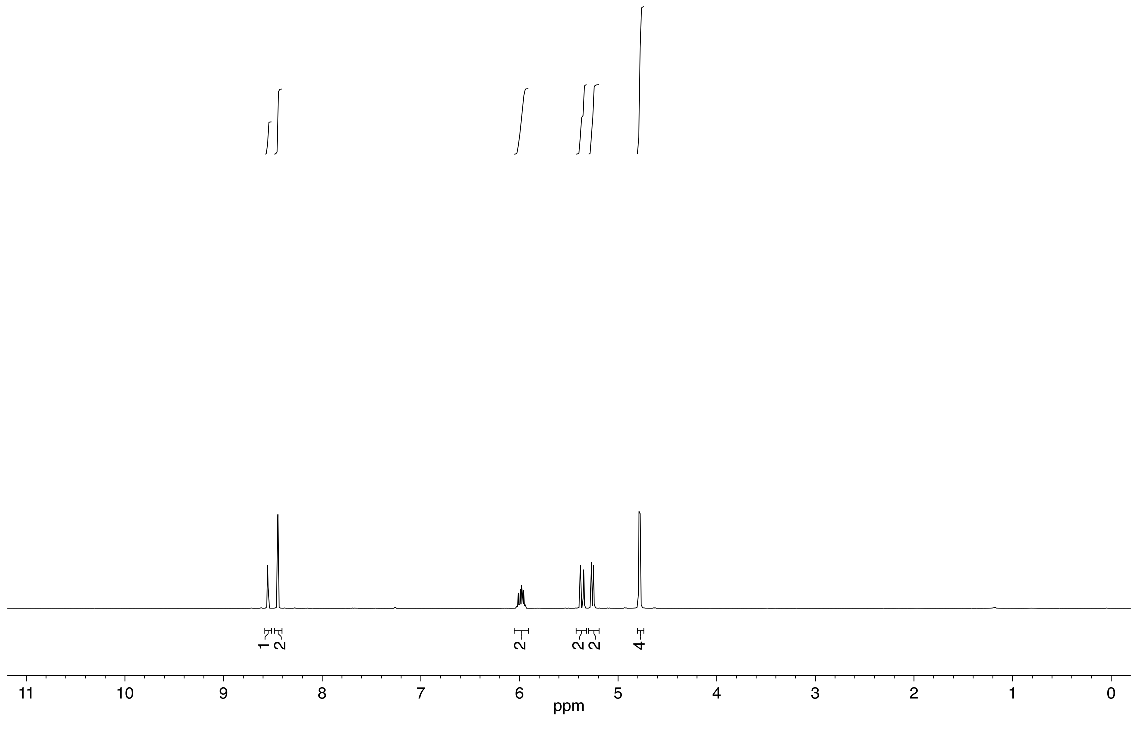

**Diethyl 1,1'-(5-iodoisophthaloyl)bis(piperidine-4-carboxylate) (22)**

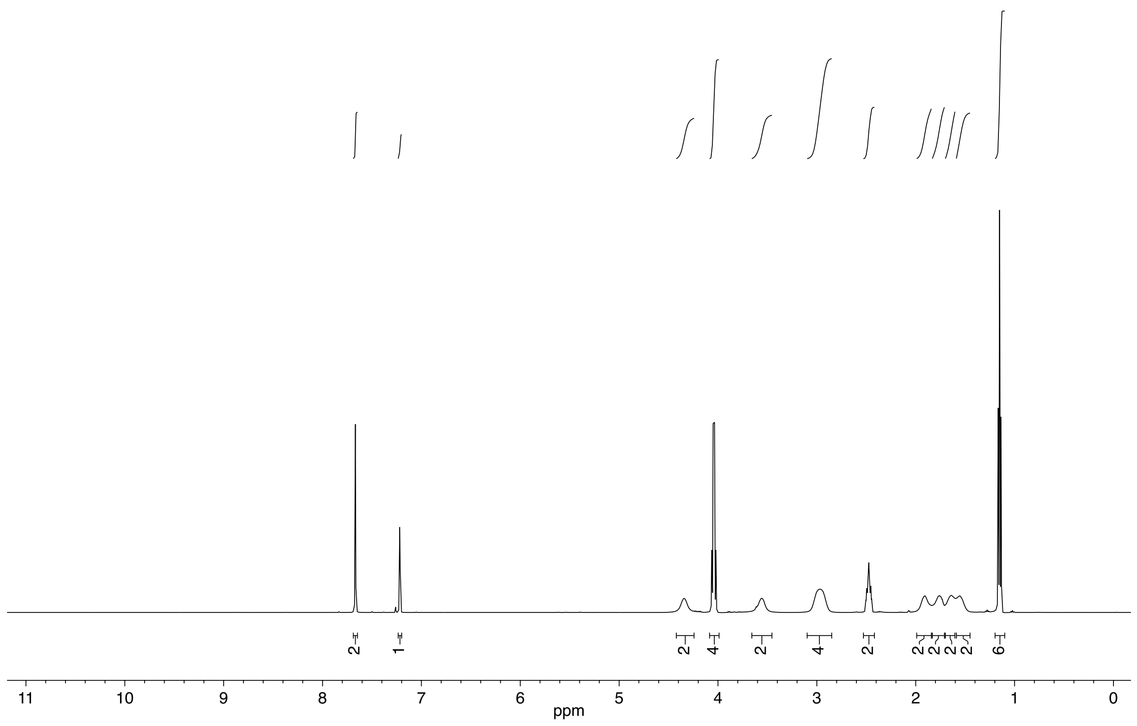

**(4-(Hydroxy(6-nitrobenzo[*d*][1,3]dioxol-5-yl)methyl)phenyl)(morpholino)methanone (23)**

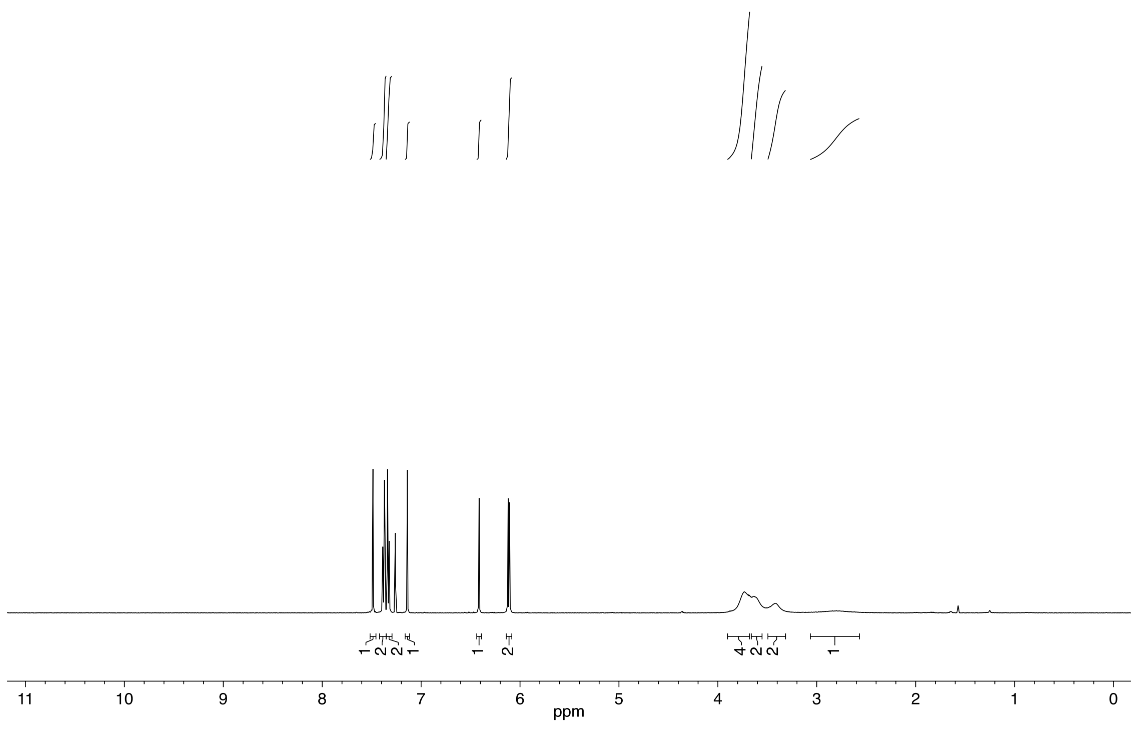

***N*-(*tert*-Butyl)-4-(hydroxy(6-nitrobenzo[*d*][1,3]dioxol-5-yl)methyl)benzamide (24)**

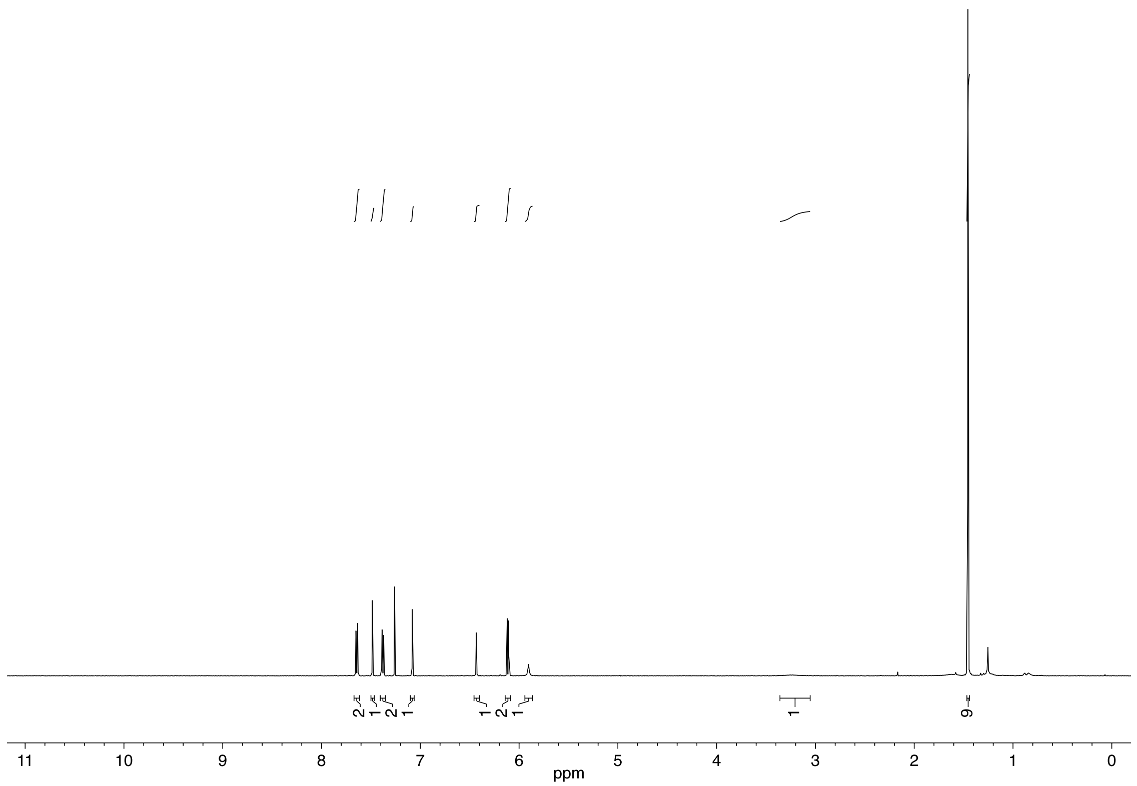

**Ethyl 1-(4-(hydroxy(6-nitrobenzo[*d*][1,3]dioxol-5-yl)methyl)benzoyl)piperidine-4-carboxylate (25)**

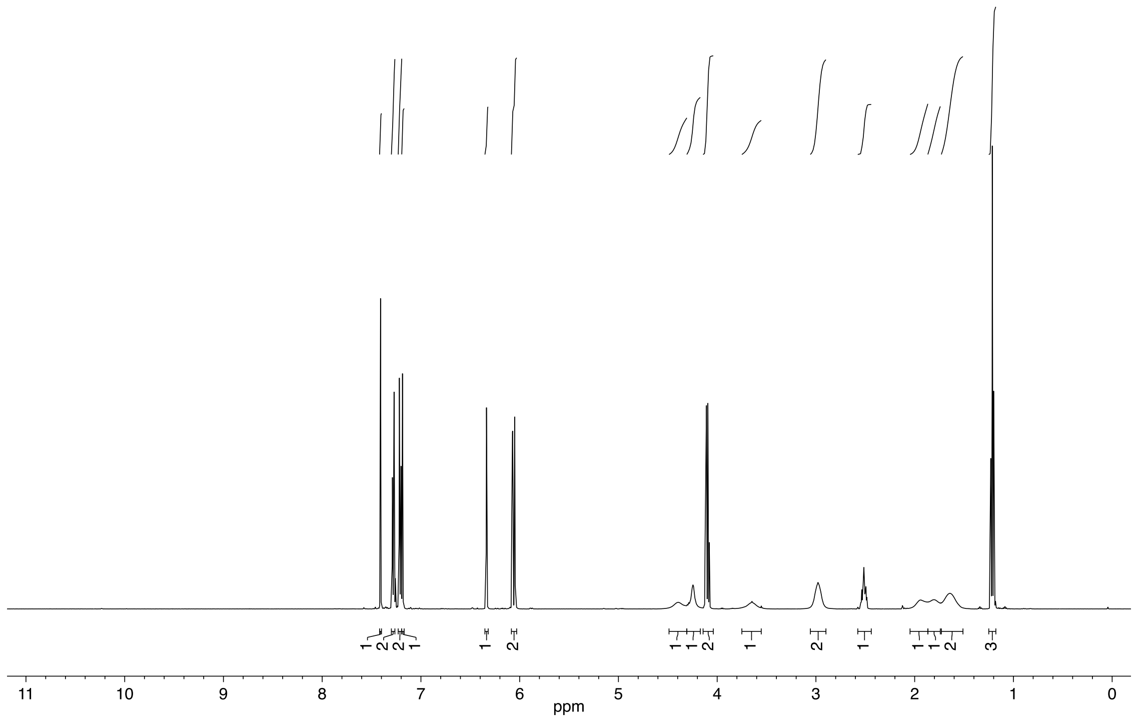

**5-(Hydroxy(6-nitrobenzo[*d*][1,3]dioxol-5-yl)methyl)-*N*[1],*N*[1],*N*[3],*N*[3]-tetramethylisophthalamide (26)**

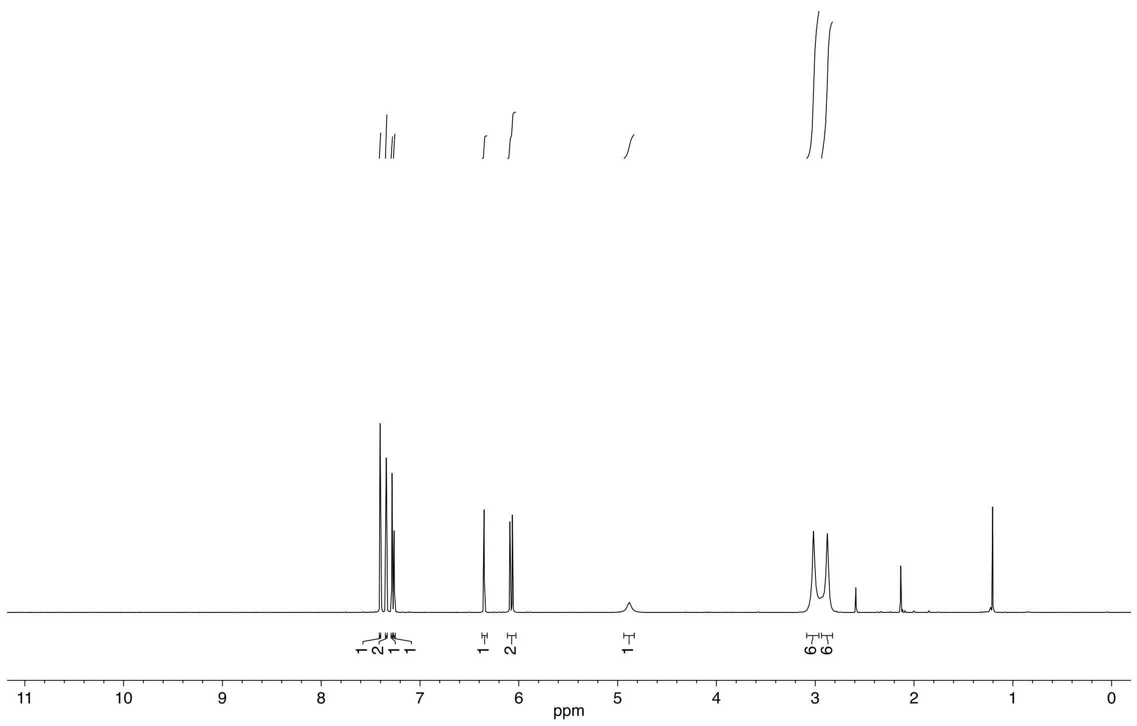


**(5-(Hydroxy(6-nitrobenzo[*d*][1,3]dioxol-5-yl)methyl)-1,3-phenylene)bis(morpholinomethanone) (27)**

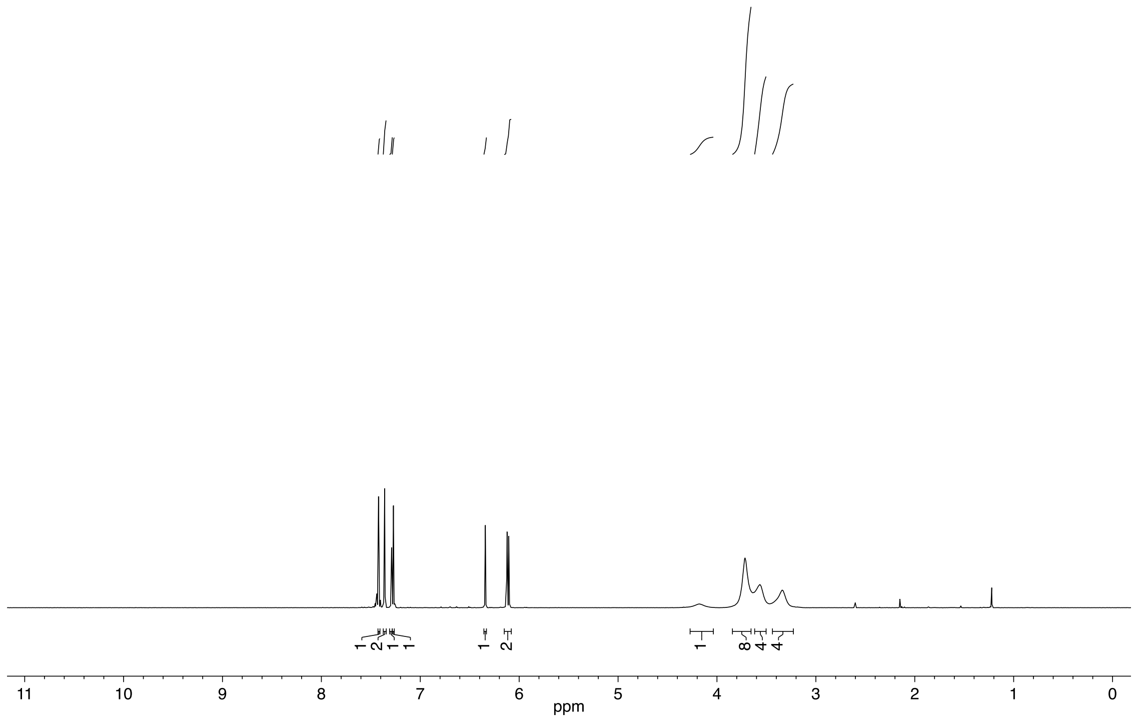

**Allyl 4-(hydroxy(6-nitrobenzo[*d*][1,3]dioxol-5-yl)methyl)benzoate (28)**

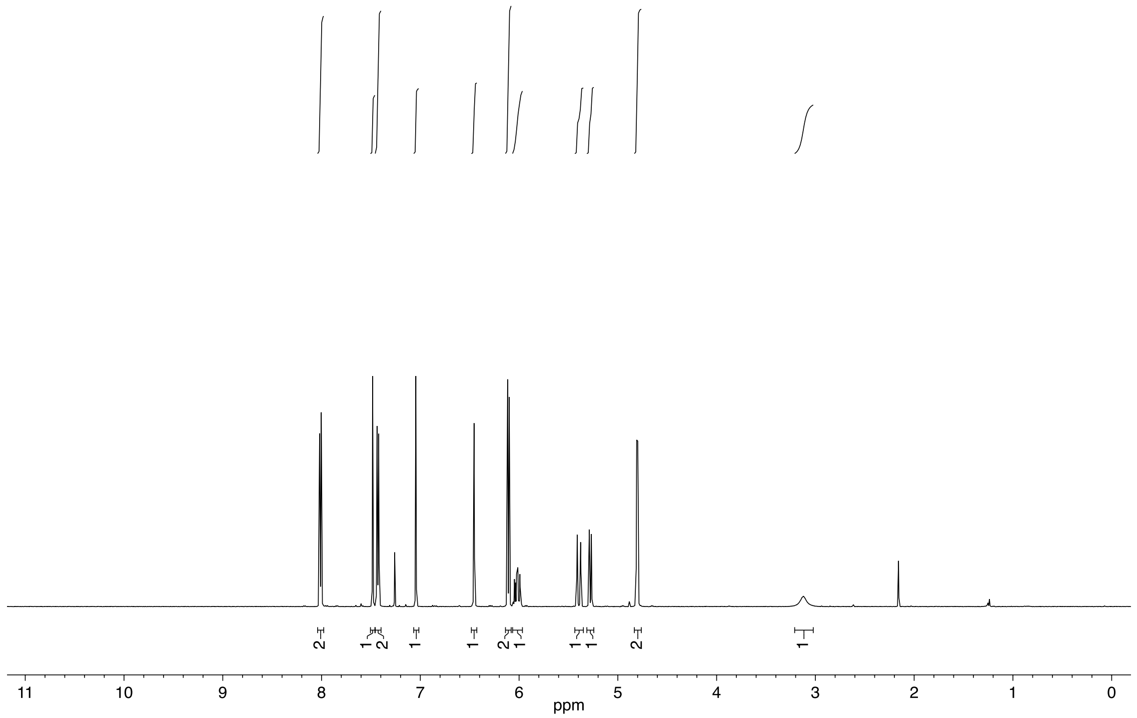


**Diallyl 5-(hydroxy(6-nitrobenzo[*d*][1,3]dioxol-5-yl)methyl)isophthalate (29)**

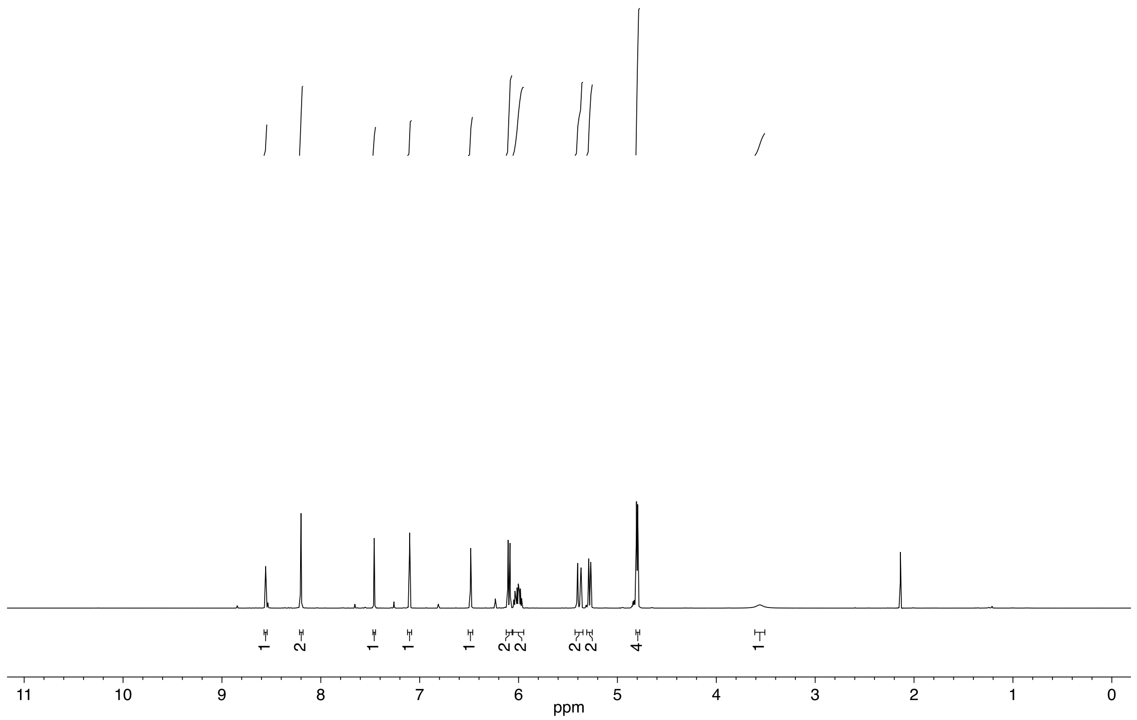

**Diethyl 1,1'-(5-(hydroxy(6-nitrobenzo[*d*][1,3]dioxol-5-yl)methyl)isophthaloyl)bis(piperidine-4-carboxylate) (30)**

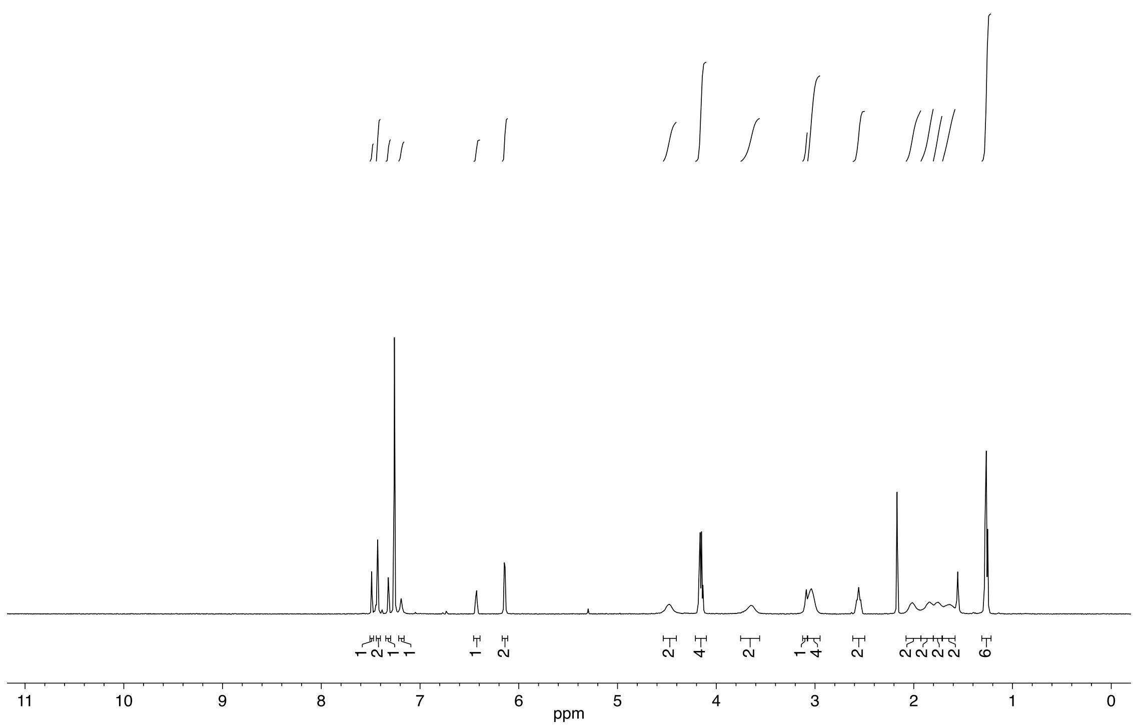

**Diallyl 1,1'-(5-(hydroxy(6-nitrobenzo[*d*][1,3]dioxol-5-yl)methyl)isophthaloyl)bis(piperidine-4-carboxylate) (31)**

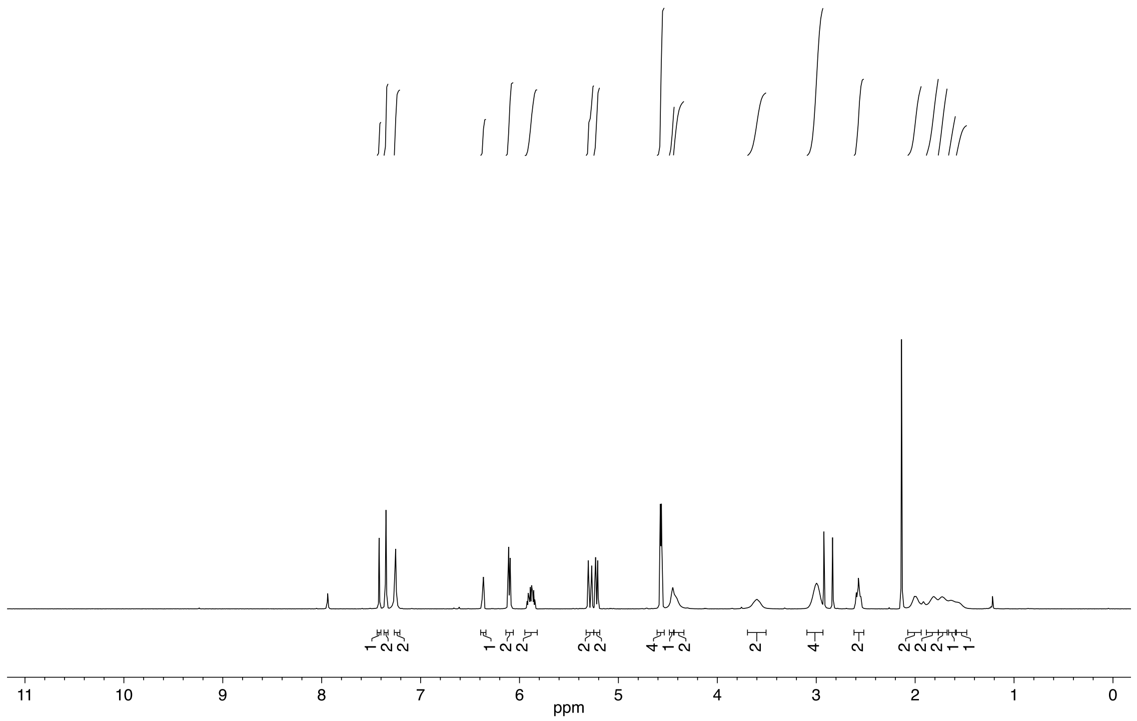

**(1-(6-Nitrobenzo[*d*][1,3]dioxol-5-yl)ethyl) succinimid-N-yl carbonate (32)**

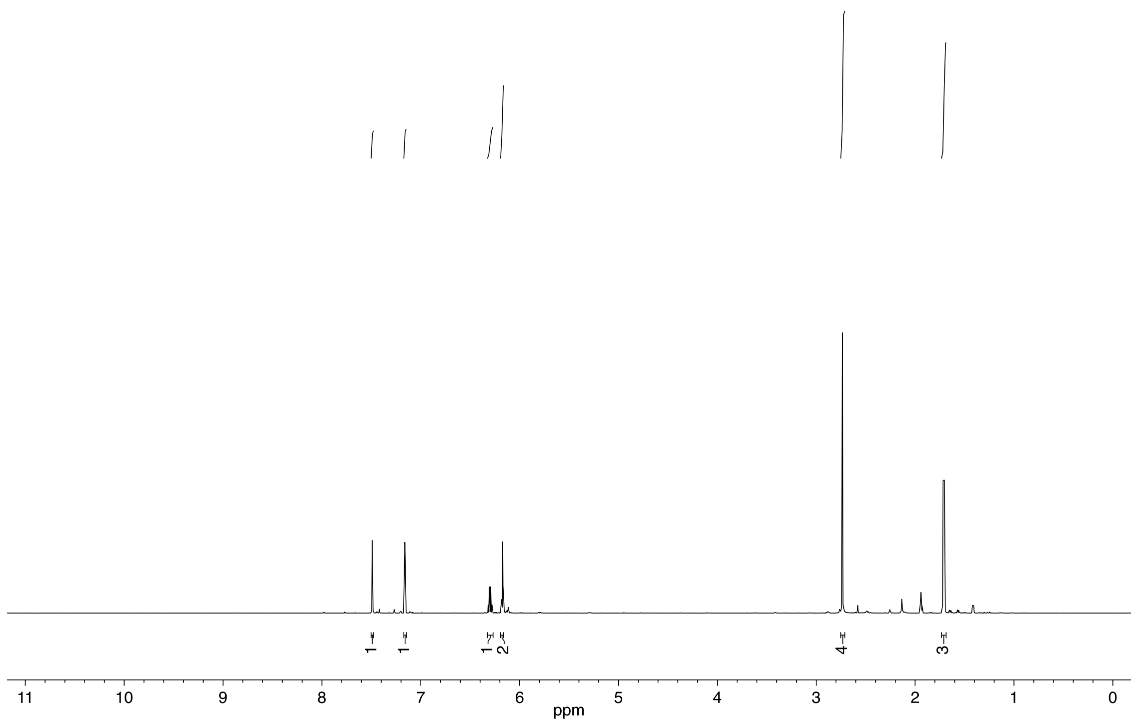


**((4-(Morpholine-4-carbonyl)phenyl)(6-nitrobenzo[*d*][1,3]dioxol-5-yl)methyl) succinimid-N-yl cabonate (33)**

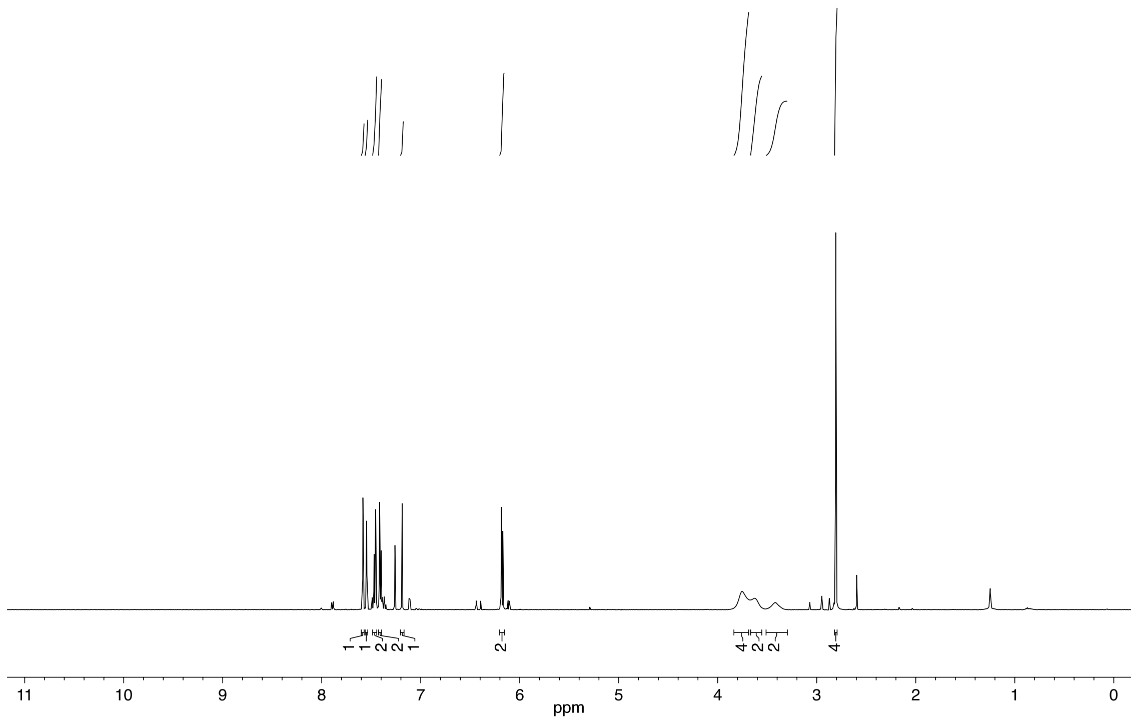

**(4-(*tert*-Butylcarbamoyl)phenyl)(6-nitrobenzo[*d*][1,3]dioxol-5-yl)methyl succinimid-N-yl carbonate (34)**

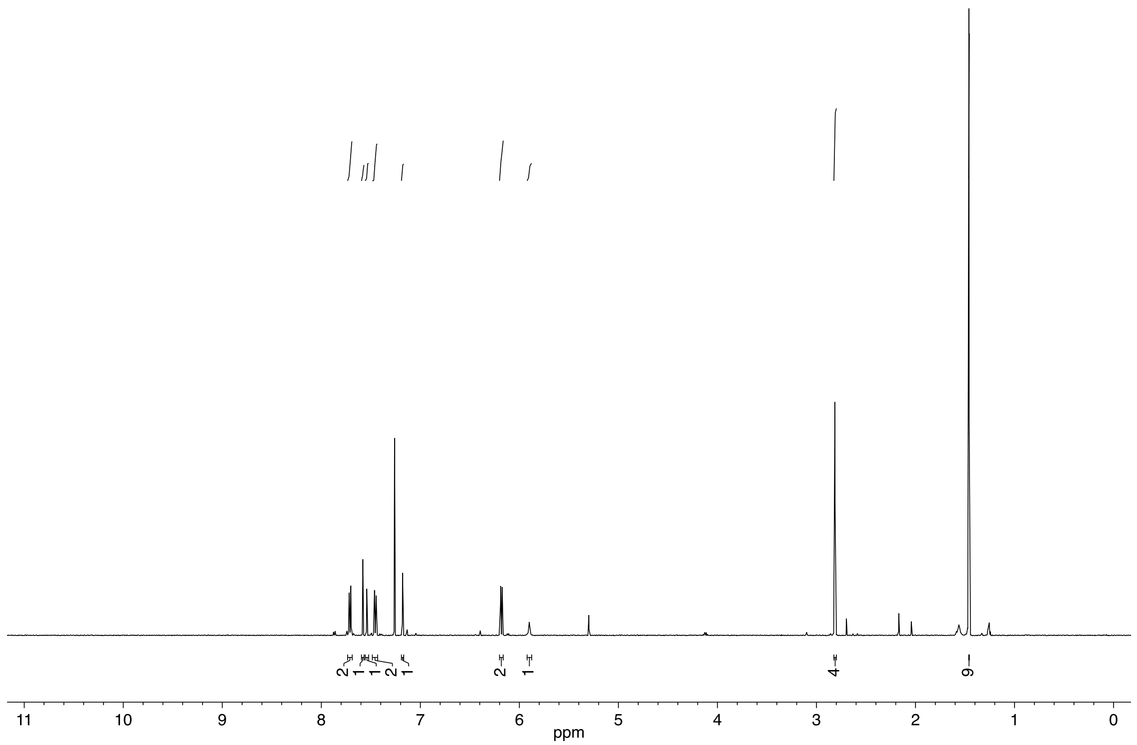

**(4-(4-(Ethylcarbonoyl)piperidine-N-carbonyl)phenyl)(6-nitrobenzo[*d*][1,3]dioxol-5-yl)methyl succinimid-N-yl carbonate (35)**

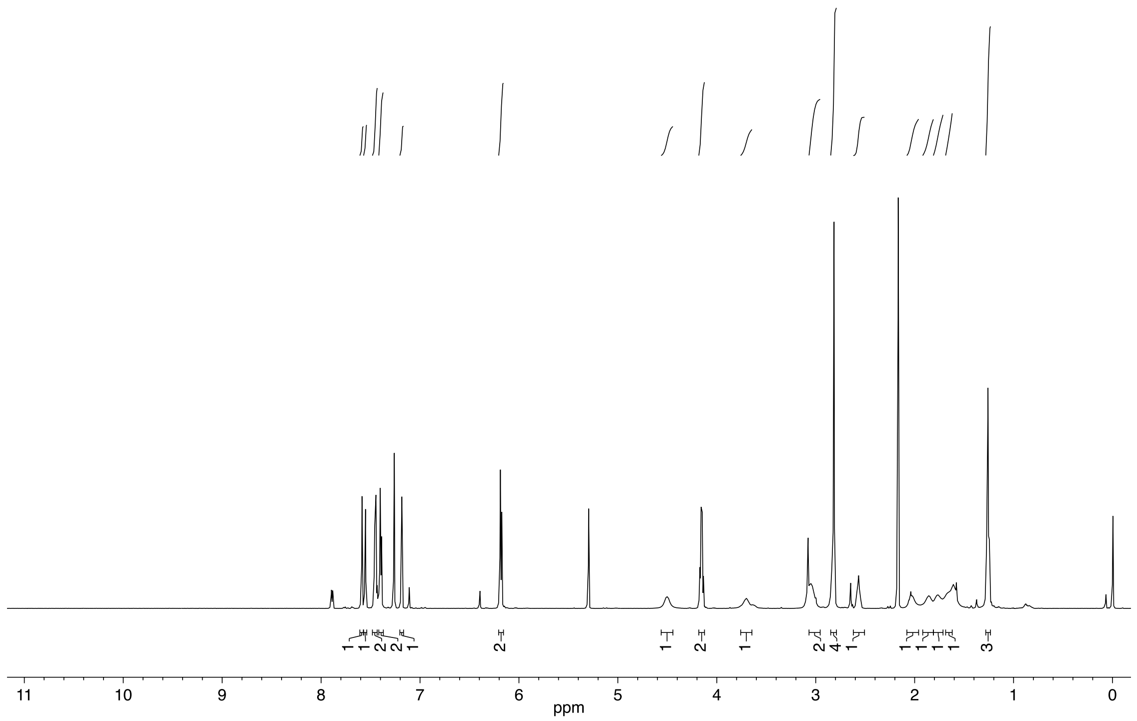

**(4-(Allylcarbonoyl)phenyl)(6-nitrobenzo[*d*][1,3]dioxol-5-yl)methyl succinimid-N-yl carbonate (36)**

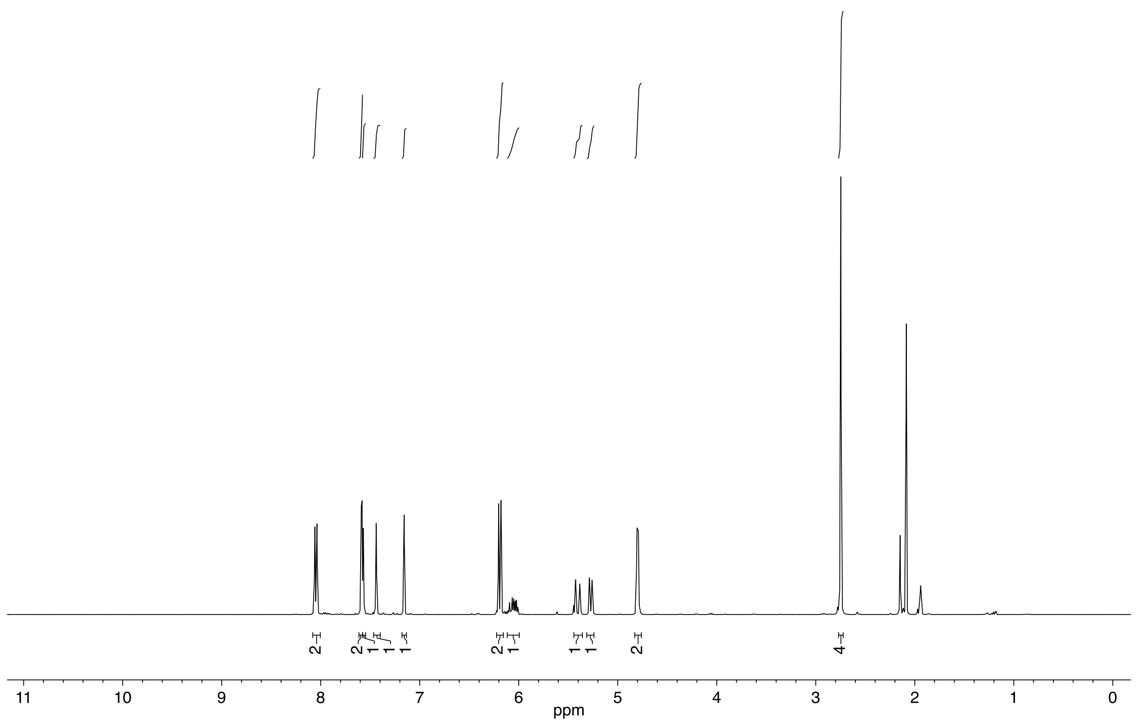


**(4-(Carbonoyl)phenyl)(6-nitrobenzo[*d*][1,3]dioxol-5-yl)methyl succinimid-N-yl carbonate (37)**

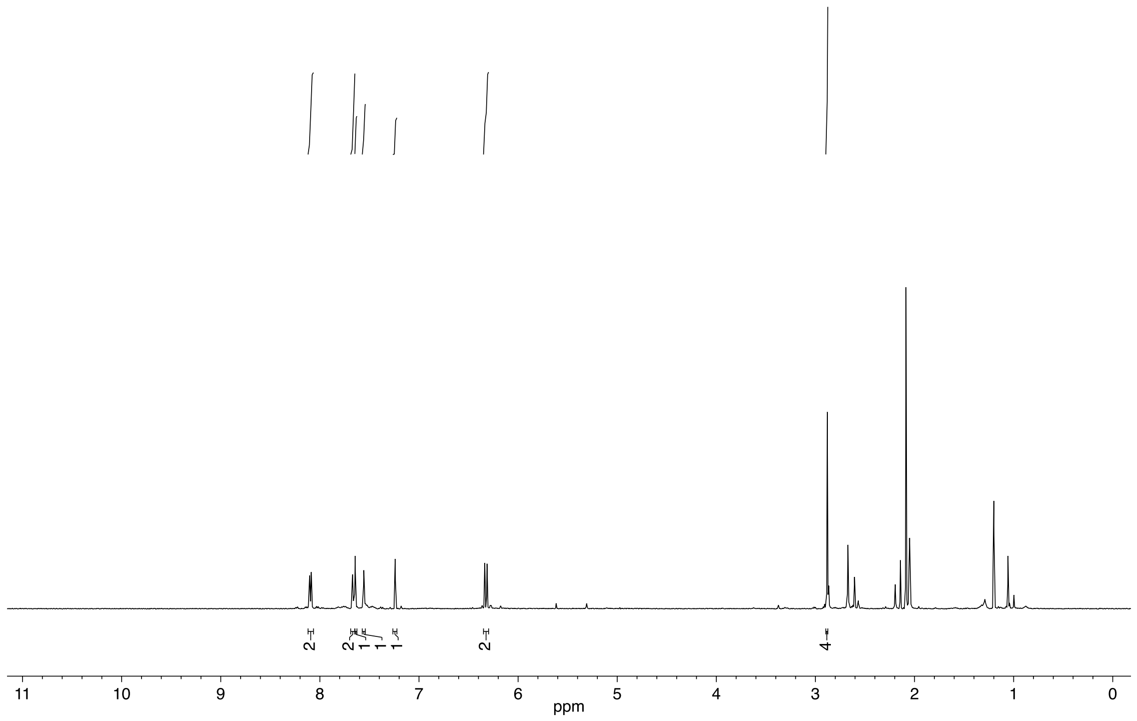

**(3,5-Bis(dimethylcarbamoyl)phenyl)(6-nitrobenzo[*d*][1,3]dioxol-5-yl)methyl succinimid-N-yl carbonate (38)**

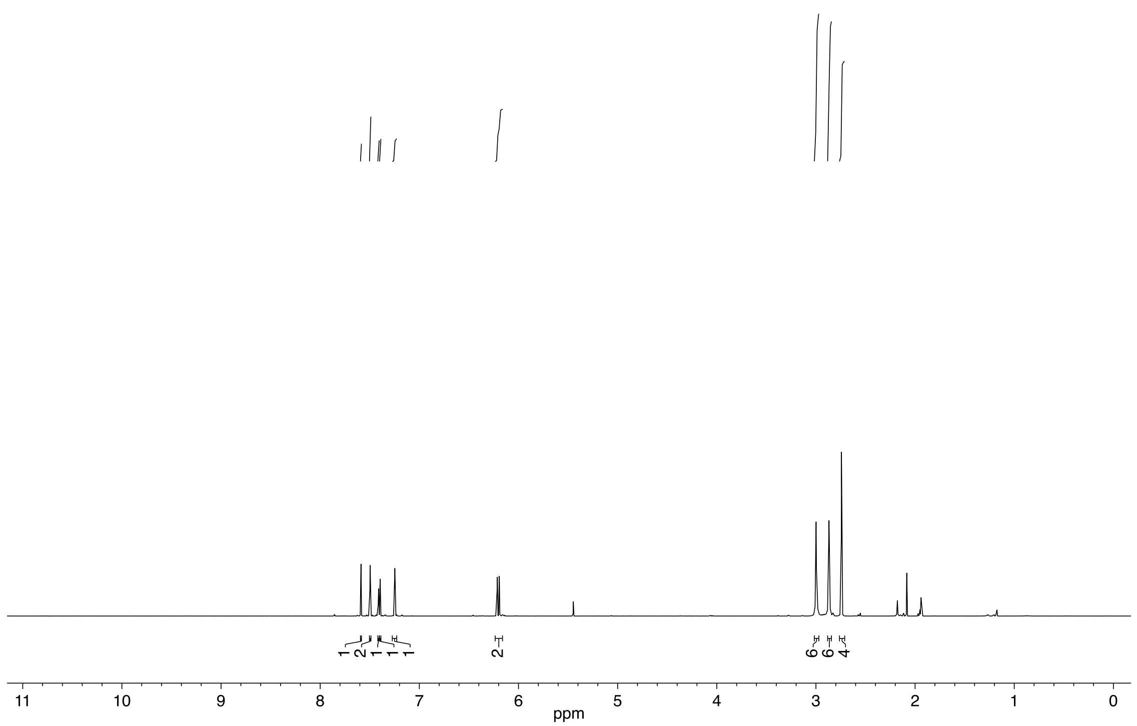


**(3,5-Di(morpholine-4-carbonyl)phenyl)(6-nitrobenzo[*d*][1,3]dioxol-5-yl)methyl succinimid-N-yl carbonate (39)**

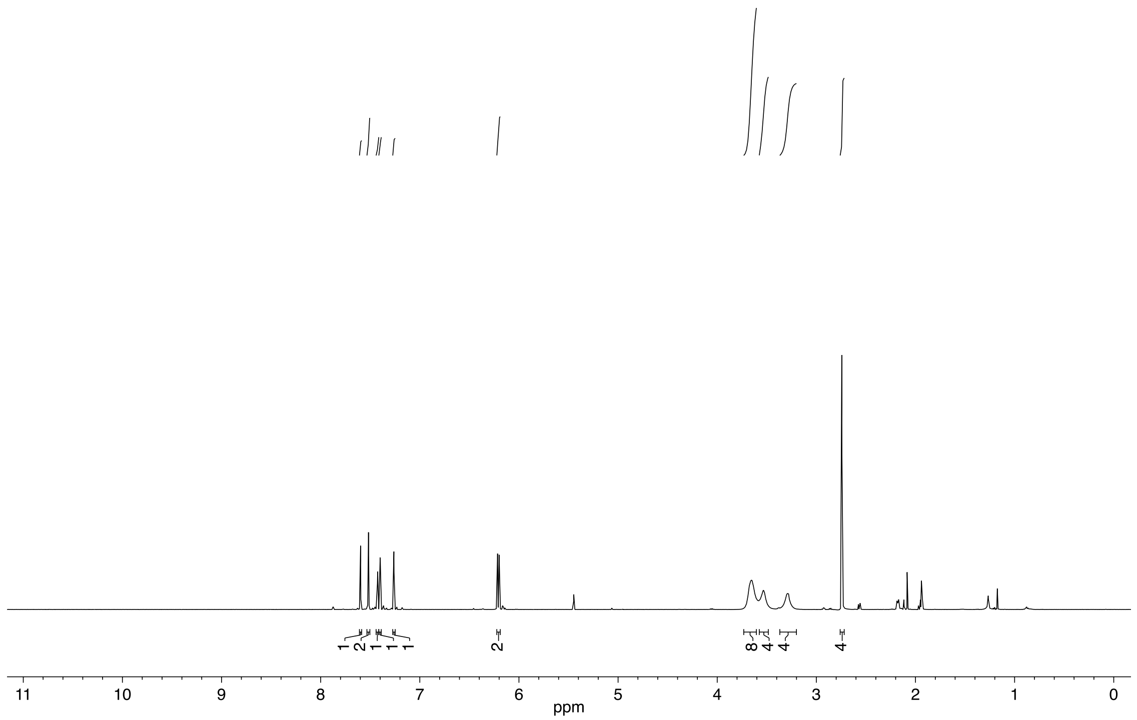

**(3,5-Di(allylcarbonoyl)phenyl)(6-nitrobenzo[d][1,3]dioxol-5-yl)methyl succinimid-N-yl carbonate (40)**

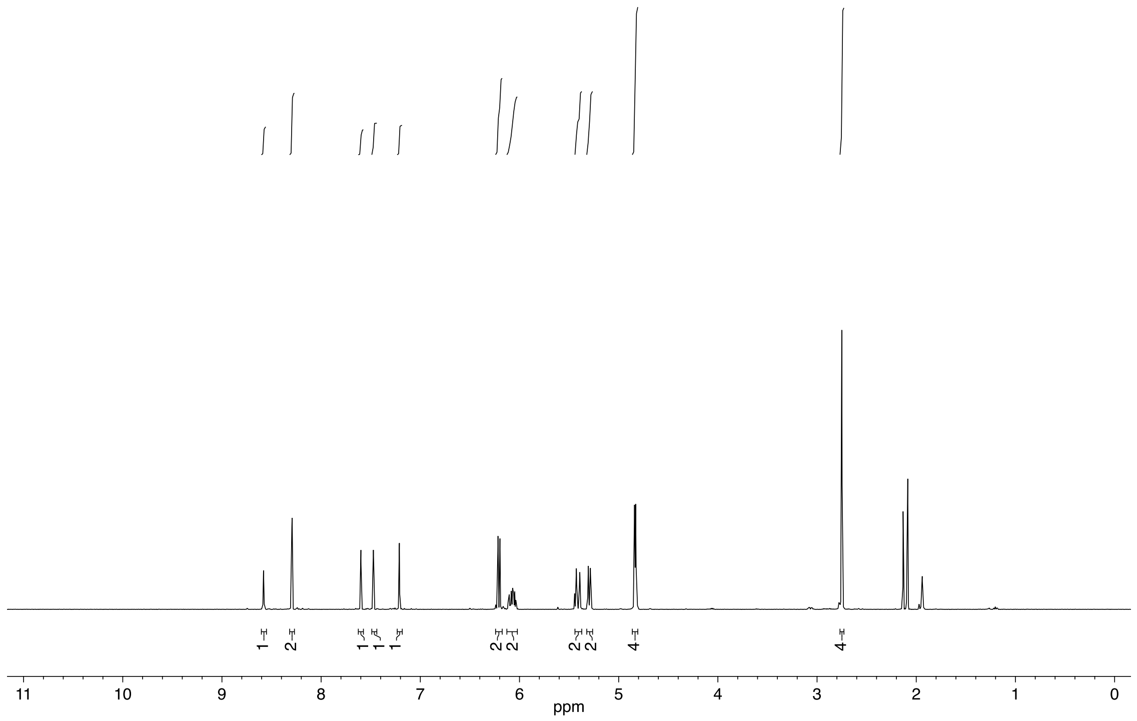


**(3,5-Dicarbonoylphenyl)(6-nitrobenzo[d][1,3]dioxol-5-yl)methyl succinimid-N-yl carbonate (41)**

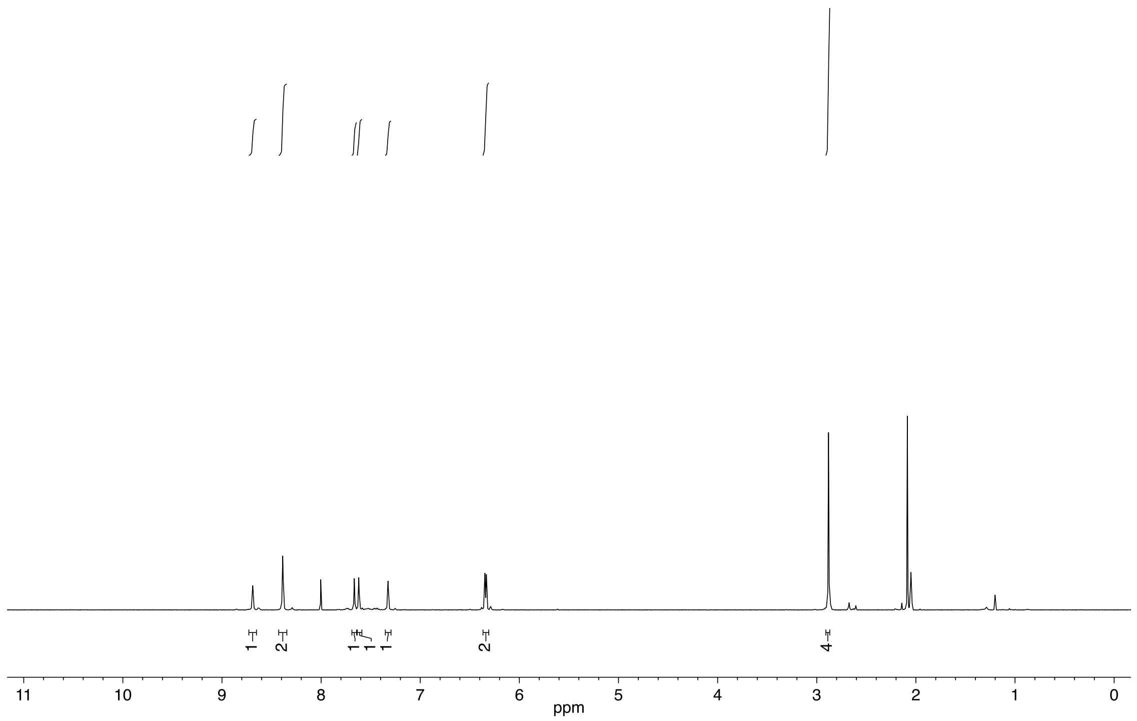

**(3,5-Di(4-(allylcarbonoyl)piperidine-N-carbonyl)phenyl)(6-nitrobenzo[*d*][1,3]dioxol-5-yl)methyl succinimid-N-yl carbonate (42)**

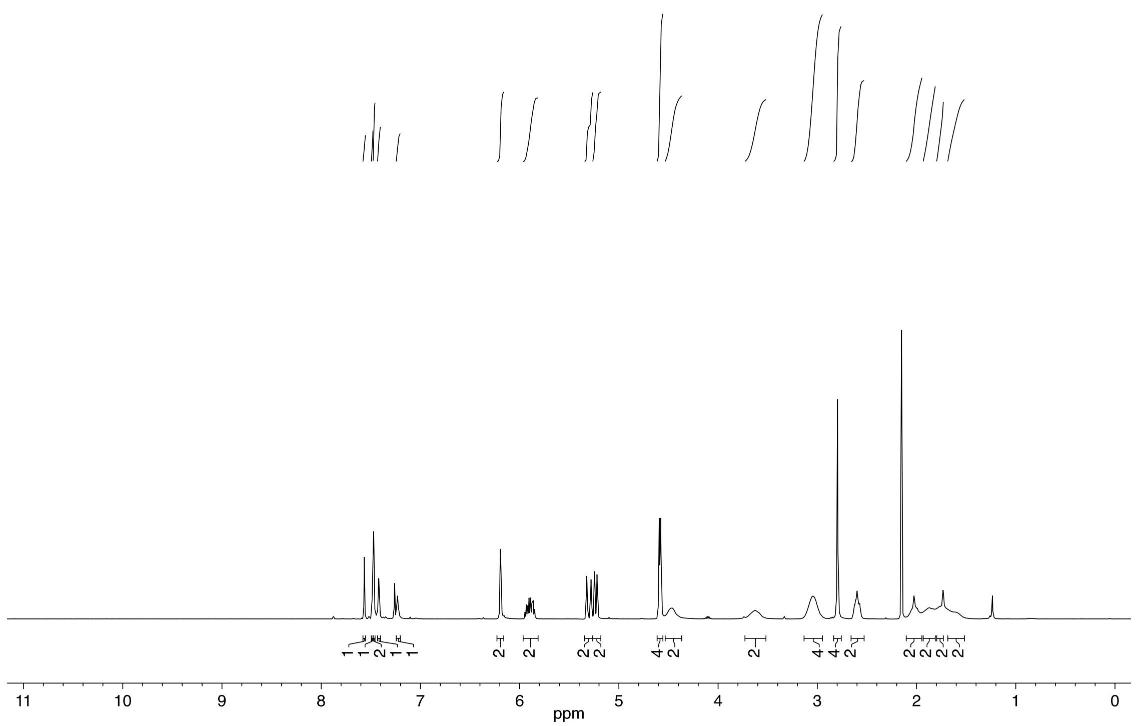

**(3,5-Di(4-carbonoylpiperidine-N-carbonyl)phenyl)(6-nitrobenzo[*d*][1,3]dioxol-5-yl)methyl succinimid-N-yl carbonate (43)**

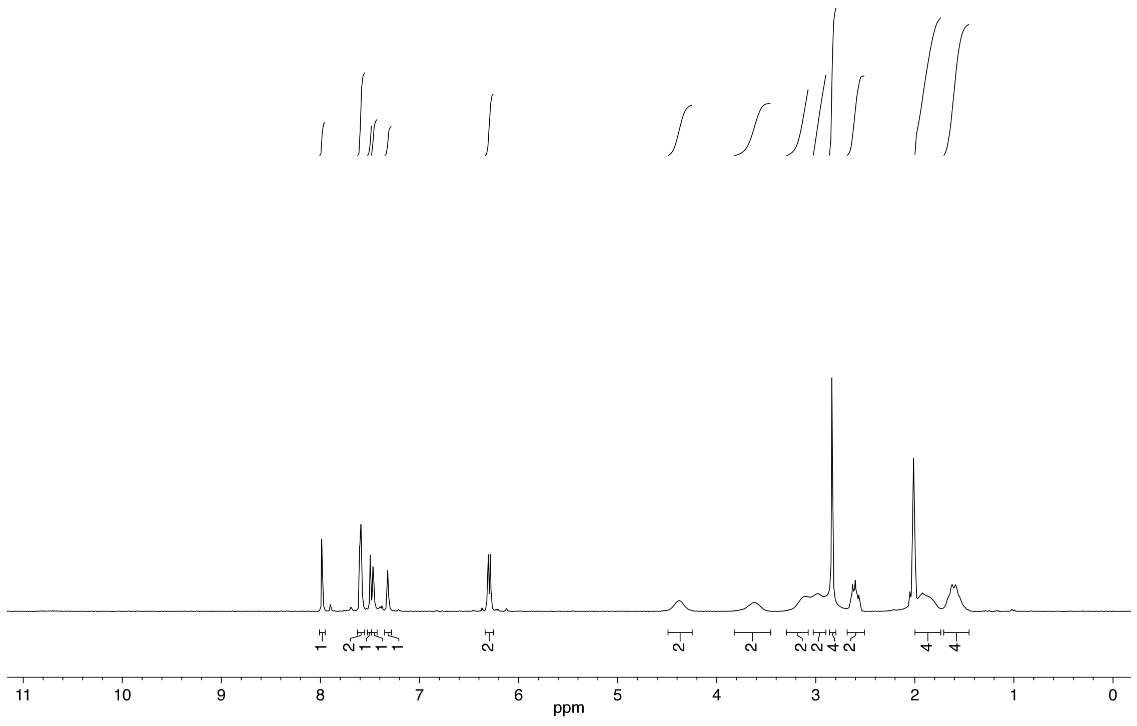

**(1-(6-Nitrobenzo[*d*][1,3]dioxol-5-yl)ethyl) saxitoxin N21-ethylcarbamate (2)**

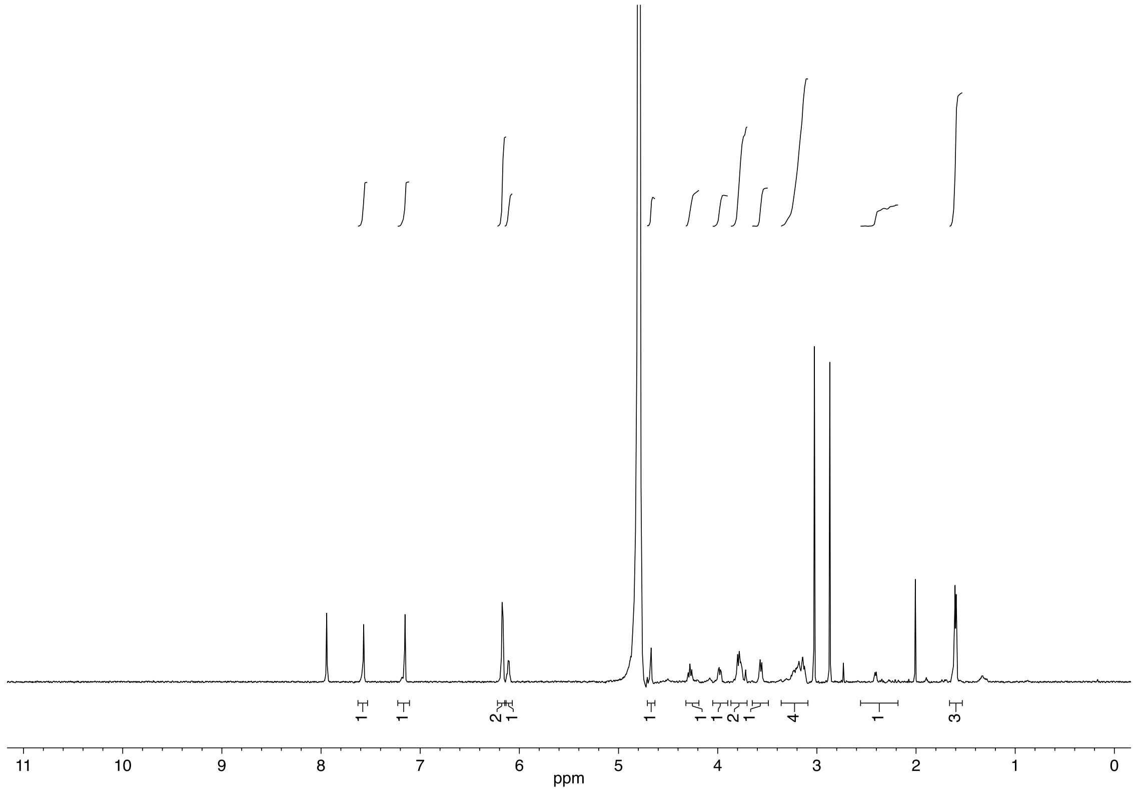


**((4-(Morpholine-4-carbonyl)phenyl)(6-nitrobenzo[*d*][1,3]dioxol-5-yl)methyl) saxitoxin N21-ethylcarbamate (3)**

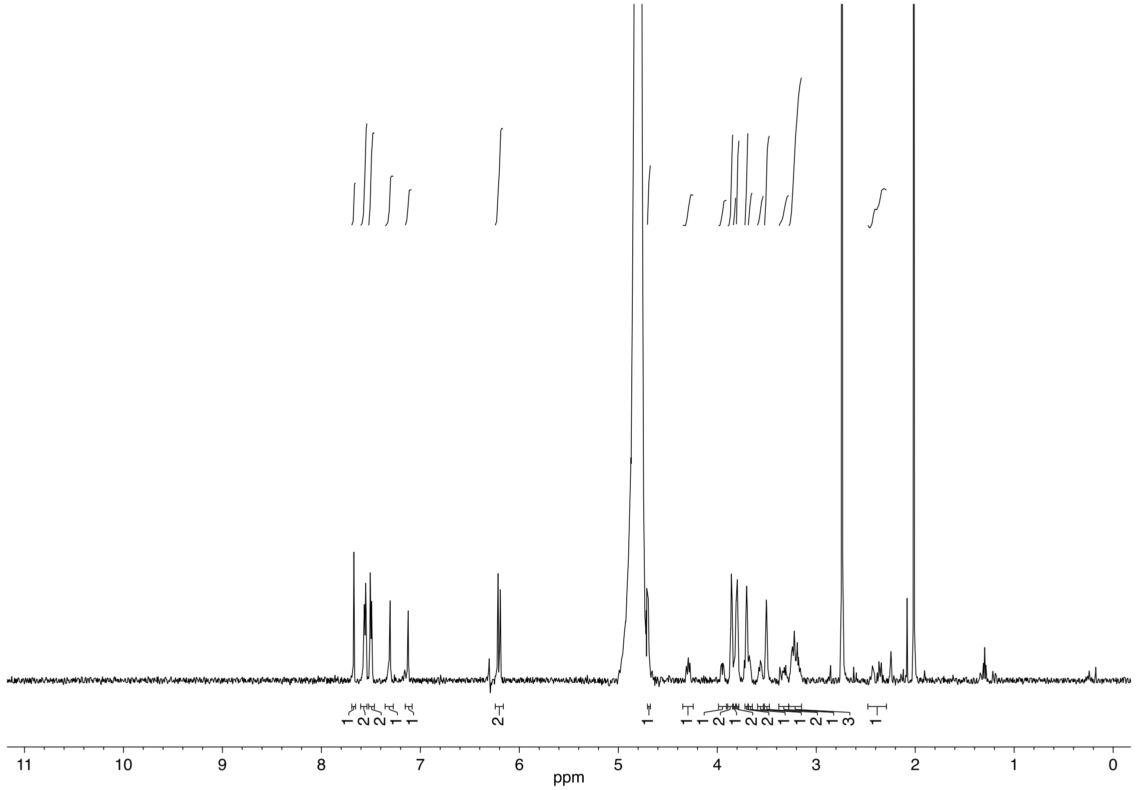

**(3,5-Di(morpholine-4-carbonyl)phenyl)(6-nitrobenzo[*d*][1,3]dioxol-5-yl)methyl axitoxin N21-ethyl-carbamate (4)**

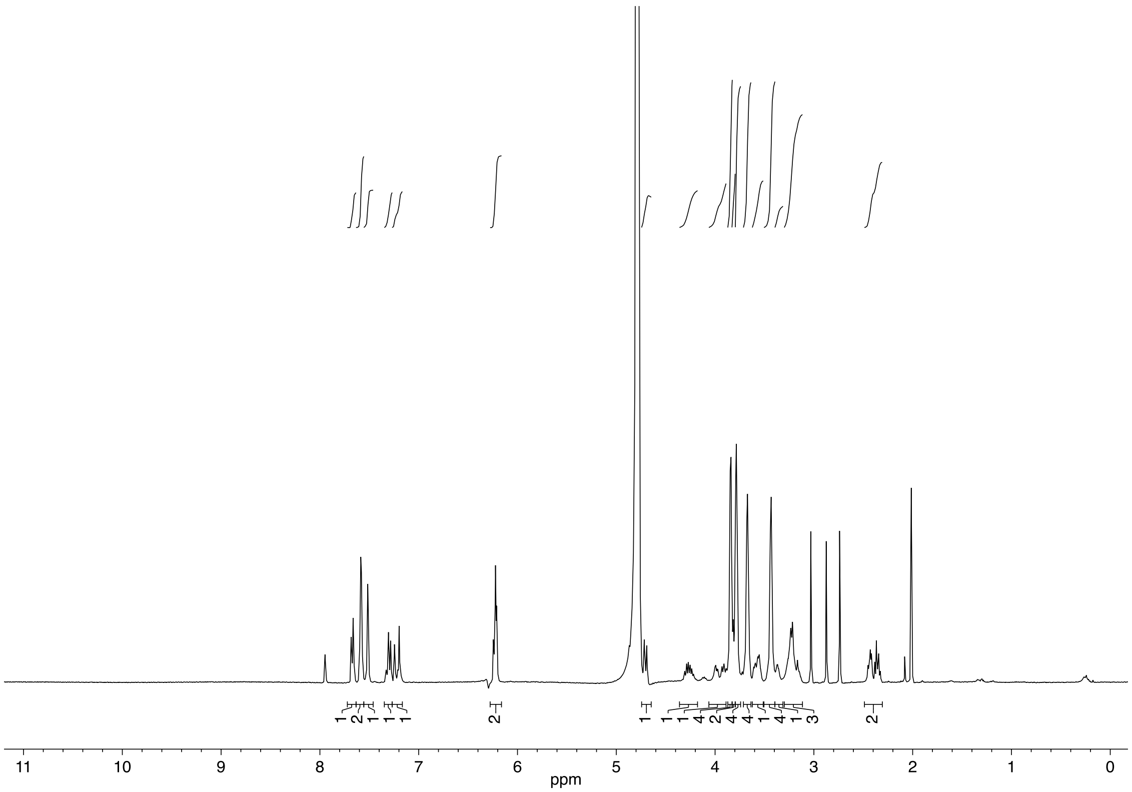


**(4-(*tert*-Butylcarbamoyl)phenyl)(6-nitrobenzo[*d*][1,3]dioxol-5-yl)methyl saxitoxin N21-ethylcarba-mate (5)**

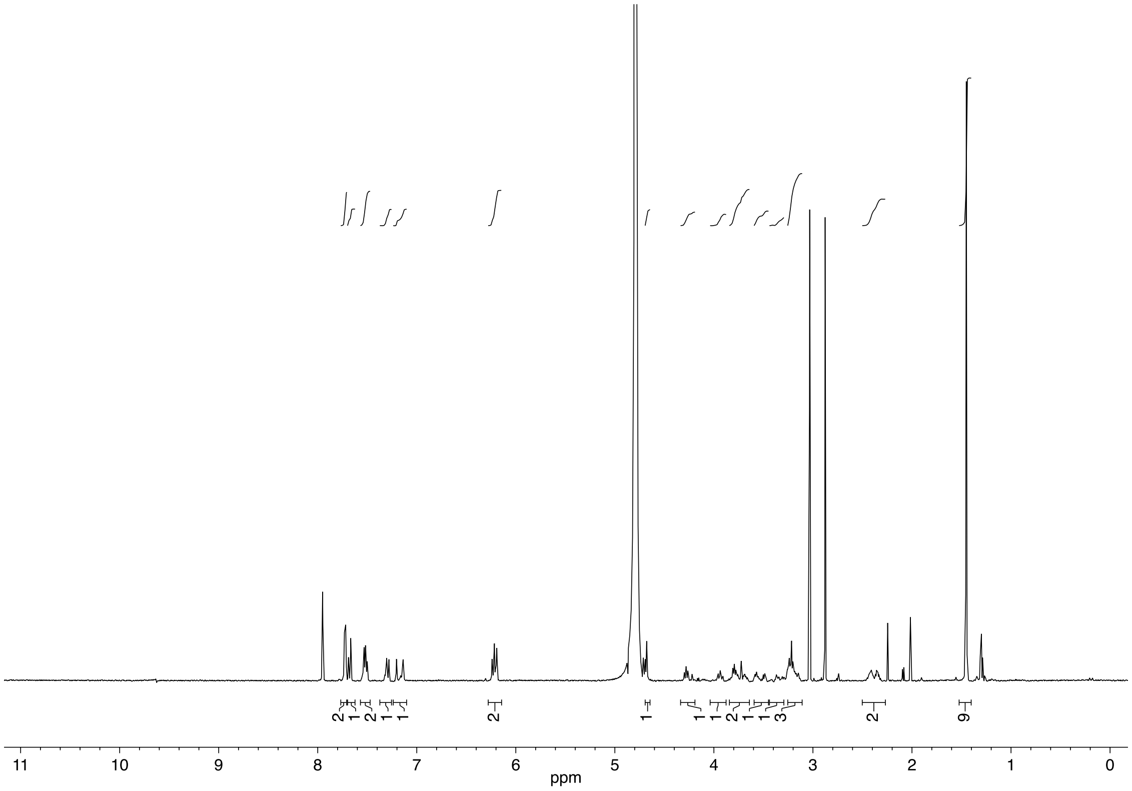

**(4-(4-(Ethylcarbonoyl)piperidine-N-carbonyl)phenyl)(6-nitrobenzo[*d*][1,3]dioxol-5-yl)methyl saxitoxin N21-ethylcarbamate (6)**

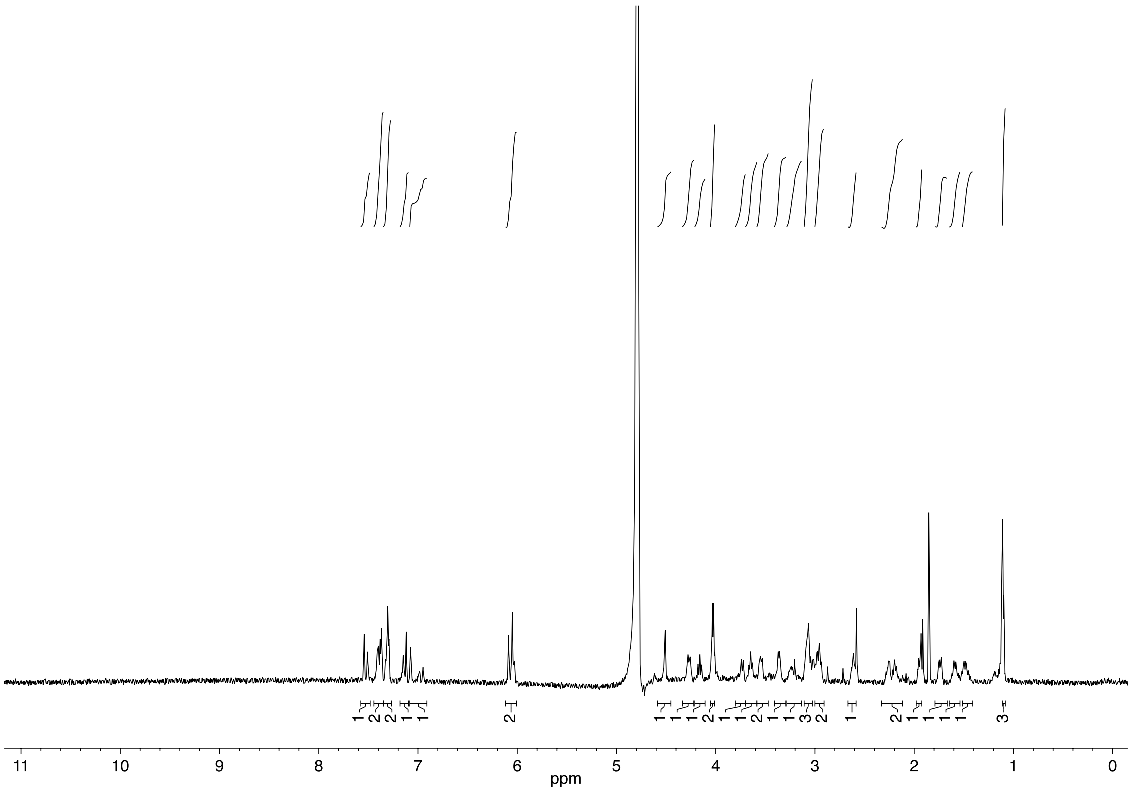


**(3,5-Bis(dimethylcarbamoyl)phenyl)(6-nitrobenzo[*d*][1,3]dioxol-5-yl)methyl saxitoxin N21-ethylcarbamate (7)**

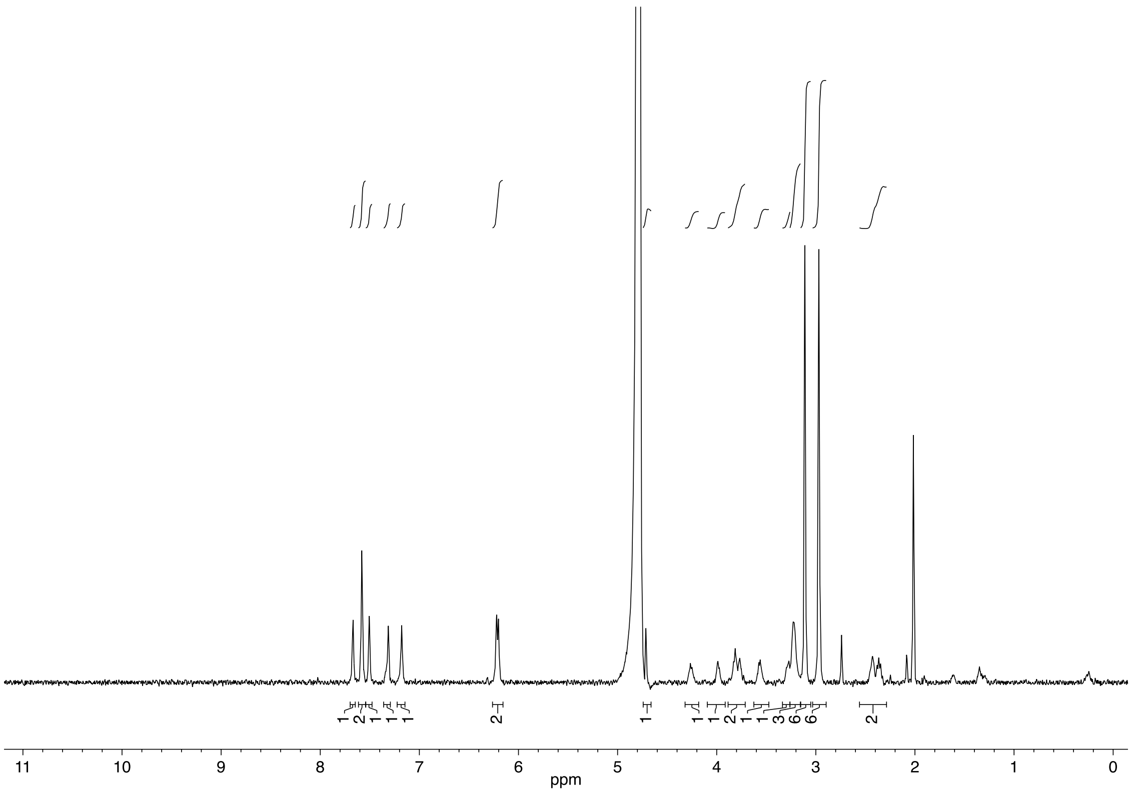

**(4-(Allylcarbonoyl)phenyl)(6-nitrobenzo[*d*][1,3]dioxol-5-yl)methyl saxitoxin N21-ethylcarbamate (8)**

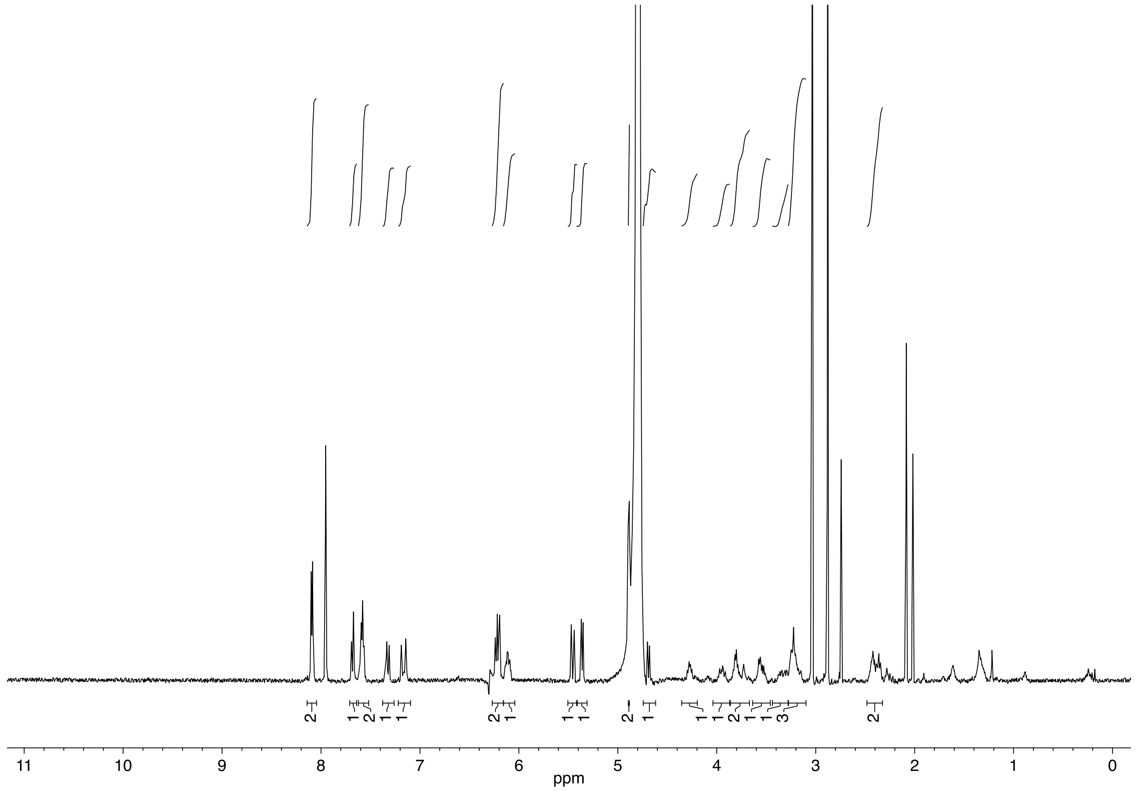

**(3,5-Di(allylcarbonoyl)phenyl)(6-nitrobenzo[d][1,3]dioxol-5-yl)methyl saxitoxin N21-ethylcarbamate (9)**

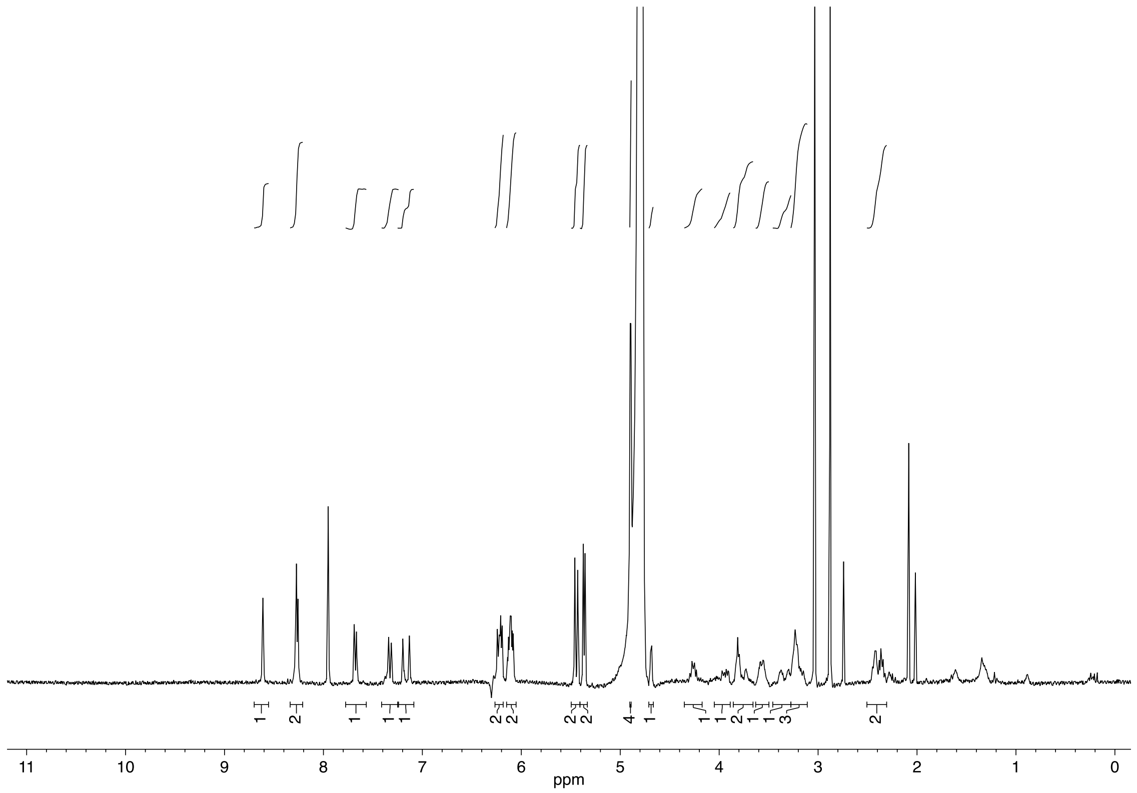

**(3,5-Di(4-(allylcarbonoyl)piperidine-N-carbonyl)phenyl)(6-nitrobenzo[*d*][1,3]dioxol-5-yl)methyl saxitoxin N21-ethylcarbamate (10)**

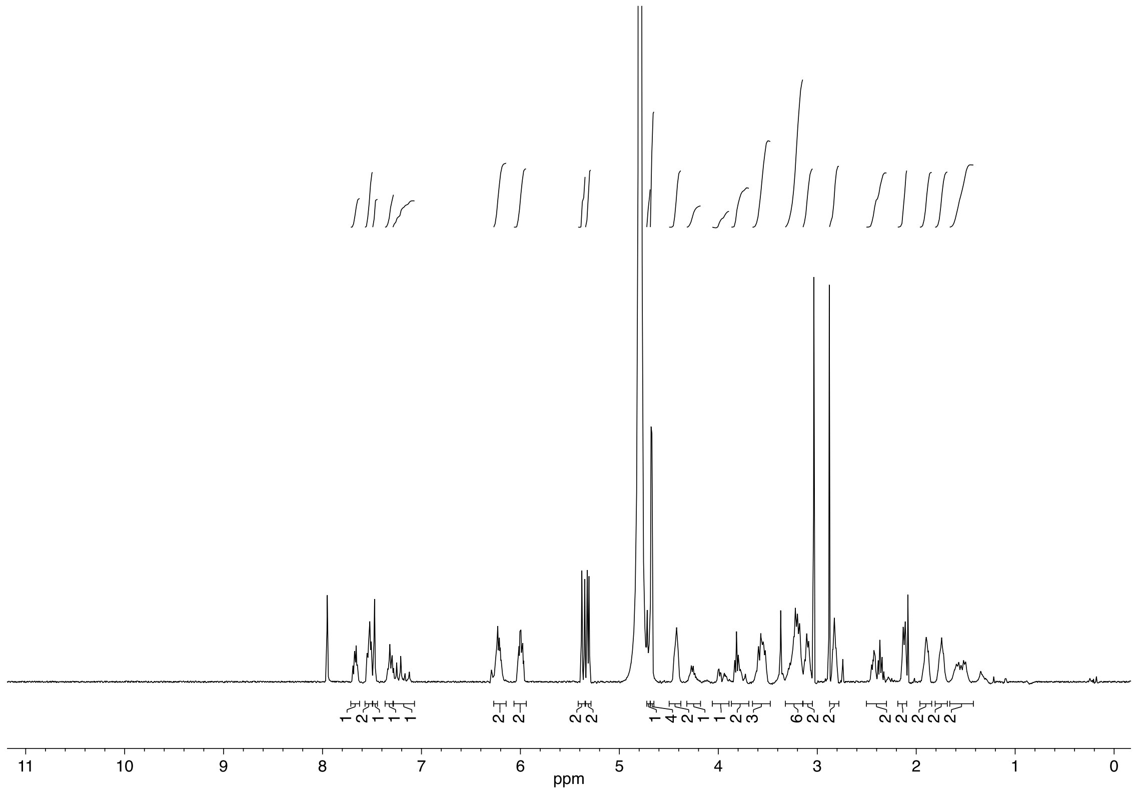


**(4-(Carbonoyl)phenyl)(6-nitrobenzo[*d*][1,3]dioxol-5-yl)methyl saxitoxin N21-ethylcarbamate (11)**

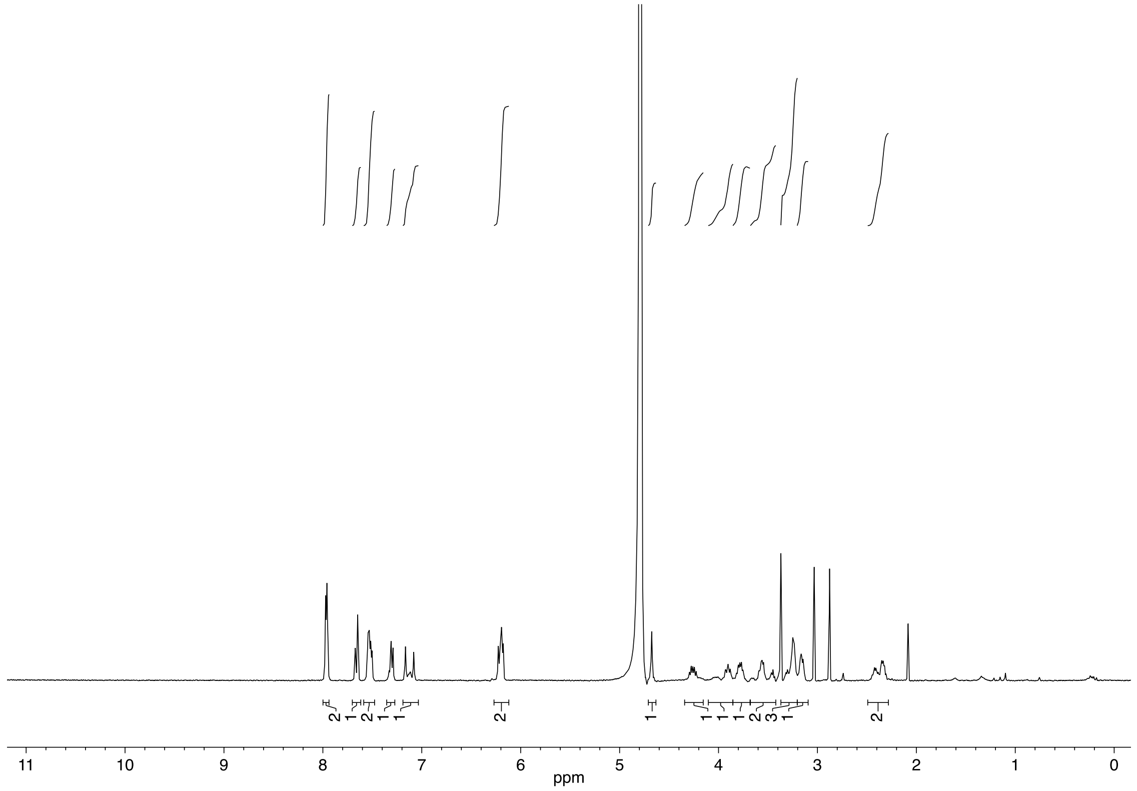

**(3,5-Dicarbonoylphenyl)(6-nitrobenzo[d][1,3]dioxol-5-yl)methyl saxitoxin N21-ethylcarbamate (12)**

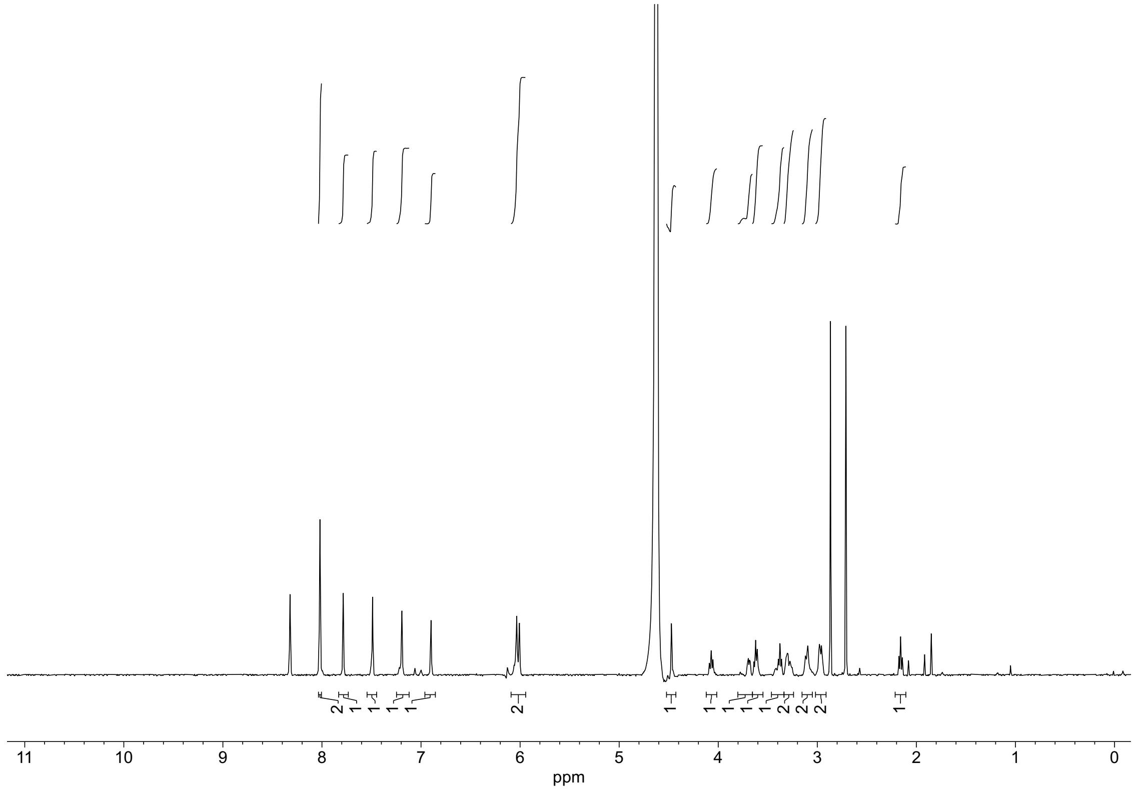


**3,5-Di(4-carbonoylpiperidine-N-carbonyl)phenyl)(6-nitrobenzo[*d*][1,3]dioxol-5-yl)methyl saxitoxin N21-ethylcarbamate (13)**

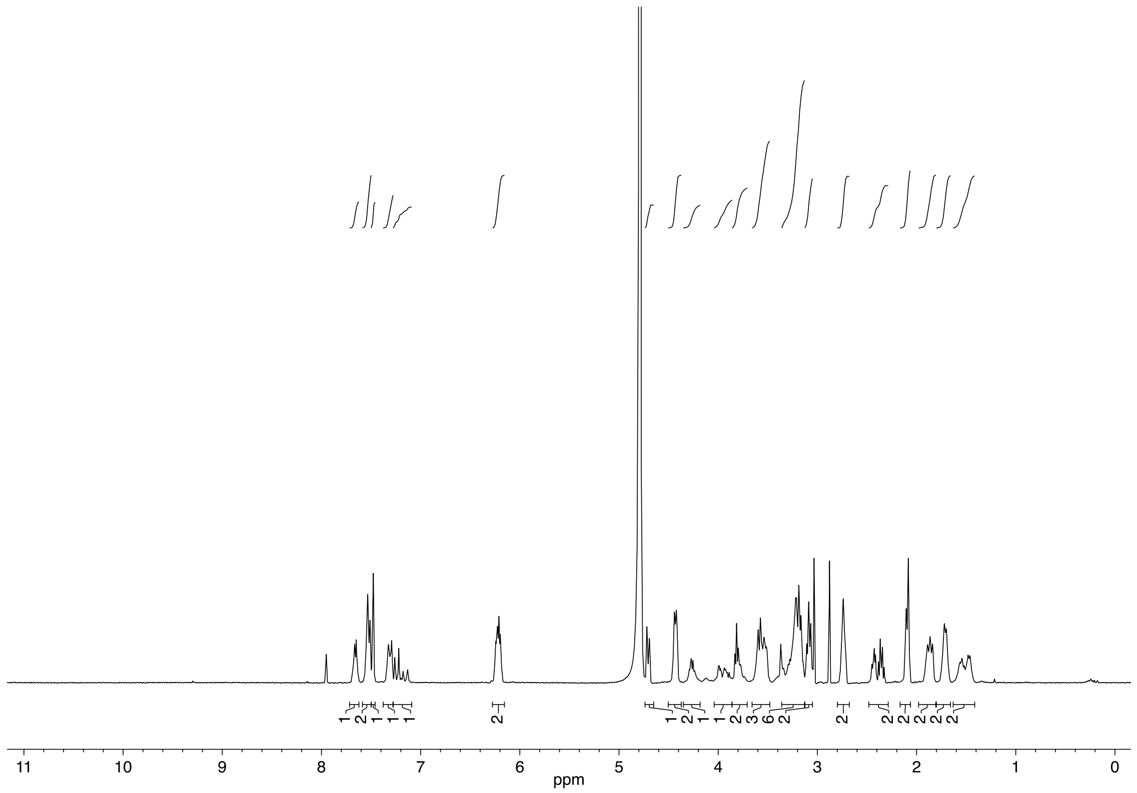
